# Metabolic adaptations and aquifer connectivity underpin the high productivity rates in the relict subsurface water

**DOI:** 10.1101/2023.06.25.546375

**Authors:** Betzabe Atencio, Eyal Geisler, Maxim Rubin-Blum, Edo Bar-Zeev, Eilon M. Adar, Roi Ram, Zeev Ronen

## Abstract

**Background:** Diverse microbes catalyze biogeochemical cycles in the terrestrial subsurface, yet the corresponding ecophysiology was only estimated in a limited number of subterrestrial, often shallow aquifers. Here, we detrained the productivity, diversity, and functions of active microbial communities in the Judea Group carbonate and the underlying deep (up to 1.5 km below ground) Kurnub Group Nubian sandstone aquifers. These pristine oligotrophic aquifers, recharged more than tens to hundreds of thousands years ago, contain fresh/brackish, hypoxic/anoxic, often hot (up to 60°C) water and serve as habitats for key microbial producers.

**Results:** We show that recent groundwater recharge, inorganic carbon and ammonium strongly influence chemosynthetic primary productivity in carbonate and sandstone aquifers (4.4-21.9 µg C d^-1^ L^-1^ and 1.2-2.7 µg C d^-1^ L^-1^, respectively). These high values indicate the possibility that the global aquifer productivity rates may be underestimated. Metagenome analysis revealed the prevalence of chemoautotrophic pathways, particularly the Calvin-Benson-Bassham cycle. The key chemosynthetic lineages in the carbonate aquifer were Halothiobacillales, whereas Burkholderiales and Rhizobiales occupied the sandstone aquifer. Most chemosynthetic microbes may oxidize sulfur compounds or ammonium, using oxygen or oxidized nitrogen as electron acceptors. Abundant sulfate reducers in the anoxic deeper aquifer have the potential to catabolize various organics, fix carbon via the Wood Ljungdahl pathway, and often possess nitrogenase, indicating diazotrophic capabilities. Our data suggest that connectivity between the aquifers and their exposure to energy inputs and surface water may play a key role in shaping these communities, altering physicochemical parameters and selecting taxa and functions. We highlight the metabolic versatility in the deep subsurface that underpins their efficient harnessing of carbon and energy from different sources.

## Introduction

Deep terrestrial subsurface habitats, including groundwater ecosystems, host ∼15% of the total biomass in the biosphere [1]. It mainly comprises bacteria and archaea, including novel lineages that play an essential role in biogeochemical cycling [1–4]. These microorganisms can thrive under extreme environmental conditions such as high temperatures and salinities, based on limited and diverse energy resources [5,6]. Even under these challenging conditions, diverse microorganisms thrive and are involved in biogeochemical transformations [5,7–12]. Yet, microbial lineages’ abundance, productivity and functionality, and relationship with their environment within such groundwater have been sparsely studied [13].

Environmental parameters shape the groundwater microbiome, as the availability of electron donors and acceptors driving microbial activity in groundwater depends on exposure to dissolved minerals, hydrochemical constituents, and the heterogeneity of the water-bearing formations [14,15]. Whereas these local conditions affect microorganisms’ spatial distribution and activity, the complex structure and dynamics of groundwater ecosystems may play an additional role in selecting microbial communities [16–18]. For instance, fractures and faults prevalent in the Earth’s crust may serve as hydrologic conduits or barriers for the dispersal and migration of nutrients among distinct water-bearing formations, thereby enabling or preventing microbial life [19]. Thus, microbial community dynamics are likely controlled by local terrestrial environmental parameters and geological processes, which are still poorly understood due to the challenges of accessing these deep, complex environments [19]. One of the fundamental questions concerns the mechanisms of energy and carbon acquisition by chemoautotrophic microbes that assimilate inorganic carbon in the dark underground, using multiple metabolic routes and electron donors [20–22]. These microbes have the potential to thrive in such habitats, including old groundwater, where even trace amounts of microbially-produced oxygen can support aerobic metabolism and boost productivity [23]. However, how the soluble and particular organics from the groundwater primary producers are reworked by the heterotrophic secondary producers is still not fully understood [22].

To date, most studies of groundwater microbes and their activity have focused on shallow aquifers [9,10,24–26], whereas our knowledge of deep aquifers with pristine ancient groundwater is limited [23,27,28]. This study aimed to address this knowledge gap by investigating microbial communities in deep (up to 1.5 km), mostly confined aquifers with highly pressurized ancient groundwater. Specifically, we focused on two deep regional aquifers from the Israeli Negev Desert, including the Kurnub Group Nubian sandstone and the overlying Judea Group carbonate aquifers, hereafter referred to as the sandstone and carbonate aquifers, respectively. These potentially interconnected aquifers have distinct histories and lithologies (See extended review provided in methods and supplementary information), providing an opportunity to study the ecophysiology of microbes in the deep, pristine, ancient groundwater, where interconnectivity between the aquifers may play a role in shaping the intrinsic communities. The access to a series of production wells containing fresh to brackish groundwater, spanning from hypoxic to anoxic conditions, facilitated the investigation of these deep microbial communities.

We asked if: i) the availability of electron donors and acceptors, as well as organic load, can alter microbial productivity rates and mechanisms of carbon fixation along the flow paths in the carbonate and underlying sandstone aquifers, ii) the diversity and function of microbial communities differ between the carbonate and sandstone aquifers, due to discrepancies in hydrogeological features, hydro-physiochemistry and various recharge sources, and iii) hydraulic interconnectivity between the aquifers may lead to the connectivity of microbial populations in specific sections along the flow paths.

To address these questions, we interrogated the long-term hydrogeochemical data of a series of production wells, assessed primary (PP) and secondary productivity (SP) (assimilation of H^14^CO_3_^-^ and ^3^H-leucine, respectively) and used metagenomics to explore the diversity and metabolic potential of microbes along the groundwater flow path in the Negev of these potentially interconnected aquifers. By employing a multi-faceted analysis encompassing various data types, across multiple sampling sites, we aim to advance our understanding of microbial communities within deep pristine aquifers containing ancient water mixed with younger water in certain sections. Using the productivity estimates in this study, we aim to update the rates of carbon sequestration in global aquifer systems.

## Results and Discussion

### Exceptional productivity rates in the ancient aquifers

We measured high primary productivity rates, in particular in groundwater within the carbonate aquifer. PP was significantly higher in the overlying carbonate aquifer (C wells) compared to the deeper sandstone (S wells), with mean values of 13.1±6.7 and 2.2±0.6 µg C L^-1^ d^-1^, respectively (MWT, p=0.03, **Table 1**). While the cell abundance (CA) had considerable variability (8.7 x10^6^ to 4.3 x10^8^ cells L^-1^, **Table 1**), productivity rates per cell in the carbonates aquifer were 0.05-0.73 pg of C cell^-1^d^-1^, as opposed to 0.10-0.31 pg C cell^-1^d^-1^ in the sandstone aquifer. Previous studies reported experimental carbon fixation rates varied from 0.043 ± 0.01 to 0.23 ± 0.10 µg C L^-1^ d^-1^ in shallow subsurface, and 0.0095 to 0.0560 µg C L^-1^ d^-1^ in southeastern Sweden’s deep crystalline bedrock [9,27]. Our estimates exceed these by orders of magnitude, suggesting the possibility that global groundwater productivity may be underestimated.

**Table 1.** Primary productivity (PP) and secondary productivity (SP) rates, cell abundance (CA) and specific primary productivity per well.

| ID | PP±stdv | SP±stdv | CA | Specific PP |
| --- | --- | --- | --- | --- |
| | $\mu\text{g C d}^{-1} \text{ L}^{-1}$ | $\mu\text{g C d}^{-1} \text{ L}^{-1}$ | cells $\text{L}^{-1}$ | $\text{pg C cell}^{-1} \text{ d}^{-1}$ |
| Carbonate aquifer |  |  |  |  |
| C1 | n.a. | 0.06±0.08 | 3.9 x10<sup>7</sup> | n.a. |
| C2 | 16.8±10.9 | 0.005±0.003 | 1.3x10<sup>8</sup> | 0.13 |
| C3 | 4.4 | 0.006±0.01 | 6.2x10<sup>7</sup> | 0.07 |
| C4 | 21.9 | <0.001 | 4.3x10<sup>8</sup> | 0.05 |
| C5 | 9.2±1.6 | 0.052±0.01 | 1.3x10<sup>7</sup> | 0.73 |
| Sandstone aquifer |  |  |  |  |
| S1 | 1.2±0.9 | 0.017±0.001 | 1.2x10<sup>7</sup> | 0.10 |
| S2 | 2.7±1.1 | 0.017±0.007 | 8.7x10<sup>6</sup> | 0.31 |
| S3 | 2.3±0.4 | 0.072±0.001 | 1.5x10<sup>7</sup> | 0.15 |
| S5 | 2.5±2.1 | 0.005±0.001 | 1.9x10<sup>7</sup> | 0.13 |

n.a.= No measurements available.

Given that 2.26 million km^3^ of global groundwater is stored in carbonate aquifers [9] and the average carbon fixation rates of 13.1±6.7 µg C L^-1^ d^-1^ observed in our carbonate wells, the high maximum estimates would reach a total of 10.8±5.6 Pg fixed annually in these ecosystems, representing 10 % of the global net primary production in marine and terrestrial ecosystems (104.9 Pg C/year, [32]). However, it is crucial to note that during the PP incubations we did not account for the influence of confining pressures experienced down the boreholes, which can reach up to 126 bars at depths exceeding 1 km below the surface (**Fig. 1c**). Thus, the influence of such pressures on primary production in deep aquifers remains elusive, highlighting the need for future investigations.

**Figure 1.**
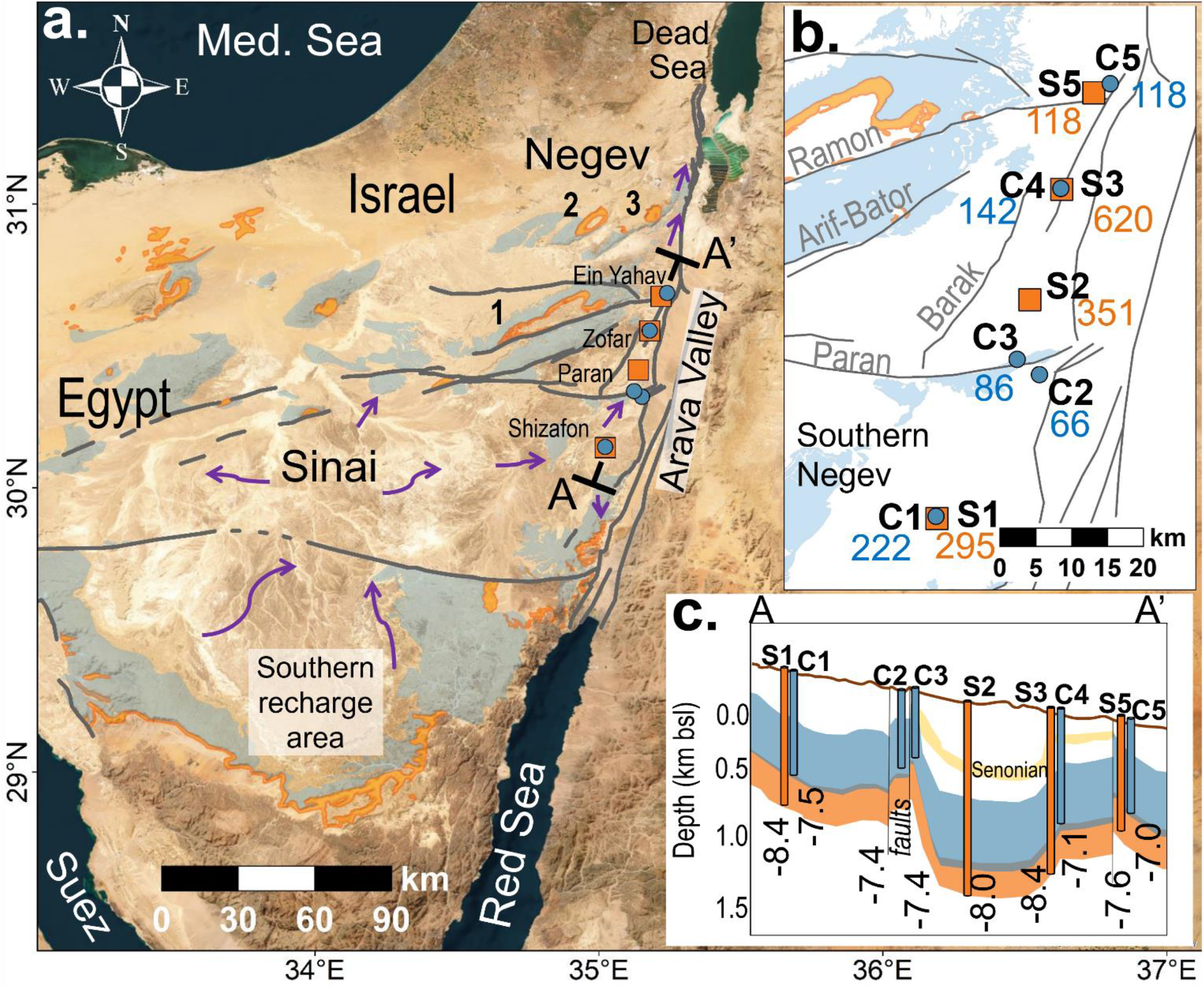
Location of wells tapping the sandstone and the overlying carbonate aquifers (S and C wells, respectively) along the eastern groundwater flow path (purple arrows), stretches from the southern central Sinai, through the Arava Valley, and towards the outlet south of the Dead Sea. Also shown are sandstone and carbonate outcrops (orange and light blue shades, respectively), and major E-W faults over the Negev Desert (gray lines and labels). S wells are denoted by orange squares, while C wells are represented by and blue circles. Numbers in subfigure (a) indicate some of the major anticlines in the studied area, including (1) Ramon, (2) Hatzera, and (3) Hatira. (b) ^81^Kr ages of the sandstone and carbonate aquifer wells (orange and blue labels, respectively) follow previous studies [29–31], and are reported in thousands of years (kyr). (c) A schematic cross-section A-A’ of the nine sampled wells along the eastern flow path (not to scale). Numbers annotate δ^18^O values (‰). Satellite imagery source: Esri, Maxar, GeoEye, Earthstar Geographics, CNES/Airbus DS, USDA, USGS, AeroGRID, IGN, and the GIS User Community.

### Primary productivity may be driven by aquifer recharge, carbonate rock weathering, and nitrogen cycling

Spearman correlation analysis indicated that PP rates were significantly correlated to δ^18^O, bicarbonate (HCO_3_^-^), and ammonium (NH_4_^+^) (See **Supplementary Fig. S1**). Thus, the high productivity in the carbonate aquifer is potentially due to the synergistic effect of several factors, including freshwater recharge and carbonate rock weathering.

The significant positive correlation between the PP rates and δ^18^O values supports the contribution of a local recharge component to primary productivity (**Table 2**, **Fig. 1c**). In general, samples with low δ^18^O values are ^81^Kr-depleted (**Table 2**), suggesting that they originated from older precipitation, while samples with higher δ^18^O show higher ^81^Kr activity, and are likely affected by more recent recharge events [30,31]. Since recharge water flows downward, microorganisms consume oxygen and nutrients over time [33]. Hence, older groundwater with low δ^18^O values, such as those in the deep sandstone aquifer, might provide less favorable conditions for autotrophic growth than groundwater with higher δ^18^O values, possibly leading to the use of alternative electron acceptors other than oxygen, and, thus lower PP rates.

**Table 2.** Hydrogeochemical and physical characteristics of the sampled wells in molar concentrations.

| ID | Well name | Age<sup>a</sup> | Median screen depth | <sup>81</sup>Kr<sup>a</sup> | DO | EC | T | HCO<sub>3</sub><sup>-1</sup> | SO<sub>4</sub><sup>-2</sup> | HS<sup>- b</sup> | TOC | NH<sub>4</sub><sup>+</sup> | DIC | δ<sup>18</sup>O<sup>a</sup> |
| --- | --- | --- | --- | --- | --- | --- | --- | --- | --- | --- | --- | --- | --- | --- |
|  |  | kyr | m below ground level (mbgl) | R/Ra | μmol L<sup>-1</sup> | mS cm<sup>-1</sup> | °C | mmol L<sup>-1</sup> | mmol L<sup>-1</sup> | μmol L<sup>-1</sup> | mmol L<sup>-1</sup> | μmol L<sup>-1</sup> | mmol L<sup>-1</sup> | (‰) |
| Carbonate aquifer |  |  |  |  |  |  |  |  |  |  |  |  |  |  |
| C1 | Shizafon 11 | 222 | 666 | 0.51 | 3.1<sup>c</sup> | 3.2 | 50 | 5.80 | 10.00 | 4.0 | 0.08 | 233 | n.a. | -7.5 |
| C2 | Paran 129 | 86 | 580 | 0.76 | 2.8 | 4.1 | 37 | 5.50 | 14.30 | 15.2 | 0.05 | 209 | 8.9 | -7.4 |
| C3 | Paran 21 | 66 | 718 | 0.82 | 4.7 | 2.8 | 39 | 5.70 | 5.80 | 48.1 | 0.07 | 85 | 8.8 | -7.4 |
| C4 | Zofar 220 | 142 | 471 | 0.65 | 2.8 | 2.5 | 41 | 5.60 | 5.80 | 287.2 | 0.07 | 261 | 7.6 | -7.1 |
| C5 | Ein Yahav 7 | 118 | 635 | 0.70 | 4.7 | 3.2 | 39 | 4.70 | 5.90 | 74.5 | 0.09 | 122 | 5.2 | -7.0 |
| Sandstone aquifer |  |  |  |  |  |  |  |  |  |  |  |  |  |  |
| S1 | Shizafon 1 | 295 | 854 | 0.41 | 0.9 | 3.6 | 53 | 3.4 | 10.2 | 9.5 | 0.08 | 101 | 5.0 | -8.4 |
| S2 | Paran 20 | 350 | 1415 | 0.35 | 1.3 | 3.4 | 60 | 3.3 | 6.4 | 25.4 | 0.08 | 112 | 4.3 | -8.0 |
| S3 | Zofar 20 | 620 | 920 | 0.15 | 1.3 | 10.2 | 51 | 4.2 | 8.6 | 43.4 | 0.03 | 146 | 3.5 | -8.4 |
| S5 | Ein Yahav 6 | 118 | 822 | 0.70 | 2.2 | 3.1 | 44 | 5.2 | 5.8 | 92.0 | 0.08 | 59 | 5.6 | -7.6 |

a. From [30,31,46].
b. From Mekorot National Water Company and the Israeli Water Authority database. Unpublished data.
c. Well log from the Geological Survey of Israel-1998. Unpublished data.

Dissolved oxygen (DO) can enhance primary productivity in various ecosystems given the high energy yields of reactions that use oxygen as a terminal electron acceptor (e.g., [23,34,35]). The recharge component of fresh water, facilitated by overland flows and floods from nearby outcrops and desert streams, respectively, can introduce DO into the overlying carbonate aquifer [36,37]. In addition, microbial chlorite (ClO_2_) dismutation in groundwater could release trace oxygen levels, potentially fueling chemosynthesis [23]. We found genes for chlorite reduction (*cld*) in 16 MAGs, such as those of *Blastomonas*, *Bradyrhizobium*, *Deinococcus*, JAAYVI01 (Elusimicrobiota), and the archaeal *Nitrosotenuis*, some of which were prominent in wells S1, S5, S2, and C2. This hints at the possibility of dark oxygen-driven productivity in the Negev aquifers. We note that both the carbonate and sandstone aquifers have low steady-state DO with values below 6 µmol L^-1^ (0.2 mg L^-1^) (**Table 2**), whereas DO sources and respiration rates in these aquifers remain to be estimated.

Furthermore, inorganic carbon availability may determine primary production rates [38,39]. The major source of dissolved inorganic carbon (DIC) in the aquifers is likely from carbonate rock weathering, a major source of HCO_3_^-^ [40]. We observed high concentrations of bicarbonate (5.5±0.4 and 4.0±0.8 mmol L^-1^ in carbonate and sandstone aquifers, respectively, MWT, *p*=0.032, **Table 2**), suggesting that ample inorganic carbon is available for autotrophs, in particular, in the carbonate aquifer. Other studies have shown that most sandstone wells near the Arava Rift geological fault line (**Fig. 1**) contained a high contribution of mantle-derived gases, such as CO_2_ [41,42]. These results are in line with previous observations stressing that inorganic carbon is a significant factor influencing the community structure of CO_2_-fixing bacteria in karst wetland groundwater [43].

Lastly, we observed a positive relationship between the PP and NH_4_^+^ concentrations. On one hand, NH_4_^+^ could accumulate due to biomass degradation. Alternatively, dissimilatory nitrate reduction to ammonia (DNRA) can contribute to overall productivity under hypoxic conditions, as observed in a karst aquifer [44]. In support of this assumption, we identified 44 MAGs that encoded the NrfA nitrite reductase often used in DNRA, especially prominent among anaerobes such as Chloroflexota and Acidobacteriota (**Fig. 2**). Ammonium, in turn, can serve as an electron donor and fuel the primary production of nitrifying bacteria and archaea [34,45].

**Figure 2:**
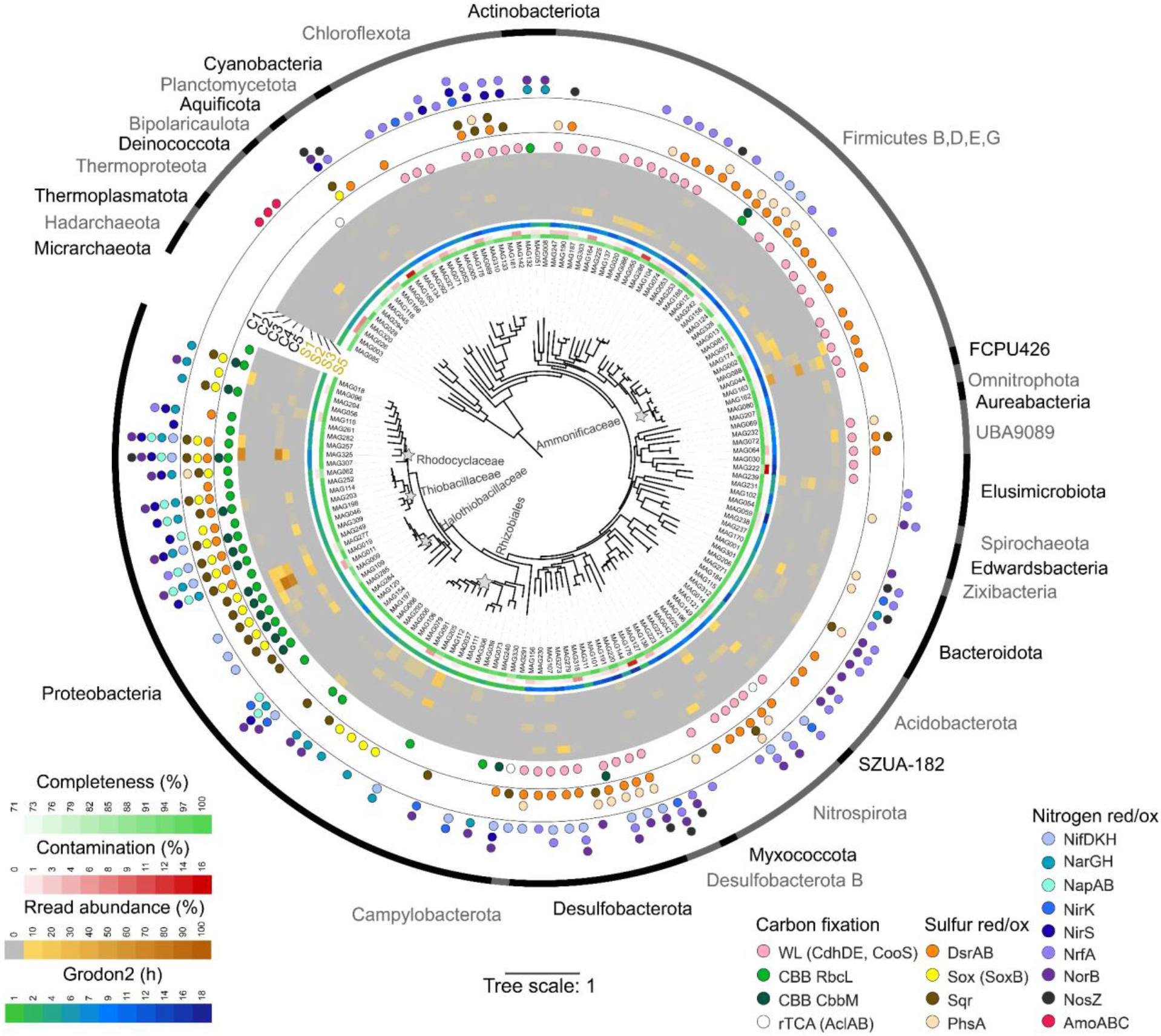
Phylogeny, diversity and key functions of bacteria and archaea in sandstone and carbonate aquifers. The iTOL representation is based on the GTDB-tk marker gene set and treeing. Key metabolic pathways are indicated in different colors. A heatmap shows the median read coverage of each MAG per sample site. MAGs representing the key lineages at the family level are highlighted in bold. Three MAGs were excluded from the tree due to insufficient marker gene hits (27 - Riflebacteria, 189 – Desulfotomaculaceae and 217 – Rhizobiales).

### Aquifer microbes have the metabolic potential to fix carbon

We reconstructed 140 bacterial and 8 archaeal high-quality metagenome-assembled genomes, spanning 32 phyla (MAGs, CheckM 0.94±0.05 completeness, and 0.02±0.03 contamination, **Fig. 2**). Archaea were usually scarce (0 to 6% read abundance), following previous studies which reported similar patterns in groundwater [19,47–49]. The key pattern that we observed was the predominance of Halothiobacillales 28-57-27 lineage in the overlain carbonate aquifer (**Fig. 2**). The representative cultivar in this clade is *Halothiobacillus neapolitanus,* a sulfur-oxidizing chemolithoautotrophic bacterium in which carboxysomes were studied in detail [50,51]. The southmost Shizafon site (wells C1 and S1) stood out, as the sulfur-oxidizing Rhodocyclaceae UBA2250 sp004323595 lineage (Burkholderiales, corresponds to the GCA_004323595.1 NCBI submission, identified previously as *Sulfuricystis*, [52]) was predominant in both carbonate and sandstone aquifers. Moreover, the ammonia-oxidizing *Nitrosotenuis* was prominent in S1 (circa 6% read abundance). Other sandstone stations were mainly inhabited by Ammonifexales, Thermodesulfovibrionales, Rhizobiales, and Desulfotomaculales (see alpha and beta diversity analyses in **Supplementary Note 5, Supplementary Figs. S2 and S3**).

Based on taxonomic inference, we hypothesized key lineages had the metabolic potential for autotrophy. Calvin-Benson-Bassham (CBB) cycle accounted for at least 20% of the read abundance across all the samples, suggesting that chemosynthesis is a key process in these aquifers (**Fig. 3a**). 32 MAGs encoded the ribulose-1,5-bisphosphate carboxylase/oxygenase (RubisCO, form I -16, from II – 2, 14 both). Both forms were encoded by the most abundant MAGs, representing the populations of Rhodocyclaceae UBA2250 sp004323595 (MAG325), Halothiobacillaceae 28-57-27 (e.g., MAGs 9,11, 19, 109, 277 and 285) and *Thiobacillus* (MAG114). These RuBisCo forms differ in their affinity to oxygen and inorganic carbon, as form I enzymes function better under high O_2_ and low CO_2_ conditions, and form II is the opposite [53–55]. Thus, the key autotrophs may be adapted to fluctuating concentrations of oxygen or CO_2_. These bacteria also encoded carbonic anhydrases, suggesting the ability to maintain homeostasis of their carbonate system. The abundance of Halothiobacillaceae was strongly correlated with the average PP rates (Spearman correlation, rho=0.90; p=0.005). Also, their less than 5 h maximum growth rate estimates exceeded those of most other autotrophs (e.g., Burkholderiales MAG325 - 8.2 h, Ammonifexales MAG163-11.6 h, Thermodesulfovibrionales MAG42 -10.7 h, **Fig. 2**). Thus, the Halothiobacillaceae autotrophy likely maintains the exceptional activity of carbonate wells.

**Figure 3.**
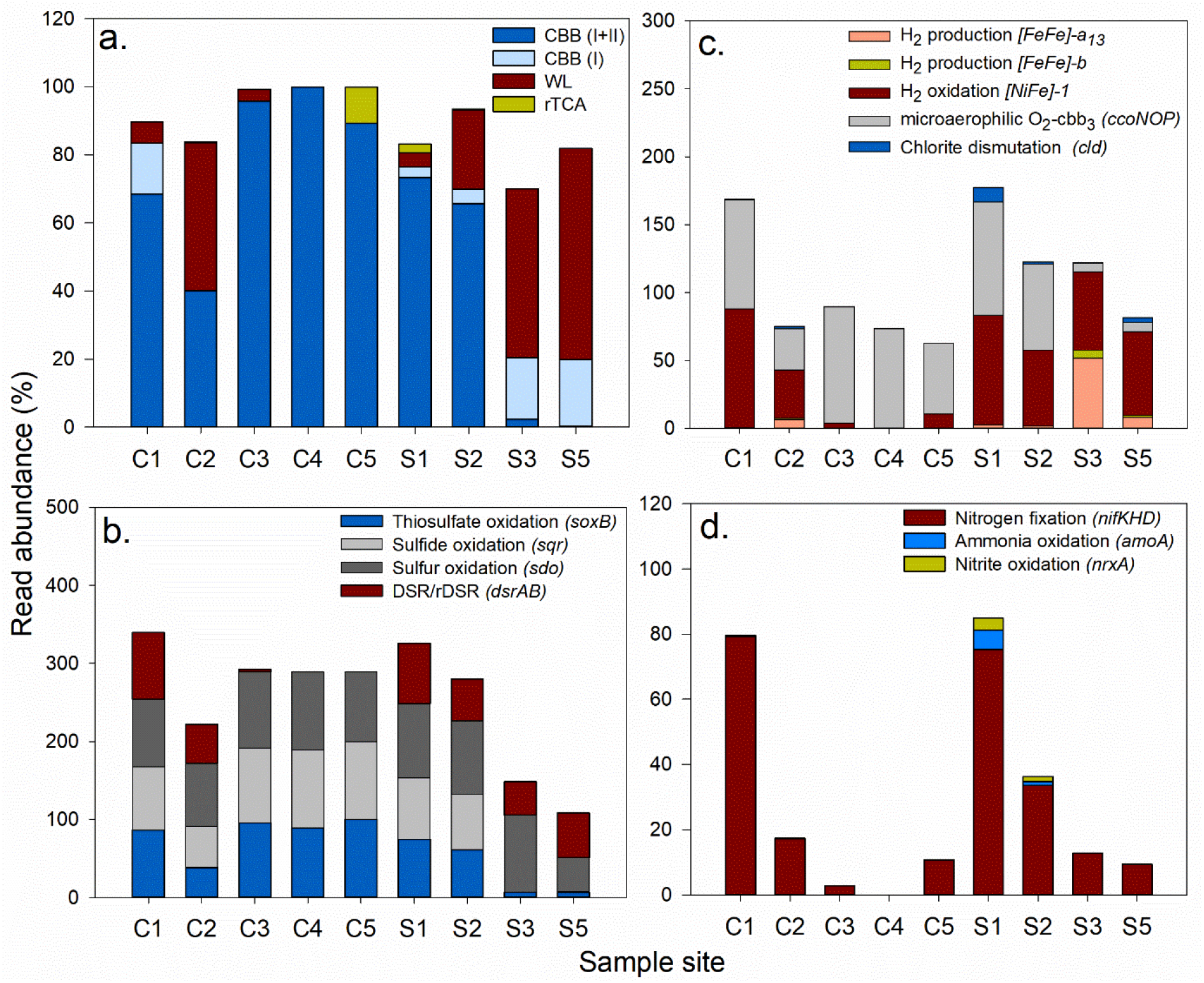
Key metabolic features of aquifer microbes. The cumulative read abundance of MAGs that encode: a) carbon fixation b) dissimilatory sulfur cycling; c) hydrogenases, microaerophilic oxygen metabolism-cytochrome c oxidase cbb_3_, and chlorite dismutation genes; d). nitrogen fixation and nitrification; CBB – Calvin-Benson-Basham cycle, rTCA – reductive Krebs cycle; WL-Wood Ljungdahl pathway. DSR/rDSR-dissimilatory sulfur metabolism.

We note that type I RuBisCo was also encoded by *Methylovirgula*, a clade of facultative methylotrophs [56], which can employ the CBB cycle to assimilate excess CO_2_, using energy from formaldehyde oxidation [57]. We found a complete TCA cycle in this MAG, suggesting that methylotrophic autotrophy is not obligate. *Methylovirgula* (MAG112) was prominent in S3 and S5 wells, suggesting that small organic molecules can fuel primary productivity in the sandstone aquifer. However, methanotrophs were scarce, as particulate methane monooxygenases were found only in two rare lineages from S2 well (*Fontimonas* MAG197 and Binatia MAG176, both below 1% read abundance), and no *mcr* genes, indicative of anaerobic methane oxidizers, were found.

Additional carbon fixation pathways were attributed to less abundant microbes. A limited number of MAGs encoded the rTCA cycle, in particular, the AclAB subunits of the ATP-citrate lyase (Nitrospiraceae MAG149 and *Sulfuricurvum* MAG291, but also the less studied taxa such as *Bipolaricaulis* MAG166 and Ozemobacteraceae MAG27). Among these lineages, *Sulfuricurvum* was abundant in carbonate aquifer C5 (circa 10% read abundance), where DIC was the lowest, and notable influence from younger recharge events (**Table 2**). The archaeal MAG contained *amo* genes needed for nitrification and encoded the key enzymes of the autotrophic 3-Hydroxypropionate/4-Hydroxybutyrate (3-HP/4-HB) cycle (4-hydroxybutyryl-CoA dehydratase and 3-hydroxybutyryl-CoA dehydratase, Berg *et al.*, 2007), underpinning its ability to fix carbon based on nitrification (**Supplementary Table S5**). The considerable abundance of *Nitrosotenuis* in S1 (MAG28, 5.8% read abundance) and Nitrospiraceae (MAG149, 2.6% read abundance) indicate that archaeal and bacterial nitrification can contribute to carbon fixation in this well, alongside the sulfur-oxidizing Rhodocyclaceae.

The abundance of microbes encoding the energy-demanding CBB cycle markedly decreased in well C2 in the carbonate aquifer, and following the flow direction downstream the sandstone aquifer from well S2 towards S3 and S5 (**Fig. 3a**) [59,60]. In these wells, microbial communities appear to use the efficient Wood Ljungdhal pathway (WL) for carbon fixation [61]. We found all the three key *cdhD*, *cdhE* and *cooS* genes of the Wood Ljungdahl pathway in 50 MAGs, spanning the Actinobacteriota, Chloroflexota, Desulfobacterota, Firmicutes, Nitrospirota, SZUA-182, UBA9089 and Thermoplasmatota phyla (**Fig. 2, 3a** and **Supplementary Fig. S3**). Most of these phyla comprise the anaerobic sulfate reducers and acetogens which were abundant in the sandstone aquifer, and one well from the carbonate aquifer (C2). However, sulfate reducers and other acetoclastic microbes use this route not only to fix CO_2_ but also for energy conservation [62,63]. Thus, we could not rule out the possibility that some organisms encoding for this pathway may not be fixing carbon [64].

We note that oxidation of organics, fermentation and acetate oxidation are the most characteristic traits of the aquifer microbiome (**Fig. 4)**. Therefore, alongside the specialist producers, we find a diverse heterotrophic/mixotrophic community that benefits from reworking the necromass and exudates of the primary producers. These organisms drive secondary productivity, which is considerably smaller than PP (PP/SP = 3.2×10^1^ – 6.0×10^4^ see **Supplementary Note 4**). This may indicate the slow heterotrophic metabolism in a hypoxic to anoxic environment but also hints at the caveats in the leucine uptake assay. We note that similar proportions between PP and SP were observed in chemosynthetic mats associated with gas seepage in the deep sea (Rubin-Blum et al., in prep.) and in the polar oceans [65].

**Figure 4.**
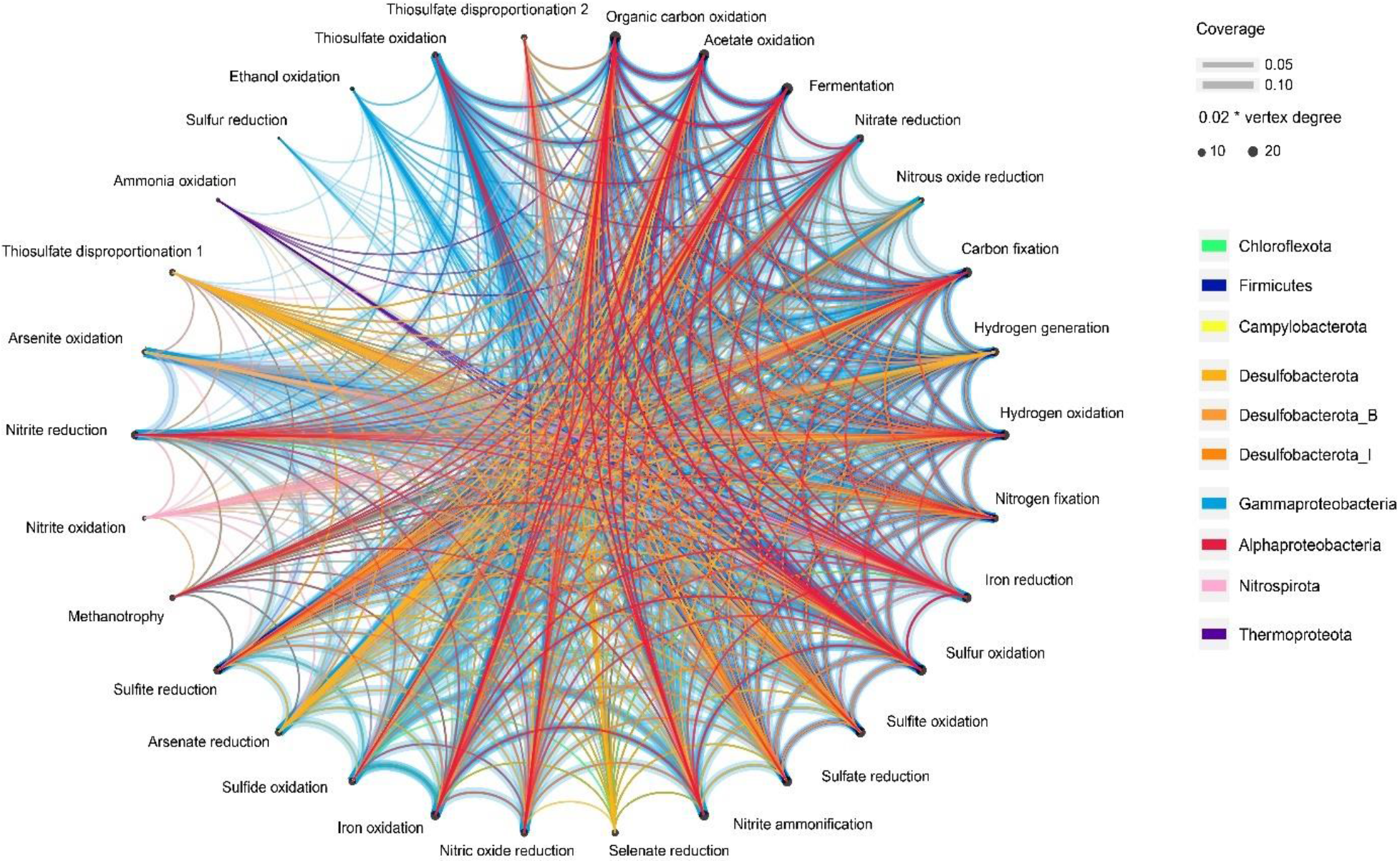
Taxonomic affiliation of metabolic functions in the aquifer community. Key lineages are highlighted in colors, the others are shown in the background (grey). Vertex sizes correspond to the number of connections. Taxa are presented at the phylum level (only Gammaproteobacteria and Alphaproteobacteria are at the Class level, to display the marked differences within Proteobacteria).

### Dissimilatory sulfur, hydrogen and nitrogen turnover provide electron donors and acceptors

Our data suggest an array of sulfur, hydrogen and nitrogen transformations in the studied aquifers (**Supplementary Fig S4**). We identified 22 MAGs encoding the Sox Kelly-Friedrich pathway for thiosulfate/sulfide oxidation in both the sandstone and carbonate aquifers (mostly the chemosynthetic taxa Halothiobacillales, Burkholderiales, Rhizobiales and Campylobacterales, **Figs. 2 and 3b**). We identified 57 MAGs containing the dissimilatory sulfate reduction (DSR) and/or the reverse rDSR (encoded by the *dsrAB*) (**Figs. 2 and 3b**). Yet, the DSR/rDSR were most common in the sandstone aquifer (circa 55%), but rare in some wells (C3 to C5) from the overlain carbonate aquifer (circa 0.01-3%). Sulfate reduction is likely the prevailing anaerobic respiration pathway in both aquifers, following the high sulfate and sulfide concentrations reaching up to 14.3 mmol L^-1^ and 0.29 mmol L^-1^, respectively (**Table 2**). Some key lineages can disproportionate thiosulfate (**Fig. 2 and Supplementary Table S3**). Following the previous studies in groundwater [66], our results highlight the potential use of hydrogen as an energy source, potentially driving carbon fixation [21,22,67–69]. Among the 57 MAGs containing *dsrAB*, we identified 43 MAGs encoding for [NiFe]- group I hydrogen uptake hydrogenase (**Supplementary Table S6**), which can couple hydrogen oxidation to sulfate respiration [39]. Furthermore, 23 MAGs encoded the WL pathway (of 43). For instance, Nitrospirota MAG 42, Firmicutes MAG 74 and Desulfobacterota MAG 220 contained the *hndB* genes (EC 1.12.1.3) predicted to be involved in anaerobic hydrogen oxidation coupled with sulfate reduction, thereby potentially producing energy and fixing carbon via the WL pathway (**Supplementary Table S7**) [70]. In addition, we observed the presence of [FeFe]-hydrogenases, which are typically linked to the production of H_2_ during fermentation (**Fig. 3c**)[23,71]. This was evident in Firmicutes MAG 13, 20 and 303, common in well S3. The occurrence of various hydrogenases (**Fig. 3c and Supplementary Fig. S3**) suggests that hydrogen may fuel microbial activity in the Negev Desert aquifers, particularly the deeper sandstone aquifer, yet concentrations and sources of hydrogen remain to be determined.

Nitrification is another putative driver of carbon fixation in the aquifers. Bacterial and archaeal ammonia oxidizers were present, primarily in carbonate wells C1 as well as in sandstone wells S1, and S2 (**Fig. 3d**). 6 MAGs encoded for nitrite oxidation to nitrate (*nrxA*), 2 of which could fix carbon via the rTCA and WL pathway (Nitrospiria MAG 149 and Thermodesulfovibrionia MAG 25). Given that nitrite concentrations were below the detection limits across all samples (<0.016 mmol L^-1^), there is likely a cryptic turnover of NOx. These observations suggest that nitrification can fuel productivity even in hypoxic to anoxic aquifers. Oxygen is a likely electron acceptor for the chemosynthetic microbes, as they encoded terminal oxidases, in particular the cbb_3_-type type (*ccoN*), adapted to low oxygen levels [72,73] (**Supplementary Fig. S3**). This adaptation may enable microaerophilic respiration (**Fig. 3c, 6**) [69].

In these oxygen-limited aquifers, denitrification appears to play an important role in microbial energy conservation. 24 MAGs encoded nitrate reduction to nitrite, including the prominent Burkholderiales (**Fig. 2 and Supplementary Fig. S4**). We could trace the complete denitrification to N_2_, mainly by gammaproteobacterial lineages, including the dominant Rhodocyclaceae UBA2250 MAG325, which encoded the Nap periplasmic nitrate reductase, the NirBD and NirS nitrite reductases, NorBC and NosZ. However, most microbes could only complete parts of the full route, for example, most Thiobacillaceae encoded all the steps, excluding the final reduction of N_2_O to N_2_, whereas some anaerobes encoded only the last step. Overall, 21 MAGs encoded nitrate respiration via the NarGHIJ respiratory nitrate reductase, 40 encoded NirBD, 9 NirK, 18 NirS, 34 NorB (17 NorC) and 10 NosZ. This indicates that aquifer communities can denitrify, either egoistically (mainly in wells S1, S2 and C1) or synergistically (**Supplementary Fig. S4**). Our findings agree with previous studies that reported the importance of the nitrogen cycle in groundwater with low levels of inorganic nitrogen [22,74–76].

### Genomic evidence for nitrogen fixation in pristine groundwater

We identified 26 MAGs encoding the key proteins of the nitrogenase complex (*nifD*, *nifK* and *nifH*), within diverse taxa, including the anaerobic sulfate reducers Desulfobacterota, Firmicutes (Desulfotomaculia) and Nitrospirota (Thermodesulfovibrionia), the poorly studied lineages such as SZUA-182, and the prominent chemosynthetic lineages, including the common Rhodocyclaceae and Thiobacillaceae (including the abundant UBA2250 MAG 325), various Halothiobacillaceae, Sulfurimonadaceae and Nevskiaceae (**Fig. 2 and 3d**). The high energy demand of nitrogen fixation may be achieved under the aquifer’s low DO levels, which is likely met by sulfur oxidation and reduction dynamics, using the trace oxygen or nitrate as an electron acceptor. However, the high ammonium concentrations (59 to 261 µmol L^-1^,**Table 2**) and low C:N (<1.5) indicate that nitrogen may not be a limiting nutrient in the pristine groundwater [77]. It has been hypothesized that fixed nitrogen may support subsurface biomass [78], yet nitrogen budgets and rates of N_2_ fixation in these systems remain to be elucidated.

### Microbial Composition Provides Clues to Aquifer Connectivity

Some adjacent wells had similar community structures, despite originating from different aquifers, either indicating aquifer connectivity and intermixing of microbial populations, or selection by similar physicochemical characteristics. The most striking example is the southern C1 and S1 wells with highly similar communities dominated by Rhodocyclaceae UBA2250 MAG325 (**Fig 5a**). This is despite belonging to different aquifer formations, with differences in the origin of recharge as suggested by their distinct ^81^Kr age (70 kyr gap, **Fig. 1b**) and water stable isotope composition (δ^18^O values of -7.5 and -8.4 ‰, respectively). The co-occurrence of the same genotypes indicates active hydraulic interconnection between these aquifers, as previously suggested [37,79].

**Figure 5.**
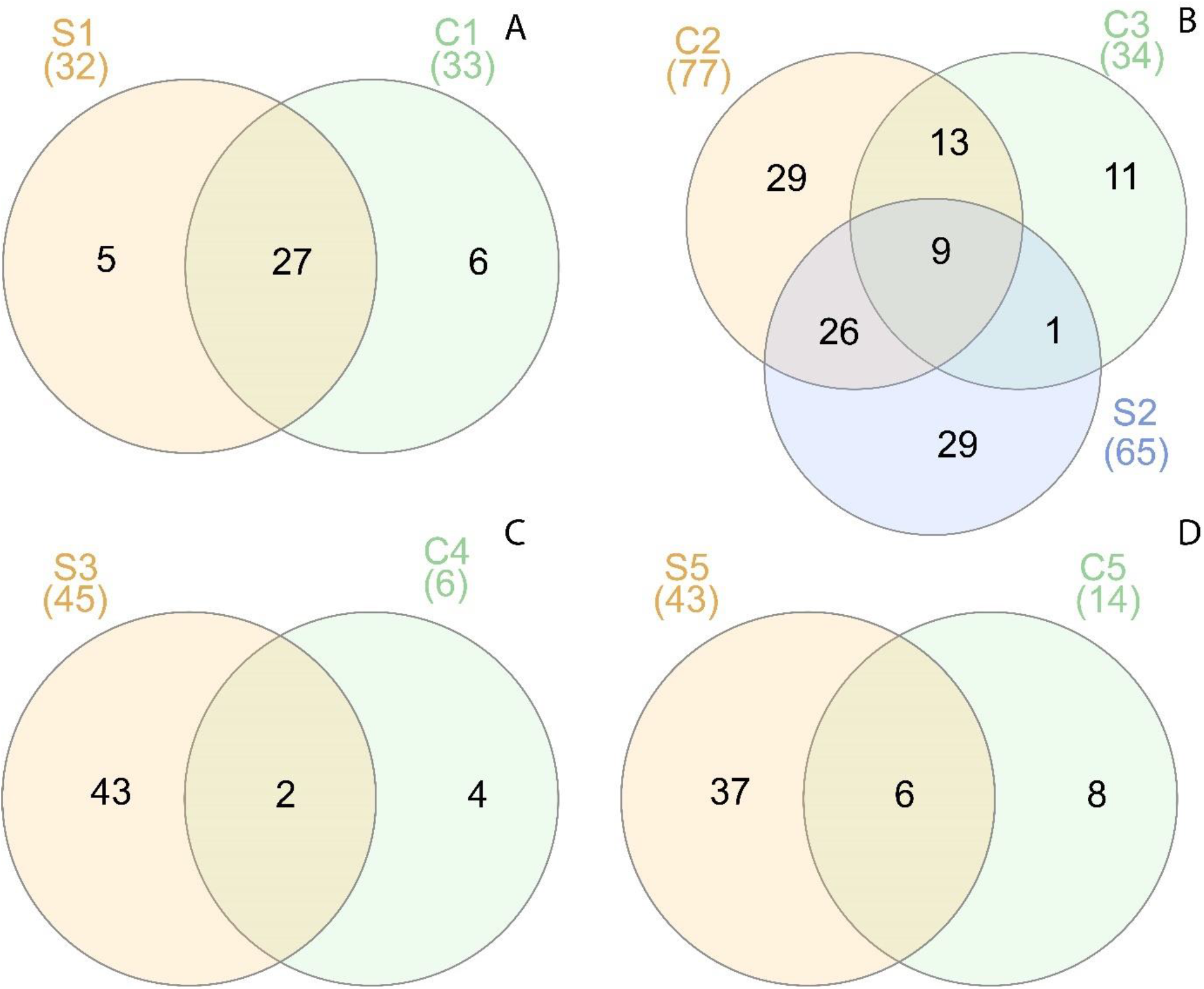
Co-occurrence of metagenome-assembled genomes in adjacent aquifers. A) The marked overlap of communities in the southern Shizafon C1 and S1 wells. B) Some taxa co-occur in Paran wells C2, C3 and S2. Fewer taxa co-occur in Ein Yahav (C) and Zofar (D) sandstone and carbonate wells.

The preferential groundwater flow paths along geological faults in the Negev Desert may enhance groundwater intrusion into the sandstone aquifer from neighboring and shallower aquifers. The Paran fault (**Fig. 1b**), for instance, may enable groundwater flow from the above carbonate formations into the underlying sandstone aquifer in the Paran basin, where well S2 is located. We found nine co-occurring MAGs between Paran S2 (sandstone), C2 and C3 (carbonate) wells, including the dominant Halothiobacillales MAGs (**Figs. 2 and 5b**). This hints that the composition of microbial communities in complex aquifer systems may be influenced not only by deterministic and selective environmental factors [15] but also by stochastic hydro-meteorological events [18,19].

This was not the case for the northern S3, C4 (Zofar, **Fig. 1**) and S5/C5 (Ein Yahav **Fig. 1**) wells as only a few MAGs co-occurred in the sandstone and carbonate aquifers, excluding the dominant taxa (**Figs. 2 and 5c,d**). In each formation, the continuity of the aquifers is indicated by a considerable overlap between the communities in S3-S5 and C4-C5 wells (**Fig. 2**). It was previously postulated that groundwater rejuvenation downstream the sandstone aquifer’s path, towards S5 serves as an indication of younger recharge, facilitated by the surroundings geological faults, that affect this part of the aquifer [30] (**Fig. 1ab**). Given that microbial populations don’t intermix between the aquifers, we speculate that either i) the groundwater in the sandstone aquifer rejuvenates from alternative, yet not identified sources of younger water (e.g. the nearby Senonian chalk formations), or ii) carbonate aquifer microbes are being selected against by the sandstone aquifer environment, or iii) physical or chemical barriers within the faults limit the exchange of populations.

## Conclusions

We still lack knowledge on how life is sustained in the large pristine aquifers. Our study highlighted the exceptionally high carbon fixation rates in an extensive carbonate aquifer, whereas autotrophic activity in the underlying sandstone aquifer is also substantial. Our measurements add a new high estimate of global productivity in groundwater. We identified multiple factors that can boost aquifer productivity, such as groundwater recharge and rock weathering yielding inorganic carbon. We emphasize the importance of local selection and aquifer interconnectivity, both between the aquifers and with the surface, as these factors may determine community composition and function, and therefore its ability to sequester carbon. The balance between electron donors and acceptors may play a key role in determining the productivity of these systems. The sources of these key molecules, such as oxygen and nitrate, sulfide, ammonium and methanol are not fully understood.

For example, previous analyses of sulfate and carbon isotopes in the deep sandstone aquifer suggested that bacterial sulfate reduction is a significant microbial process in the aquifer’s confined zone[80]. To date, sulfate reduction is considered a main source of sulfide in the carbonate aquifer of the northwestern Negev [81], ∼ 70 km northwest of the study area. The carbonate aquifer may receive water rich in organic matter from an overlying aquifer found within the Senonian rocks of the Mt. Scopus Group (**Fig. 1c**), yielding hydrogen sulfide in the carbonate layers due to microbial oxidation of organics [81]. Yet, we did not find prominent sulfate reducers in wells C3, C4, and C5 which had high sulfide levels (48.1, 74.5, and 287.2 µmol L^-^, respectively). Instead, these wells were dominated by Halothiobacillales chemoautotrophic sulfur oxidizers. Alternatively, organic matter is degraded before reaching the carbonate formation [82] and sulfidic water is introduced to the aquifer. To validate this hypothesis, further investigations, including analysis of sulfate isotopes across all wells in the Negev and Arava basins, are necessary.

We show that the metabolically versatile aquifer microbes can employ different strategies to efficiently harness the energy and acquire carbon in these habitats, where oxygen and organic carbon are depleted (**Fig. 6)**. The complex metabolic handoffs and other interactions in these communities are still not fully understood. Other aquifer microbiota, including eukaryotes and viruses, and their role in these systems remains to be explored.

**Figure 6.**
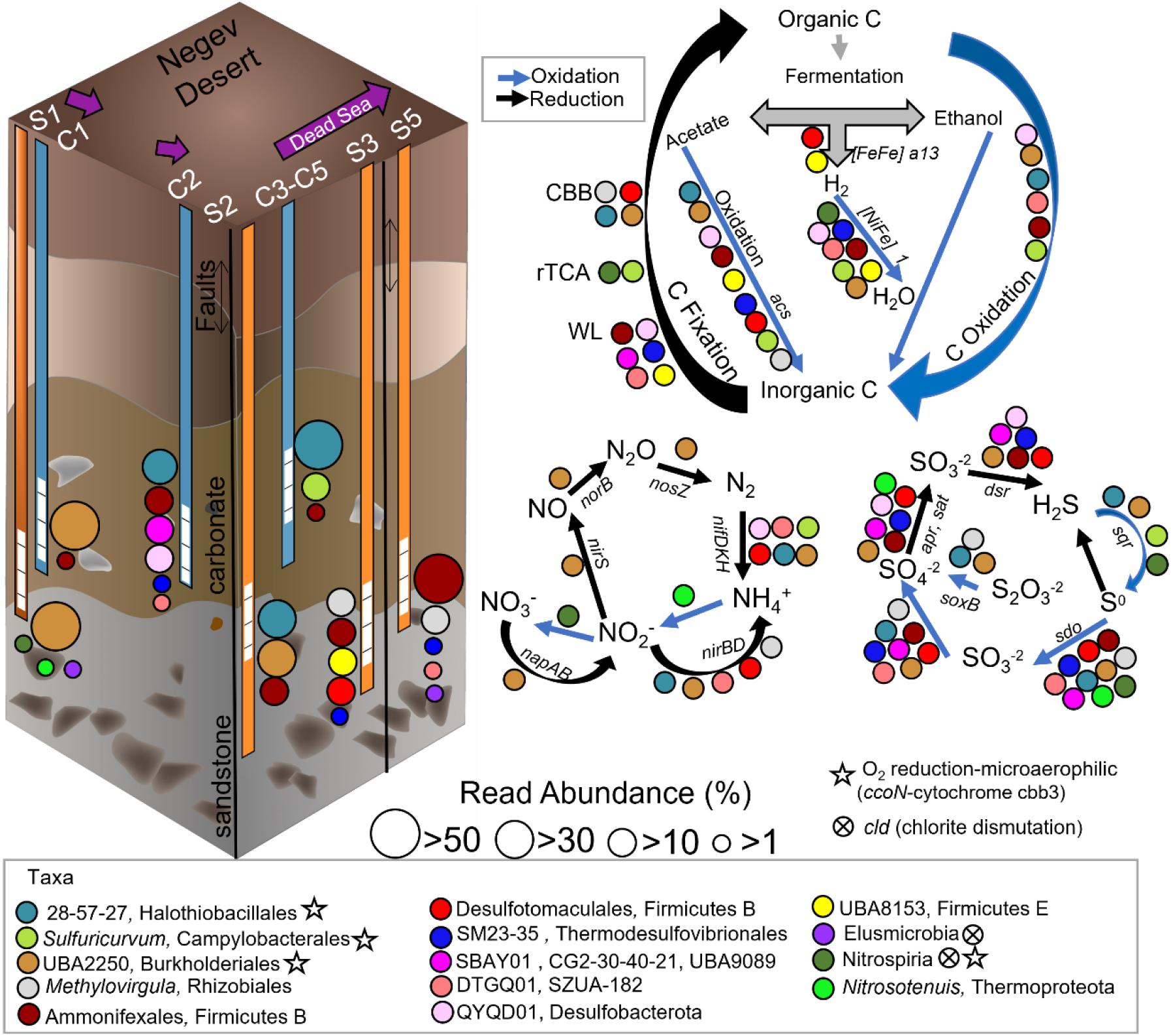
Metabolic reconstruction at a community scale along the eastern groundwater flow path of the two aquifers (purple arrows), including some of the most abundant lineages colored and grouped by their taxonomic identity. The figure summarizes biogeochemical cycling processes (carbon, nitrogen, and sulfur cycle). Each arrow indicates a single step within the circle; genomes involved in each stage are next to each arrow; some genes encoding key enzymes are also included. Carbon fixation pathways: CBB – Calvin-Benson-Basham cycle; (r)TCA – (reverse) Krebs cycle. Wells from the sandstone (orange) and carbonate (blue) aquifers are shown.

## Methods

### Study site - The deep regional aquifers of the Negev Desert

The deep sandstone and the overlying carbonate rocks (Lower and Upper Cretaceous eras, respectively) are vital resources of fresh to brackish groundwater for domestic, agriculture, mining industries, and local desalination plants in the Negev Desert and the Sinai Peninsula (Israel and Egypt, respectively; **Fig.1**). The hydrogeological, hydro-geochemical and isotopic characteristics of the deep sandstone have been previously described in detail [30,31,46,83,84]. The immense sandstone aquifer is mainly composed of clastic sands and silts interbedded with thin clay layers [85]. The aquifer extends between the Arava and Suez rift valleys (**Fig. 1**) and has an estimated storage of several hundred billion cubic meters [86]. The aquifer lies at a considerable depth below the surface, typically hundreds of meters, over most of the basin, except for the southern recharge area in Sinai and along the northern Negev and Sinai anticlines, where the sandstone formations outcrop [86,87]. Due to the deep confinement, the aquifer is characterized by elevated groundwater temperatures and pressures of up to 60 °C and 140 atm, respectively. The overlying carbonate mainly comprises limestone and dolomite, with interbedded chalk, marl, and shales [88]. Although containing some ancient water (pre-Holocene), the carbonate aquifer is assumed to be currently recharged over its outcrops [36,37].

Although mostly confined, the sandstone seems to be connected to adjacent aquifers. This is supported by an intrusion of deeper, brackish, ancient (>600 kyr) groundwater body which was recently unveiled [30] as well as connectivity between the sandstone and the overlying carbonate aquifers. These two regional aquifers are separated by a low permeable layer over most of the basin, while in some areas, they might be locally connected [37,79]. In places, groundwater is diluted by younger water recharged through the limited sandstone outcrops along the northern Sinai and Negev anticlines (**Fig. 1**). Such a contribution could also result from the infiltration and cross-formational flow from shallower carbonate aquifer formations, facilitated by local fractures and faults. The combination of water stable isotope composition with the relative abundance of the ^81^Kr radioisotope, an emerging long-term age tracer, revealed different paleo-recharge components which flow in the two deep regional aquifers [29–31].

The sandstone aquifer is anoxic and characterized by elevated salinities, with Cl^-^ content ranging from 17 to 100 mmol L^-^ [31,80]. High sulfate levels of up to 9.3 mmol L^-1^ have been detected in wells from both aquifers [36,80]. In wells, Ein Yahav 6 (S5) and Zofar 220 (C4) (**Fig. 1**), sulfide concentrations were estimated as 470 µmol L^-1^ and 820 µmol L^-1^, respectively [89]. Sulfate reduction is likely a significant microbial process in the sandstone aquifer’s confined zone, as well as in the phreatic and confined parts of the carbonate aquifer [80,81].

### Field and laboratory methods

We collected groundwater samples from production wells in the Negev Desert and the Arava Valley(Fig. 1). Samples were collected from four production wells tapping the deep sandstone (S wells) and five from the overlain carbonate aquifer (C wells) (**Fig. 1**). The screens of the wells are perforated at various depths ranging from 471 to 718 mbgl for the carbonate aquifer wells; and from 765 to 1478 mbgl, for the sandstone aquifer wells.

Physical parameters, such as pH, electrical conductivity-EC, and DO, were monitored throughout the sampling period to ensure representative stable groundwater. Filtered water samples for major ions (SO_4_^-2^, Cl^-1^, NO_3_^-1^, Ca^+2^, Mg^+2^, Na^+1^, K^+1^), were collected in falcon tubes and measured and analyzed using by Dionex Ion Chromatography (ICS 5000) with Chromeleon software (Thermo Scientific) and by Inductively Coupled Plasma (SPECTRO Analytical Instruments GmbH). Concentrations of HCO_3_^-1^ were detected by automated titration-end point detection using a Methrohm titroprocesor (Metrohm AG). The analytical methods of δ^13^C and DIC measurement have been described in detail previously [90]. Briefly, samples for δ^13^C (results not shown) and DIC analyses were immediately filtered through 0.22 µm filters, stored at 4°C, and estimated using an isotopic ratio mass spectrophotometry (IRMS, DeltaV Advantage, Thermo). DIC is obtained as a byproduct of the δ^13^C analysis. TOC samples were acidified with HCl (pH<2) and measured on a Sievers InnovOX TOC analyzer (Veolia Water Technologies & Solutions). Ammonia nitrogen (NH_4_^+-^N) was determined by the Nessler method [91] using the wavelength of 420 nm on a Tecan i-control (Infinite 200). Historical sulfide (HS^-^) measurements were provided by Mekorot National Water Company and the Israeli Water Authority database.

### Primary productivity rates

Groundwater was sampled into pre-sterilized serum bottles (60 mL), overfilled from the bottom, and then crimp-sealed with rubber septa. We transported the samples from the field into the anaerobic hood (Coy Laboratory Products) and transferred them into 2 mL glass vials. The ^14^C bicarbonate stock solution (aqueous, activity 5 mCi/5 mL, PerkinElmer, Inc.) was diluted to a final specific activity of 5 µCi/50µL and added to the groundwater sample (43 mL, activity of 1 µCi/µL). Samples were incubated in slow rotation for 12 h at room temperature under dark conditions (Benchmark Scientific Roto-Therm Plus, H2024). At the end of the incubation, all the samples were filtered through glass fiber filters (GF/F, GF/F, Cytivia, 1825025). 50 µl of hydrochloric acid (HCl 37 %) was added to filters to purge the sample from non-assimilated bicarbonate overnight.

For the added activity (**See Supplementary Note 2**), 50 µl was taken at the beginning of incubation and placed in GF/F filters, and 50 µl of ethanolamine was added to block the purging of radiolabeled bicarbonate from samples. Radioactively labeled bicarbonate in the collected particulate material was detected as counts per minute (CPM) by liquid scintillation (Packard Tri-Carb 2100 TR Liquid Scintillation Analyzer). This method was adapted from previous work [92] with minor adjustments as we used the bicarbonate concentrations of each well as the dissolved inorganic carbon (DIC) value in equation 1 in Supplementary Note 2. The PP rates were measured in two different sampling times (Supplementary Table S1), except for samples C3 and C4, for which PP rates were measured just once.

### Secondary productivity rates

SP rates were estimated by measuring the assimilation rate of radioactively (^3^H) tagged leucine [93]. Briefly, sample handling was identical for PP measurements. In the anaerobic hood, 1.7 mL groundwater was transferred to a 2mL-HPLC vial x4(3 subsamples + 1 blank per well). 100 μL of leucine (^3^H-Leucine, Amersham, Specific activity: 160 Ci/mmol, radioactive concentration: 1.0 mCi/ml) was added to the samples, and they were incubated for 12 hours in dark conditions to a 2mL HPLC vial x4 (triplicates + 1 blank) in slow rotation as mentioned above. The incubation was terminated by adding trichloroacetic acid to the samples, and the samples were kept at 4 °C until measurements (**See Supplementary Note 2**).

### Cell abundance

Subsamples (1.7 mL groundwater) were fixed with glutaraldehyde (Sigma-Aldrich, G7651, final concentration, 0.2 %) for 10 min, snap frozen in liquid nitrogen, and stored at -80 °C until measurements. Samples were thawed at 37°C, and the total prokaryotes were stained with SYBR GREEN I (S7563, Invitrogen, final concentration, 1 nM, Ex = 497 nm, Em = 520). We analyzed the samples using an Attune-Next acoustic flow cytometer (Applied Biosystems), equipped with 350 and 450 nm lasers under a flow rate of 500 to 700 µl min^-1^ until it reached the threshold of 20,000 events per sample (minor adjustments from [94]).

### DNA sampling, extraction, and sequencing

Groundwater was pumped from active production wells into clean containers rinsed in situ with the abstracted groundwater before the sampling and transported to the laboratory facility for filtration between 12 to 24 hours after the collection. Samples S1 and C1 were collected in June 2021, while the rest (C2 to C5; and S2, S3, S5) in September 2021. Approximately 13 to 50 L of groundwater was filtered using a sterile Stainless Steel 142 mm Filter holder fitted with a 0.22 μm Durapore filter (Millipore-142 mm) (Cat. No. YY3014236, Millipore, Darmstadt, Germany). Filters were snap-frozen using liquid nitrogen and stored at -80 °C until DNA extraction with the DNeasy Power Water kit (Qiagen) using the manufacturer’s protocol.

Library preparation using the NEBNext® Ultra™ IIDNA Library Prep Kit (Cat No. E7645) and sequencing were performed at the Novogene AIT Genomics facility in Singapore. The nine DNA samples were sequenced using 150 bp paired-end reads on NovaSeq 6000 platform, with a read depth of ∼15 Gbp per sample.

### Assembly, binning, and metabolic predictions

The DNA reads were assembled with SPAdes V3.14 (--meta, k=21,33,66,99,127) [95], following adapter trimming and error correction with a tadpole.sh, using the BBtools suite (sourceforge.net/projects/bbmap/). Downstream mapping and binning of metagenome-assembled genomes (MAGs) were performed using DAStool, Vamb, Maxbin 2.0, and Metabat2 [96–99]) within the Atlas V2.11 framework [100], using the genome dereplication nucleotide identity threshold of 0.975. Functional annotation was done using DRAM [101] and METABOLIC [102], using default settings and a pathway completeness threshold 0.75. Key functions were verified using BLAST against the NCBI nr/nt and RefSeq databases [103]. Maximum growth rates were predicted using gRodon2 [104]. DNA raw reads and high-quality MAGs were deposited to NCBI with project accession number NCBI BioProject PRJNA983765.

### Statistical methods and data analyses

All data analysis, including statistics and visualization, was conducted in R version 4.2.3.[105]. We investigate microbial communities’ structure and their association with hydrogeochemistry using the R package Phyloseq [106]. Spearman’s correlation was conducted and visualized using an R package corrplot [107] to explore the correlation between the environmental variables and the PP, SP, and CA. Nonparametric tests Wilcoxson-Man Whitney test (MWT) was applied to examine the differences in PP, SP, and environmental variables between the aquifers. SigmaPlot 12.5 (Systat Software, Inc) was also used for data visualization.

## Supporting information

Supplementary calculation and data tables and figures

## Authors’ contributions

The study was conceived and funded by ZR, MR-B, and EA, and with the help of EG and MR-B, ZR, BA, RR, and EA, collected the samples. EBZ and EG performed the productivity measurements; MR-B performed the bioinformatics work. BA, MR-B, and ZR wrote the paper with contributions from all co-authors. The final manuscript was read and approved by all authors.

## Funding

This work was funded by the Israel Science Foundation (grant number:1068/20), MR-B is funded by the Ministry of Energy (Grant 221-17-002), and the Ministry of Science and Technology Grant (proposal # 001126). The research has also been supported by grants from BA by the National Secretariat of Science, Technology, and Innovation (SENACYT) of the Republic of Panama.

## Acknowledgment

The authors thank the Israel Water Authority and Mekorot Israel National Water Company for providing access to wells for sampling and sharing data. Special thanks are extended to Oded Beja, Alina Pushkarev, and Ariel Chazan (Technion-Israel Institute of Technology) for their assistance with the field sampling method. We also thank Damiana Diaz Reck, Almog Gafni, Adva Speter, Amos Russak, and Efrat Eliani Russak (Ben Gurion University of the Negev) for their precise analytical work and technical assistance; Alexandra Gonzalez (Technological University of Panama) is thanked for her assistance with preparing the location map.

## Availability of data and material

The raw metagenomic reads are available on NCBI as BioProject PRJNA983765.

## Declarations

### Ethics approval and consent to participate

Not applicable.

### Consent for publication

Not applicable.

### Competing interests

The authors declare that the research was conducted in the absence of any commercial or financial relationships that could be construed as a potential conflict of interest.

## Notes

https://www.ncbi.nlm.nih.gov/bioproject/PRJNA983765

### Competing Interest Statement

The authors have declared no competing interest.

