## Supplementary calculation and data tables and figures for "Metabolic adaptations and aquifer connectivity underpin the high productivity rates in the relict subsurface water"

1. Department of Environmental Hydrology and Microbiology, Zuckerberg Institute for Water Research. Ben-Gurion University of the Negev
2. Department of Marine Biology, Israel Oceanographic and Limnological Research
3. Geological Survey of Israel

### 1. Physicochemical description of water in the sampled wells

Groundwater analyses from the two aquifers showed distinct physical and hydrogeochemical features (Table 1). The median temperatures were 39 °C and 52 °C in the carbonate and deep sandstone aquifers. All the samples are within the category of hypoxic to anoxic conditions (<0.006 mmol $O_2$ $L^{-1}$ or 0.2 mg $L^{-1}$). Electrical conductivity (EC) levels were generally consistent among the samples, with a 3.2 mS cm $^{-1}$ range. However, sample S3 was an exception as it had elevated salinity levels, with an EC of 10.2 mS cm $^{-1}$. The redox potential in wells from both aquifers was below -200 mV (unpublished data).

Based on water stable isotopes [1–3], the origin of recharge also differed between the aquifers, where the deep sandstone aquifer contained more depleted $\delta^{18}O$ values compared to the overlain carbonate group (MWT, $p$=0.020), except for well S5 with higher $\delta^{18}O$ [1]. $^{81}Kr$ dating of the groundwater samples indicates that the residence time is more than 50 kyr [1–3]. The sandstone aquifer has a spatial age evolution along the flow path from S1 to S2, followed by a sharp increase of more than 200 kyr in well S3, [1]. Groundwater in well S3 may represent an intrusion from a deep and older highly pressurized saline aquifer (620 kyr, 51°C), which could have elevated the groundwater age and water temperature in the deep Nubian sandstone aquifer [1,4]. Further downstream, in well S5, the groundwater age suddenly decreases, which implies younger recharge to these parts of the deep Nubian sandstone aquifer [1].

Concentrations of potential electron acceptors for microbial metabolism varied across the samples. In the carbonate aquifer, the median 5.6 mmol $L^{-1}$ bicarbonate ($HCO_3^-$) concentration was higher compared to 3.79 mmol $L^{-1}$ in the deeper Nubian sandstone aquifer (MWT, $p$=0.031). On average, the sulfate ($SO_4^{-2}$) concentrations were 7.76 ± 2.04 mmol $L^{-1}$ in the sandstone aquifer and 8.34± 3.79 mmol $L^{-1}$ in the overlain aquifer. Nitrate ($NO_3^-$) and nitrite ($NO_2^-$) concentrations were below the detection limits(<0.016 mmol $L^{-1}$). The electron donors identified in the samples were sulfide ($HS^-$) and ammonium ($NH_4^+$), while methane was not detectable. Sulfide was above 4 μmol $L^{-1}$ in all the wells, with the highest concentration found in C4 from the carbonate aquifer (287 μmol $L^{-1}$). Ammonium concentration was slightly higher in the carbonate aquifer than in the deep sandstone aquifer (median values of 208 and 107 μmol $L^{-1}$, respectively).

### 2. Estimating rates of carbon assimilation

**The rates of inorganic carbon assimilation (PP)**

$$(\mathbf{1})\mathrm{PP}\left(\mu\mathrm{g\ C\ L^{-1^{-}}\ d^{-1}}\right) = \frac{\begin{gathered}(\mathrm{CPM\ Sample} - \mathrm{CPM\ Kill}\,) \times \mathrm{DIC}\ (mg\ L^{-1} \times \mathrm{Volume\ of\ added\ activity}(L) \times \\ 1.05 \times 1000 \times 24\ (h\ d^{-1})\end{gathered}}{\mathrm{Filtered\ Volume}\ (L) \times \mathrm{CPM\ of\ added\ activity} \times \mathrm{Incubation\ time}(h)}$$

Where: the volume of added activity is the subsample volume (50 µL); added activity is the total counts per minute (CPM) after spiking; 1.05 is the fractionation factor of uneven uptake of $^{14}C$; incubation time in h; DIC is the inorganic carbon of the system in mg $L^{-1}$ using 12 gr $mol^{-1}$. Sample filtered volume in this study was 0.043 L.

**Leucine uptake rate (SP)**

$$(\mathbf{2})\mathrm{SP}\left(\mu\mathrm{g\ C\ L^{-1^{-}}\ d^{-1}}\right) = \frac{\begin{gathered}(\mathrm{DPM\ Sample} - \mathrm{DPM\ Kill}\,) \times \mathrm{nmol\ Leu\ in\ 1\ DPM} \times \mathrm{biomass\ factor} \times \\ \mathrm{Isotope\ dilution} \times \mathrm{Cellular\ carbon} \times 24\ (h\ d^{-1})\end{gathered}}{1000 \times \mathrm{Sample\ Volume}\ (L) \times \mathrm{Incubation\ time}(h)}$$

Where: 1 nmol leucine (Leu) in 1 DPM= $1.31\times10^{-8}$; the biomass factor = nmol Leu to ng Leu= 1797; isotope dilution factor= 2 (non-even uptake of radioactive Leucine); cellular carbon per protein factor = ng Leu to ng Carbon=0.86; 1000 to convert ng C to µg of C; sample volume is 0.0017 L for this protocol.

The cocktail makes the radioactivity of the sample result in pulses of light that are later translated by the counter to Disintegrations Per Minute (DPM) or Counts Per Minute (CPM).

### 3. Correlation factors between primary productivity rates and environmental factors

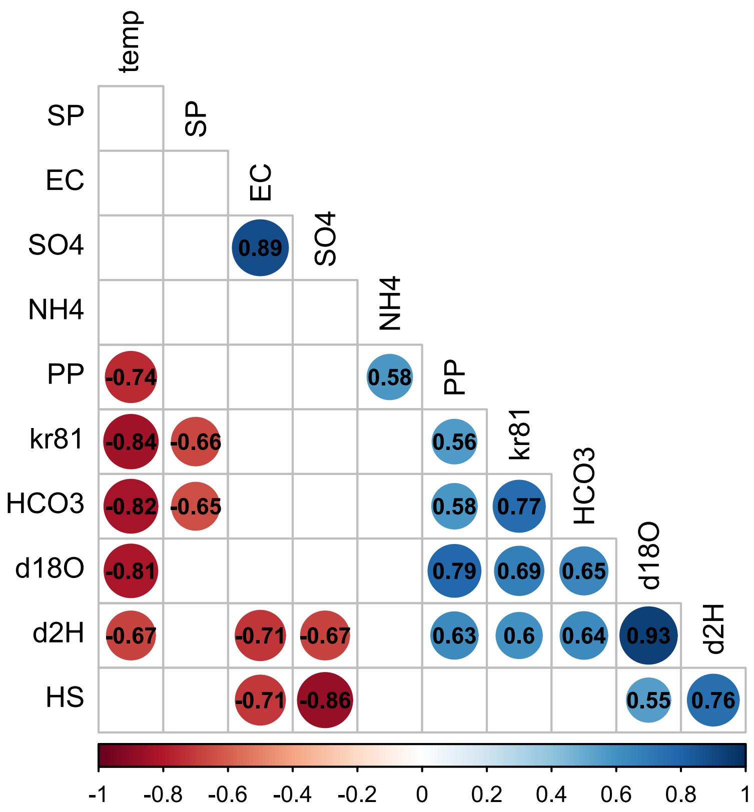


**Figure S1.** Spearman correlation factors between the PP rates and environmental factors; only significant correlation factors are added ($p \leq 0.05$). PP rates were positively associated with $\delta^{18}O$, $\delta^{2}H$, $^{81}Kr$ ratio, $HCO_3^-$, and $NH_4^+$. We did not use the averaged PP rates due to the considerable variation between replicates within the samples. Thus, we compared the individual replicates of PP rates. We excluded the data from C1 as it did not contain PP rates.

### 4. Secondary heterotrophic productivity in aquifers

Despite the high PP, the SP ranged between 0.005 to 0.07 µg C $L^{-1}$ $d^{-1}$ across both aquifers (**Table 1**), with the highest rate observed in well S3. We note that two wells from the carbonate aquifer, C2, and C4, with the lowest SP values, had the highest PP rates. Our data suggests so far that the heterotrophic secondary production in two deep, pristine aquifers is limited by the relatively low levels of organic carbon available in these habitats. Therefore, the SP rates remained low throughout the samples from both aquifers.

However, we did not detect any significant relationship between the measured SP and PP rates. Previous research has also shown extremely low heterotrophic production of up to 75.6 pmol C $L^{-1}$ $hr^{-1}$ (or 0.02 µg C $L^{-1}$ $d^{-1}$) in groundwater from an alpine karst aquifer with low dissolved organic carbon of up to 0.046 mmol $L^{-1}$ and cell abundance of 1 $x10^{6}$ cells $L^{-1}$, where the authors concluded that most of the planktonic cells are inactive or in a dormant state [5]. In this study, TOC did not vary between the aquifers with a median value of 0.07 mmol $L^{-1}$ (~ < 1 mg $L^{-1}$) (**Table 2**). These values are lower than the global mean of TOC in lake water of 5.6 mg $L^{-1}$[6,7].

No significant relationship was observed between the SP and the TNC, or TOC, which suggests that the systems may not be driven solely by the amount of available organic carbon and nutrients [8,9]. These observations are consistent with previous findings from an oligotrophic aquifer in the Bavarian Alps, where the lack of correlation between microbial production, abundance, and bioavailable carbon sources, suggests that the low levels of nutrients and low accessible carbon may not be sufficient to sustain the planktonic prokaryotic communities [9]. Furthermore, it has been noted that a rise in organic carbon over time does not always lead to enhancement in secondary production, which often may be attributed to scarcity of essential nutrients [9–11]. Furthermore, our SP rates were several folds lower than the PP rates, indicating that heterotrophic prokaryotes may not be dominant but rather strict or facultative chemoautotrophs.

### 5. Alpha and Beta diversity of the microbial communities

The diversity of the communities in both aquifers is similar (mean Shannon index of 2.44 ± 0.65 and 1.66 ± 0.80, in sandstone and carbonate aquifers, respectively, MWT $p$=0.19) (Fig. S2a). At higher taxonomic levels, Halothiobacilalles, Burkholderiales, Ammonifexales, and Rhizobiales were abundant. Besides these abundant microbial communities, several taxa were only present in a few wells. For instance, in well C2, we found unique communities belonging to the phylum Desulfobacterota and UBA9089, respectively, known as sulfate reducers [12]. DTGQ01 within the new phyla SZUA-182 was only detected in well S5 and C2, previously reported in serpentinized subsurface fluids and spring anoxic sediments encoding the capacity for sulfite reduction [13,14]. In well S3, which has a unique hydrochemical and isotopic characteristic, we identified members of Desulfotomaculales, including *Desulfotomacullum profundis*, an anaerobic and moderately thermophilic organism capable of growing autotrophically or heterotrophically [15–17].

Comparison of the microbial communities via principal component analysis (PCoA) using Bray-Curtis distances revealed no significant difference between the samples from the two aquifers (Adonis vegan package p=0.208, Fig. S2b). As shown in the S1b, there is an overlap between the two groups, where specific samples from the sandstone aquifer contained similar communities to those of the carbonate aquifer. Samples C1 and S1 clustered in the PCoA plot due to similar community composition, even though they are drilled in different aquifers and separated 10 m from each other. Likewise, sample S2 contains high proportions of the carbonate aquifer's dominant community. Thus, these similarities may indicate that the aquifers are connected, and potential mixing processes may occur between the aquifers and other unknown sources.

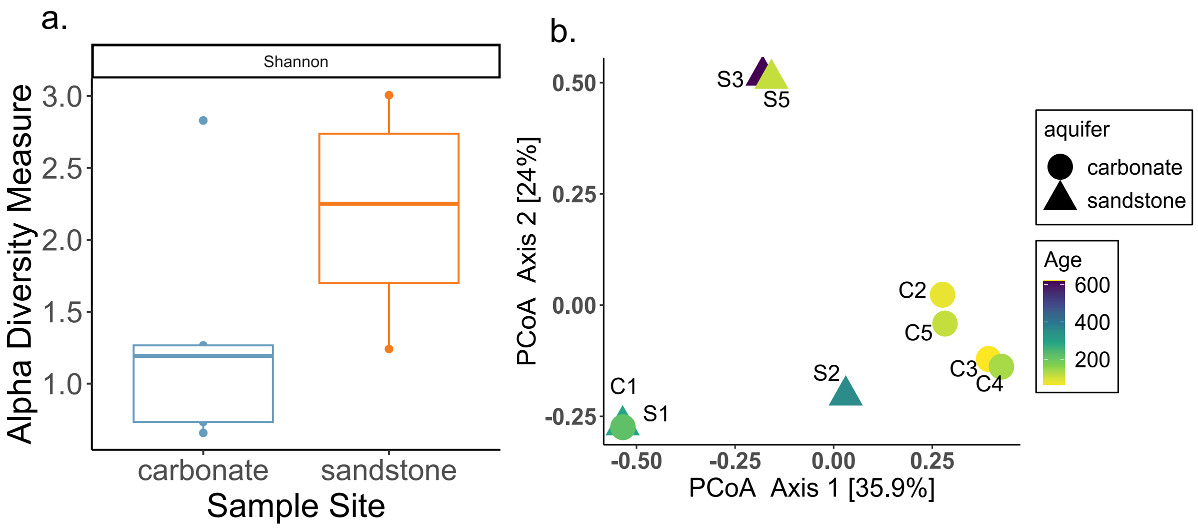


**Figure S2.** (a) Shannon diversity indexes in the aquifers. (b) PCoA analysis between the samples of the two aquifers. Age in kyr.

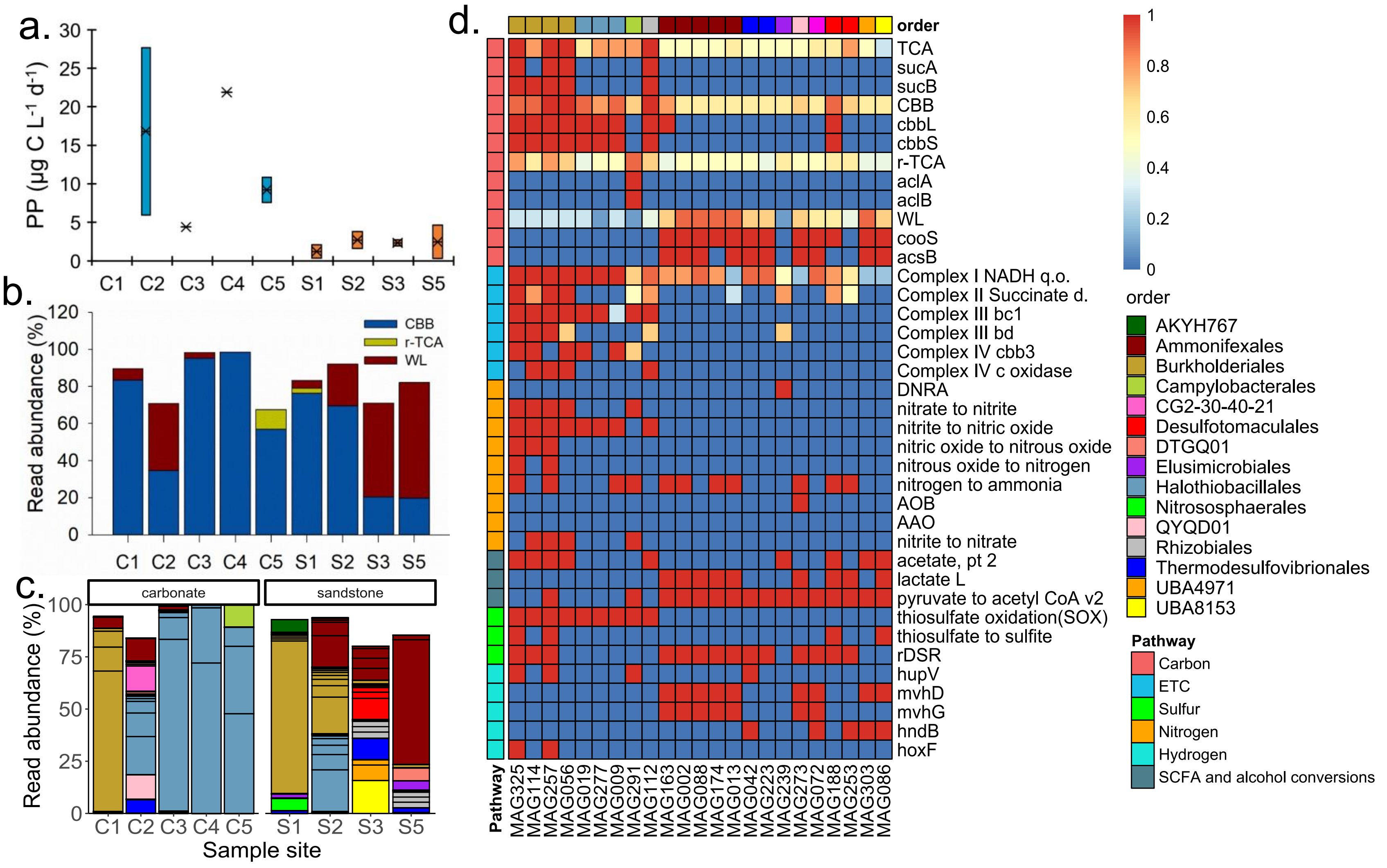


**Figure S3**. (a) Primary productivity rates across all samples, except C1; (b) cumulative read abundance of MAGs encoding for carbon fixation pathways; (c) read abundance of the top 15 MAGs on the order level; and (d) completion pathways of the TCA, CBB, r-TCA, WL, and Electron transport chain (ETC) complexes; presence/absence of genes and metabolic pathways in selected MAGs (DRAM output). *SCFA= Short chain fatty acids. AOB= ammonia oxidation by bacteria-aerobic specific). AOA= ammonia oxidation by archaea. DNRA = dissimilatory nitrite reduction to ammonia. rDSR= dissimilatory sulfate reduction (and oxidation) sulfate to sulfide. sucAB [EC:1.2.4.2 2.3.1.61 1.8.1.4]; cooS [EC:1.2.7.4]; acsB [EC:2.3.1.169]; cbbL/S [EC:4.1.1.39]; hupV [EC:1.12.99.6]; mvhGD [EC:1.12.99.- 1.8.98.5 1.8.98.6]; hndB [EC:1.12.1.3]; hoxF [EC:1.12.1.2].*

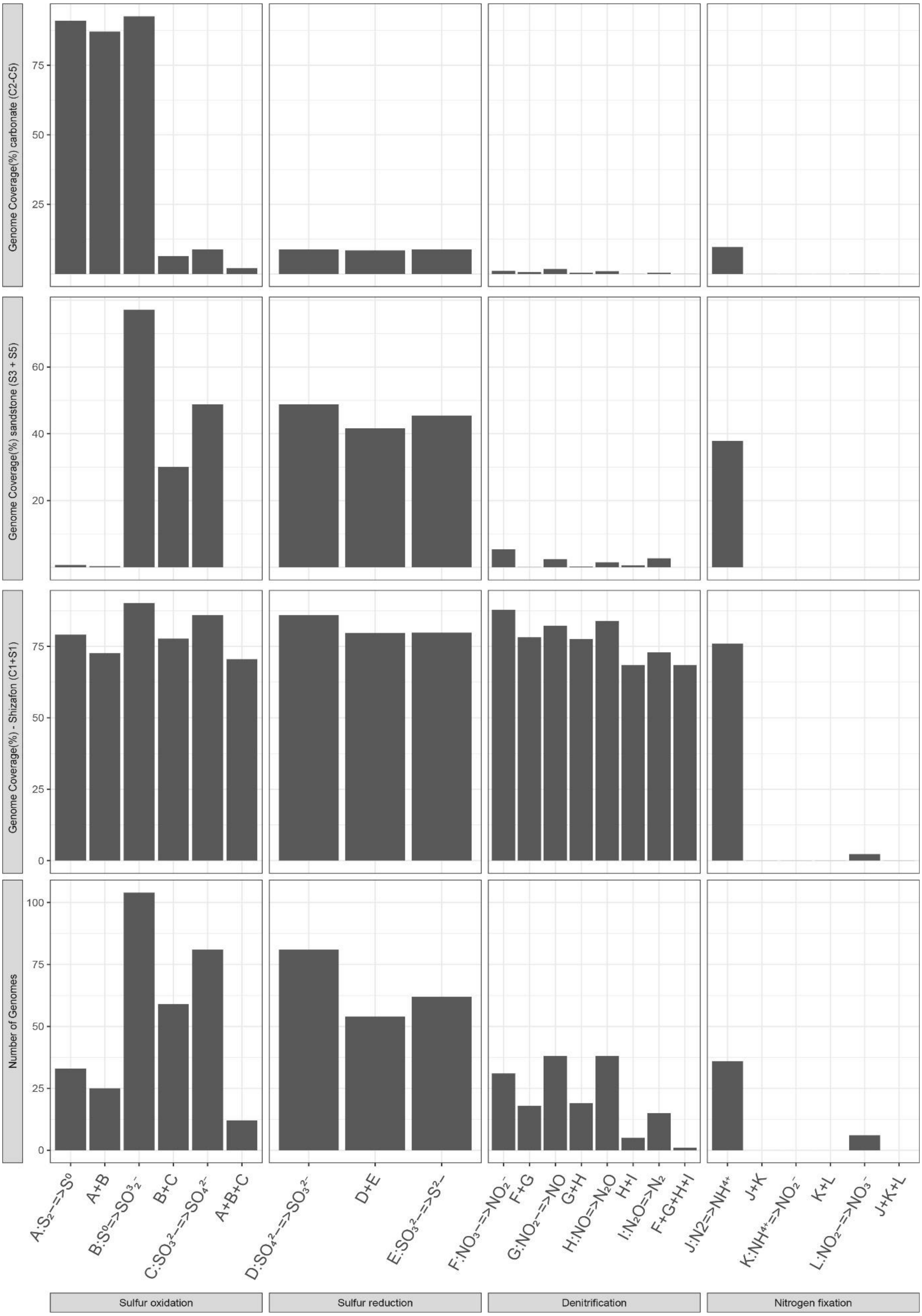


**Figure S4.** Number and read abundance of metagenome-assembled genomes (MAGs) involved in sequential redox transformations.

### Table S1. Estimation of carbon assimilation rates-PP rates

| Well name | ID | aquifer | Month | DIC (mg $L^{-1}$) | A= (DPM) (at time=0) | B=Blank (DPM) | A-B (DPM) | added activity (DPM) | PP (µg C $L^{-1}$ $d^{-1}$) |
|---|---|---|---|---|---|---|---|---|---|
| Paran 129 | C2 | carbonate | Sept, 21 | 106.8 | 708 | 17 | 691 | 30290 | 6.0 |
| Paran 129 | C2 | carbonate | July, 22 | 106.8 | 534 | 17 | 517 | 4876 | 27.7 |
| Paran 21 | C3 | carbonate | Sept, 21 | 105.3 | 1904 | 17 | 1887 | 110580 | 4.4 |
| Zofar 220 | C4 | carbonate | Sept, 21 | 91.7 | 2586 | 17 | 2569 | 26292 | 21.9 |
| Ein Yahav 7 | C5 | carbonate | Sept, 21 | 62.0 | 2128 | 17 | 2111 | 42072 | 7.6 |
| Ein Yahav 7 | C5 | carbonate | July, 22 | 62.0 | 152 | 17 | 135 | 1894 | 10.8 |
| Shizafon 1 | S1 | sandstone | Nov,21 | 59.9 | 71 | 23 | 48 | 3358 | 2.1 |
| Shizafon 1 | S1 | sandstone | Dec, 21 | 59.9 | 60 | 17 | 43 | 19201 | 0.3 |
| Paran 20 | S2 | sandstone | Sept, 21 | 51.5 | 513 | 17 | 496 | 38904 | 1.6 |
| Paran 20 | S2 | sandstone | July, 22 | 51.5 | 126 | 17 | 109 | 3623 | 3.8 |
| Zofar20 | S3 | sandstone | Sept, 21 | 42.0 | 2736 | 17 | 2719 | 102417 | 2.7 |
| Zofar 20 | S3 | sandstone | Dec, 21 | 42.0 | 75 | 17 | 58 | 3096 | 1.9 |
| Ein Yahav 6 | S5 | sandstone | July, 22 | 66.8 | 101 | 17 | 84 | 2975 | 4.6 |
| Ein Yahav 6 | S5 | sandstone | Sept, 21 | 66.8 | 797 | 17 | 780 | 435163 | 0.3 |
| *Incubation Temperature circa 50°C (not considered/discussed in the main text) | | | | | | | | | |
| Shizafon 1* | S1 | sandstone | Nov,21 | 59.9 | 101 | 23 | 78 | 3358 | 3.4 |
| Shizafon 1* | S1 | sandstone | Dec,21 | 59.9 | 100 | 17 | 83 | 1431 | 8.5 |
| Zofar 20* | S3 | sandstone | Dec,21 | 42.0 | 177 | 17 | 160 | 2124 | 7.7 |
| Paran 20* | S2 | sandstone | July,22 | 51.5 | 636 | 17 | 619 | 1650 | 47.2 |

### Table S2. Estimation of carbon assimilation based on the leucine assay (SP rates)

| Well name | ID | Month | Dpm treatment | Dpm kill | Dpm (treatment-kill) | Incubation time (h) | SP rate (µg C $L^{-1}$ $d^{-1}$) | Mean SP (µg C $L^{-1}$ $d^{-1}$) | Standad Deviation |
|---|---|---|---|---|---|---|---|---|---|
| Shizafon 11 | C1 | Jun, 21 | 658 | 405 | 253 | 12 | 0.012 | 0.0553 | 0.079 |
| | | | 565 | 405 | 160 | 12 | 0.008 | | |
| | | | 3473 | 405 | 3068 | 12 | 0.146 | | |
| Shizafon1 | S1 | Nov, 21 | 469 | 107 | 362 | 12 | 0.017 | 0.0165 | 0.001 |
| | | | 441 | 107 | 334 | 12 | 0.016 | | |
| | | | 451 | 107 | 344 | 12 | 0.016 | | |
| Paran 20 | S2 | Sep, 21 | 219 | 20 | 199 | 12 | 0.009 | 0.0173 | 0.007 |
| | | | 405 | 20 | 385 | 12 | 0.018 | | |
| | | | 523 | 20 | 503 | 12 | 0.024 | | |
| Paran 21 | C3 | Sep, 21 | 262 | 12 | 250 | 12 | 0.012 | 0.0057 | 0.005 |
| | | | 60 | 12 | 48 | 12 | 0.002 | | |
| | | | 75 | 12 | 63 | 12 | 0.003 | | |
| Paran129 | C2 | Sep, 21 | 110 | 26 | 84 | 12 | 0.004 | 0.0049 | 0.0033 |
| | | | 207 | 26 | 181 | 12 | 0.009 | | |
| | | | 72 | 26 | 46 | 12 | 0.002 | | |
| Zofar220 | C4 | Sep, 21 | 27 | 26 | 1 | 12 | 0.00005 | 0.0004 | 0.0003 |
| | | | 43 | 26 | 17 | 12 | 0.001 | | |
| | | | 31 | 26 | 5 | 12 | 0.0002 | | |
| Zofar20 | S3 | Sep, 21 | 56 | 18 | 38 | 12 | 0.002 | 0.0720 | 0.1203 |
| | | | 4444 | 18 | 4426 | 12 | 0.211 | | |
| | | | 90 | 18 | 72 | 12 | 0.003 | | |
| E.Yahav7 | C5 | Sep, 21 | 286 | 25 | 261 | 12 | 0.012 | 0.0523 | 0.0759 |
| | | | 121 | 25 | 96 | 12 | 0.005 | | |
| | | | 2958 | 25 | 2933 | 12 | 0.140 | | |
| E.Yahav6 | S5 | Sep, 21 | 68 | 24 | 44 | 12 | 0.002 | 0.0054 | 0.0050 |
| | | | 86 | 24 | 62 | 12 | 0.003 | | |
| | | | 257 | 24 | 233 | 12 | 0.011 | | |

### Table S3. Presence/absence of sulfur pathways.

| Function | Sulfide oxidation | Sulfur oxidation | Sulfur oxidation | Sulfur oxidation | Sulfur oxidation/reduction | Thiosulfate oxidation | Sulfate reduction | Sulfate reduction | Thiosulfate disproportionation |
|---|---|---|---|---|---|---|---|---|---|
| Gene | sqr | dsrA | dsrB | sdo | sor | soxB | aprA | sat | phsA |
| MAG001 | Absent | Absent | Absent | Present | Absent | Absent | Absent | Absent | Absent |
| MAG002 | Absent | Present | Present | Present | Absent | Absent | Present | Present | Absent |
| MAG003 | Absent | Absent | Absent | Absent | Absent | Absent | Absent | Absent | Absent |
| MAG005 | Absent | Absent | Absent | Absent | Absent | Absent | Absent | Present | Absent |
| MAG006 | Absent | Absent | Absent | Present | Absent | Present | Absent | Absent | Absent |
| MAG008 | Absent | Absent | Absent | Absent | Absent | Absent | Absent | Absent | Absent |
| MAG009 | Present | Absent | Absent | Present | Absent | Present | Absent | Absent | Absent |
| MAG011 | Present | Absent | Absent | Present | Absent | Present | Absent | Absent | Absent |
| MAG012 | Absent | Present | Present | Present | Absent | Absent | Present | Present | Present |
| MAG013 | Absent | Present | Present | Present | Absent | Absent | Present | Present | Absent |
| MAG014 | Absent | Present | Present | Absent | Absent | Absent | Present | Present | Absent |
| MAG018 | Absent | Absent | Absent | Present | Absent | Absent | Absent | Absent | Absent |
| MAG019 | Present | Absent | Absent | Present | Absent | Present | Absent | Absent | Absent |
| MAG020 | Absent | Absent | Absent | Present | Absent | Absent | Absent | Present | Absent |
| MAG025 | Present | Present | Present | Present | Absent | Absent | Present | Present | Present |
| MAG026 | Absent | Absent | Absent | Absent | Absent | Absent | Absent | Present | Absent |
| MAG027 | Absent | Absent | Absent | Present | Absent | Absent | Present | Absent | Absent |
| MAG028 | Absent | Absent | Absent | Present | Absent | Absent | Absent | Present | Absent |
| MAG030 | Absent | Absent | Absent | Absent | Absent | Absent | Absent | Absent | Absent |
| MAG037 | Absent | Absent | Absent | Present | Absent | Absent | Absent | Absent | Absent |
| MAG038 | Absent | Absent | Absent | Present | Absent | Absent | Absent | Absent | Absent |
| MAG042 | Absent | Present | Present | Present | Absent | Absent | Present | Present | Absent |
| MAG044 | Absent | Present | Present | Present | Absent | Absent | Present | Present | Absent |
| MAG045 | Absent | Absent | Absent | Present | Absent | Absent | Absent | Present | Absent |
| MAG046 | Present | Present | Present | Present | Absent | Present | Present | Present | Absent |
| MAG051 | Absent | Absent | Absent | Present | Absent | Absent | Absent | Present | Absent |
| MAG052 | Absent | Absent | Absent | Absent | Absent | Absent | Absent | Absent | Absent |
| MAG053 | Absent | Present | Present | Absent | Absent | Absent | Absent | Absent | Absent |
| MAG054 | Absent | Absent | Absent | Absent | Absent | Absent | Absent | Absent | Absent |
| MAG055 | Absent | Present | Present | Present | Absent | Absent | Absent | Absent | Absent |
| MAG056 | Present | Absent | Absent | Present | Present | Present | Absent | Absent | Absent |
| MAG057 | Absent | Present | Present | Present | Absent | Absent | Present | Present | Absent |
| MAG059 | Absent | Absent | Absent | Present | Absent | Absent | Absent | Absent | Absent |

| Function | Sulfide oxidation | Sulfur oxidation | Sulfur oxidation | Sulfur oxidation | Sulfur oxidation/reduction | Thiosulfate oxidation | Sulfate reduction | Sulfate reduction | Thiosulfate disproportionation |
|---|---|---|---|---|---|---|---|---|---|
| MAG062 | Absent | Present | Present | Present | Absent | Absent | Present | Present | Absent |
| MAG064 | Absent | Present | Present | Present | Absent | Absent | Present | Present | Absent |
| MAG066 | Absent | Absent | Absent | Present | Absent | Absent | Absent | Absent | Absent |
| MAG067 | Present | Absent | Absent | Absent | Absent | Present | Absent | Present | Absent |
| MAG069 | Absent | Absent | Absent | Present | Absent | Absent | Absent | Absent | Absent |
| MAG071 | Absent | Absent | Absent | Present | Absent | Absent | Absent | Present | Absent |
| MAG072 | Present | Present | Present | Present | Absent | Absent | Present | Present | Absent |
| MAG073 | Absent | Absent | Absent | Absent | Absent | Absent | Absent | Absent | Absent |
| MAG074 | Absent | Present | Present | Present | Absent | Absent | Absent | Absent | Present |
| MAG079 | Absent | Absent | Absent | Present | Absent | Present | Absent | Absent | Absent |
| MAG080 | Absent | Absent | Absent | Absent | Absent | Absent | Absent | Absent | Absent |
| MAG081 | Absent | Present | Present | Present | Absent | Absent | Absent | Present | Absent |
| MAG085 | Absent | Absent | Absent | Absent | Absent | Absent | Absent | Absent | Absent |
| MAG086 | Absent | Absent | Absent | Present | Absent | Absent | Absent | Absent | Present |
| MAG088 | Absent | Present | Present | Present | Absent | Absent | Present | Present | Absent |
| MAG089 | Absent | Absent | Absent | Absent | Absent | Absent | Absent | Present | Absent |
| MAG091 | Absent | Absent | Absent | Present | Absent | Present | Absent | Absent | Absent |
| MAG096 | Present | Absent | Absent | Present | Absent | Present | Absent | Absent | Absent |
| MAG101 | Absent | Present | Present | Present | Absent | Absent | Present | Present | Present |
| MAG102 | Absent | Absent | Absent | Present | Absent | Absent | Absent | Absent | Present |
| MAG104 | Absent | Present | Present | Present | Absent | Absent | Absent | Absent | Absent |
| MAG106 | Absent | Absent | Absent | Present | Absent | Present | Absent | Absent | Absent |
| MAG107 | Absent | Present | Present | Present | Absent | Absent | Present | Present | Absent |
| MAG109 | Present | Absent | Absent | Present | Absent | Present | Absent | Absent | Absent |
| MAG110 | Absent | Absent | Absent | Present | Absent | Absent | Absent | Absent | Absent |
| MAG111 | Present | Absent | Absent | Present | Absent | Absent | Absent | Absent | Absent |
| MAG112 | Absent | Absent | Absent | Present | Absent | Absent | Absent | Absent | Absent |
| MAG114 | Present | Present | Present | Present | Absent | Absent | Present | Present | Absent |
| MAG115 | Absent | Absent | Absent | Present | Absent | Absent | Absent | Absent | Absent |
| MAG118 | Absent | Absent | Absent | Present | Absent | Absent | Absent | Present | Absent |
| MAG120 | Absent | Absent | Absent | Present | Absent | Absent | Absent | Present | Absent |
| MAG121 | Absent | Absent | Absent | Present | Absent | Absent | Absent | Present | Absent |
| MAG124 | Absent | Present | Present | Present | Absent | Absent | Present | Present | Present |
| MAG127 | Absent | Absent | Absent | Present | Absent | Absent | Absent | Absent | Present |
| MAG132 | Absent | Absent | Absent | Present | Absent | Absent | Absent | Absent | Absent |

| Function | Sulfide oxidation | Sulfur oxidation | Sulfur oxidation | Sulfur oxidation | Sulfur oxidation/reduction | Thiosulfate oxidation | Sulfate reduction | Sulfate reduction | Thiosulfate disproportionation |
|---|---|---|---|---|---|---|---|---|---|
| MAG133 | Present | Absent | Absent | Present | Absent | Absent | Absent | Present | Present |
| MAG134 | Absent | Absent | Absent | Absent | Absent | Absent | Absent | Present | Absent |
| MAG137 | Absent | Absent | Absent | Present | Absent | Absent | Absent | Present | Absent |
| MAG138 | Absent | Absent | Present | Absent | Absent | Absent | Present | Present | Absent |
| MAG142 | Present | Absent | Absent | Absent | Absent | Absent | Absent | Present | Absent |
| MAG144 | Absent | Absent | Absent | Present | Absent | Absent | Absent | Present | Absent |
| MAG149 | Present | Absent | Absent | Present | Absent | Absent | Absent | Absent | Absent |
| MAG154 | Absent | Absent | Absent | Present | Absent | Absent | Absent | Absent | Absent |
| MAG156 | Absent | Present | Present | Present | Absent | Absent | Present | Present | Present |
| MAG158 | Absent | Present | Present | Present | Absent | Absent | Present | Present | Present |
| MAG160 | Absent | Present | Present | Present | Absent | Absent | Present | Present | Absent |
| MAG162 | Absent | Present | Present | Present | Absent | Absent | Present | Present | Absent |
| MAG163 | Absent | Present | Present | Absent | Absent | Absent | Present | Present | Absent |
| MAG164 | Absent | Absent | Absent | Absent | Absent | Absent | Absent | Absent | Absent |
| MAG166 | Absent | Absent | Present | Absent | Absent | Absent | Absent | Absent | Absent |
| MAG170 | Absent | Absent | Absent | Present | Absent | Absent | Absent | Absent | Present |
| MAG174 | Absent | Present | Present | Present | Absent | Absent | Present | Present | Absent |
| MAG175 | Absent | Absent | Absent | Absent | Absent | Absent | Absent | Present | Absent |
| MAG176 | Absent | Absent | Absent | Present | Absent | Absent | Absent | Present | Absent |
| MAG181 | Present | Present | Present | Absent | Absent | Absent | Absent | Absent | Absent |
| MAG184 | Absent | Present | Present | Absent | Absent | Absent | Present | Present | Absent |
| MAG187 | Absent | Absent | Absent | Present | Absent | Absent | Absent | Absent | Absent |
| MAG188 | Absent | Present | Present | Present | Absent | Absent | Present | Present | Present |
| MAG189 | Absent | Absent | Absent | Absent | Absent | Absent | Absent | Present | Absent |
| MAG190 | Absent | Present | Present | Present | Absent | Absent | Absent | Present | Absent |
| MAG191 | Absent | Present | Present | Absent | Absent | Absent | Absent | Present | Present |
| MAG196 | Absent | Present | Present | Present | Absent | Absent | Present | Present | Present |
| MAG197 | Present | Absent | Absent | Present | Absent | Absent | Absent | Present | Absent |
| MAG198 | Present | Present | Present | Present | Absent | Present | Present | Present | Absent |
| MAG203 | Absent | Present | Present | Present | Absent | Present | Present | Present | Absent |
| MAG204 | Absent | Absent | Absent | Present | Absent | Present | Absent | Absent | Absent |
| MAG205 | Absent | Absent | Absent | Absent | Absent | Absent | Absent | Absent | Absent |
| MAG206 | Present | Absent | Absent | Absent | Absent | Absent | Absent | Absent | Present |
| MAG207 | Absent | Absent | Absent | Absent | Absent | Absent | Absent | Absent | Absent |
| MAG217 | Present | Absent | Absent | Present | Absent | Present | Absent | Absent | Absent |

| Function | Sulfide oxidation | Sulfur oxidation | Sulfur oxidation | Sulfur oxidation | Sulfur oxidation/reduction | Thiosulfate oxidation | Sulfate reduction | Sulfate reduction | Thiosulfate disproportionation |
|---|---|---|---|---|---|---|---|---|---|
| MAG220 | Absent | Present | Present | Present | Absent | Absent | Present | Present | Present |
| MAG221 | Absent | Present | Present | Present | Absent | Absent | Present | Present | Absent |
| MAG222 | Absent | Absent | Absent | Absent | Absent | Absent | Absent | Absent | Absent |
| MAG223 | Absent | Present | Present | Present | Absent | Absent | Present | Present | Absent |
| MAG225 | Absent | Absent | Absent | Present | Absent | Absent | Absent | Absent | Absent |
| MAG230 | Absent | Present | Present | Present | Absent | Absent | Present | Present | Absent |
| MAG231 | Absent | Absent | Absent | Absent | Absent | Absent | Absent | Absent | Absent |
| MAG232 | Absent | Absent | Absent | Present | Absent | Absent | Absent | Absent | Present |
| MAG237 | Absent | Absent | Absent | Absent | Absent | Absent | Absent | Absent | Present |
| MAG238 | Absent | Present | Present | Present | Absent | Absent | Present | Present | Absent |
| MAG239 | Absent | Absent | Absent | Absent | Absent | Absent | Absent | Absent | Absent |
| MAG240 | Absent | Absent | Absent | Present | Absent | Absent | Absent | Absent | Absent |
| MAG242 | Absent | Present | Present | Present | Absent | Absent | Absent | Present | Absent |
| MAG247 | Absent | Absent | Absent | Present | Absent | Absent | Absent | Absent | Present |
| MAG249 | Present | Present | Present | Present | Absent | Present | Present | Present | Absent |
| MAG252 | Present | Present | Present | Present | Absent | Present | Present | Present | Absent |
| MAG253 | Absent | Present | Present | Present | Absent | Absent | Present | Present | Absent |
| MAG257 | Present | Present | Present | Absent | Absent | Present | Present | Present | Absent |
| MAG261 | Absent | Present | Present | Absent | Absent | Absent | Present | Present | Absent |
| MAG271 | Absent | Absent | Absent | Absent | Absent | Absent | Absent | Present | Absent |
| MAG273 | Absent | Present | Present | Present | Absent | Absent | Present | Present | Absent |
| MAG277 | Present | Absent | Absent | Present | Absent | Present | Absent | Absent | Absent |
| MAG279 | Absent | Present | Present | Present | Absent | Absent | Present | Present | Present |
| MAG282 | Absent | Absent | Absent | Present | Absent | Absent | Present | Present | Absent |
| MAG284 | Present | Absent | Absent | Present | Absent | Absent | Absent | Absent | Absent |
| MAG285 | Present | Absent | Absent | Present | Absent | Absent | Absent | Absent | Absent |
| MAG286 | Absent | Present | Present | Present | Absent | Absent | Absent | Absent | Absent |
| MAG291 | Present | Absent | Absent | Absent | Absent | Absent | Absent | Absent | Absent |
| MAG292 | Absent | Absent | Absent | Absent | Absent | Absent | Absent | Present | Absent |
| MAG293 | Present | Absent | Absent | Present | Absent | Absent | Absent | Absent | Absent |
| MAG294 | Absent | Absent | Absent | Present | Absent | Absent | Absent | Present | Absent |
| MAG301 | Absent | Absent | Absent | Absent | Absent | Absent | Absent | Absent | Absent |
| MAG303 | Absent | Absent | Absent | Present | Absent | Absent | Absent | Absent | Absent |
| MAG306 | Absent | Absent | Absent | Present | Absent | Absent | Absent | Absent | Absent |
| MAG307 | Present | Absent | Absent | Present | Absent | Present | Present | Present | Present |

| Function | Sulfide oxidation | Sulfur oxidation | Sulfur oxidation | Sulfur oxidation | Sulfur oxidation/reduction | Thiosulfate oxidation | Sulfate reduction | Sulfate reduction | Thiosulfate disproportionation |
|---|---|---|---|---|---|---|---|---|---|
| MAG309 | Present | Present | Present | Absent | Absent | Present | Present | Present | Absent |
| MAG310 | Present | Present | Present | Absent | Absent | Absent | Absent | Present | Absent |
| MAG311 | Absent | Present | Present | Absent | Absent | Absent | Present | Present | Present |
| MAG312 | Absent | Present | Present | Absent | Absent | Absent | Present | Present | Absent |
| MAG318 | Present | Absent | Absent | Present | Absent | Absent | Present | Present | Present |
| MAG320 | Absent | Absent | Absent | Absent | Absent | Absent | Absent | Absent | Absent |
| MAG321 | Absent | Present | Present | Present | Absent | Absent | Absent | Present | Absent |
| MAG325 | Present | Present | Present | Present | Absent | Present | Present | Present | Absent |
| MAG328 | Absent | Present | Present | Absent | Absent | Absent | Present | Absent | Absent |
| MAG330 | Absent | Present | Present | Present | Absent | Absent | Absent | Present | Absent |

### Table S4. Presence/absence of carbon fixation genes.

| Function | CBB cycle - Rubisco | | Wood Ljungdahl pathway | | | Reverse TCA cycle | |
|---|---|---|---|---|---|---|---|
| Gene | Form I | Form II | cdhD | cdhE | cooS | aclA | aclB |
| MAG001 | Absent | Absent | Absent | Absent | Absent | Absent | Absent |
| MAG002 | Absent | Absent | Present | Present | Present | Absent | Absent |
| MAG003 | Absent | Absent | Absent | Absent | Absent | Absent | Absent |
| MAG005 | Absent | Absent | Present | Present | Present | Absent | Absent |
| MAG006 | Absent | Absent | Absent | Absent | Absent | Absent | Absent |
| MAG008 | Absent | Absent | Absent | Absent | Absent | Absent | Absent |
| MAG009 | Present | Present | Absent | Absent | Absent | Absent | Absent |
| MAG011 | Present | Present | Absent | Absent | Absent | Absent | Absent |
| MAG012 | Absent | Absent | Absent | Absent | Present | Absent | Absent |
| MAG013 | Absent | Absent | Present | Present | Present | Absent | Absent |
| MAG014 | Absent | Absent | Absent | Absent | Absent | Absent | Absent |
| MAG018 | Present | Absent | Absent | Absent | Absent | Absent | Absent |
| MAG019 | Present | Present | Absent | Absent | Absent | Absent | Absent |
| MAG020 | Absent | Absent | Absent | Absent | Present | Absent | Absent |
| MAG025 | Absent | Absent | Present | Present | Present | Absent | Absent |
| MAG026 | Absent | Absent | Absent | Absent | Absent | Absent | Absent |
| MAG027 | Absent | Absent | Absent | Absent | Absent | Present | Present |
| MAG028 | Absent | Absent | Absent | Absent | Absent | Absent | Absent |
| MAG030 | Absent | Absent | Present | Present | Present | Absent | Absent |

| Function | CBB cycle - Rubisco | | Wood Ljungdahl pathway | | | Reverse TCA cycle | |
|---|---|---|---|---|---|---|---|
| MAG037 | Absent | Absent | Absent | Absent | Absent | Absent | Absent |
| MAG038 | Absent | Absent | Absent | Absent | Absent | Absent | Absent |
| MAG042 | Absent | Absent | Present | Present | Present | Absent | Absent |
| MAG044 | Absent | Absent | Present | Present | Present | Absent | Absent |
| MAG045 | Absent | Absent | Absent | Absent | Absent | Absent | Absent |
| MAG046 | Present | Present | Absent | Absent | Absent | Absent | Absent |
| MAG051 | Present | Absent | Absent | Absent | Absent | Absent | Absent |
| MAG052 | Absent | Absent | Present | Present | Present | Absent | Absent |
| MAG053 | Absent | Absent | Absent | Absent | Present | Absent | Absent |
| MAG054 | Absent | Absent | Absent | Absent | Absent | Absent | Absent |
| MAG055 | Absent | Absent | Present | Present | Present | Absent | Absent |
| MAG056 | Present | Present | Absent | Absent | Absent | Absent | Absent |
| MAG057 | Absent | Absent | Present | Present | Present | Absent | Absent |
| MAG059 | Absent | Absent | Absent | Absent | Present | Absent | Absent |
| MAG062 | Present | Absent | Absent | Absent | Absent | Absent | Absent |
| MAG064 | Absent | Absent | Present | Present | Present | Absent | Absent |
| MAG066 | Present | Absent | Absent | Absent | Absent | Absent | Absent |
| MAG067 | Absent | Absent | Absent | Absent | Absent | Absent | Absent |
| MAG069 | Absent | Absent | Absent | Absent | Absent | Absent | Absent |
| MAG071 | Absent | Absent | Present | Present | Present | Absent | Absent |
| MAG072 | Absent | Absent | Present | Present | Present | Absent | Absent |
| MAG073 | Absent | Absent | Absent | Absent | Absent | Absent | Absent |
| MAG074 | Absent | Absent | Present | Present | Present | Absent | Absent |
| MAG079 | Absent | Absent | Absent | Absent | Absent | Absent | Absent |
| MAG080 | Absent | Absent | Absent | Absent | Absent | Absent | Absent |
| MAG081 | Absent | Absent | Present | Present | Present | Absent | Absent |
| MAG085 | Absent | Absent | Absent | Absent | Absent | Absent | Absent |
| MAG086 | Absent | Absent | Present | Present | Present | Absent | Absent |
| MAG088 | Absent | Absent | Present | Present | Present | Absent | Absent |
| MAG089 | Absent | Absent | Absent | Absent | Absent | Absent | Absent |
| MAG091 | Absent | Absent | Absent | Absent | Absent | Absent | Absent |
| MAG096 | Present | Present | Absent | Absent | Absent | Absent | Absent |
| MAG101 | Absent | Absent | Present | Present | Present | Absent | Absent |
| MAG102 | Absent | Absent | Absent | Absent | Absent | Absent | Absent |

| Function | CBB cycle - Rubisco | | Wood Ljungdahl pathway | | | Reverse TCA cycle | |
|---|---|---|---|---|---|---|---|
| MAG104 | Absent | Absent | Present | Present | Present | Absent | Absent |
| MAG106 | Absent | Absent | Absent | Absent | Absent | Absent | Absent |
| MAG107 | Absent | Absent | Present | Present | Present | Absent | Absent |
| MAG109 | Present | Present | Absent | Absent | Absent | Absent | Absent |
| MAG110 | Absent | Absent | Absent | Absent | Absent | Absent | Absent |
| MAG111 | Absent | Absent | Absent | Absent | Absent | Absent | Absent |
| MAG112 | Present | Absent | Absent | Absent | Absent | Absent | Absent |
| MAG114 | Present | Present | Absent | Absent | Absent | Absent | Absent |
| MAG115 | Absent | Absent | Absent | Absent | Absent | Absent | Absent |
| MAG118 | Absent | Absent | Absent | Absent | Absent | Absent | Absent |
| MAG120 | Present | Absent | Absent | Absent | Absent | Absent | Absent |
| MAG121 | Absent | Absent | Present | Present | Present | Absent | Absent |
| MAG124 | Absent | Absent | Present | Absent | Present | Absent | Absent |
| MAG127 | Absent | Absent | Absent | Absent | Absent | Absent | Absent |
| MAG132 | Absent | Absent | Present | Present | Present | Absent | Absent |
| MAG133 | Absent | Absent | Present | Present | Present | Absent | Absent |
| MAG134 | Absent | Absent | Absent | Absent | Absent | Absent | Absent |
| MAG137 | Absent | Absent | Present | Present | Present | Absent | Absent |
| MAG138 | Absent | Absent | Present | Present | Present | Absent | Absent |
| MAG142 | Absent | Absent | Present | Present | Present | Absent | Absent |
| MAG144 | Absent | Absent | Absent | Absent | Absent | Absent | Absent |
| MAG149 | Absent | Absent | Absent | Absent | Absent | Present | Present |
| MAG154 | Absent | Absent | Absent | Absent | Absent | Absent | Absent |
| MAG156 | Absent | Absent | Present | Present | Present | Absent | Absent |
| MAG158 | Absent | Absent | Present | Present | Present | Absent | Absent |
| MAG160 | Absent | Absent | Absent | Absent | Absent | Absent | Absent |
| MAG162 | Absent | Absent | Present | Present | Present | Absent | Absent |
| MAG163 | Absent | Absent | Present | Present | Present | Absent | Absent |
| MAG164 | Absent | Absent | Absent | Present | Present | Absent | Absent |
| MAG166 | Absent | Absent | Absent | Absent | Absent | Present | Present |
| MAG170 | Absent | Absent | Absent | Absent | Absent | Absent | Absent |
| MAG174 | Absent | Absent | Present | Absent | Present | Absent | Absent |
| MAG175 | Absent | Absent | Absent | Absent | Absent | Absent | Absent |
| MAG176 | Absent | Absent | Absent | Absent | Absent | Absent | Absent |

| Function | CBB cycle - Rubisco | | Wood Ljungdahl pathway | | | Reverse TCA cycle | |
|---|---|---|---|---|---|---|---|
| MAG181 | Absent | Absent | Present | Present | Present | Absent | Absent |
| MAG184 | Absent | Absent | Absent | Absent | Absent | Absent | Absent |
| MAG187 | Absent | Absent | Present | Present | Present | Absent | Absent |
| MAG188 | Present | Present | Absent | Absent | Present | Absent | Absent |
| MAG189 | Absent | Absent | Present | Present | Present | Absent | Absent |
| MAG190 | Absent | Absent | Absent | Absent | Absent | Absent | Absent |
| MAG191 | Absent | Absent | Present | Present | Present | Absent | Absent |
| MAG196 | Absent | Absent | Present | Present | Present | Absent | Absent |
| MAG197 | Absent | Absent | Absent | Absent | Absent | Absent | Absent |
| MAG198 | Present | Present | Absent | Absent | Absent | Absent | Absent |
| MAG203 | Present | Absent | Absent | Absent | Absent | Absent | Absent |
| MAG204 | Absent | Absent | Absent | Absent | Absent | Absent | Absent |
| MAG205 | Absent | Absent | Absent | Absent | Absent | Absent | Absent |
| MAG206 | Absent | Absent | Absent | Absent | Absent | Absent | Absent |
| MAG207 | Absent | Absent | Absent | Absent | Absent | Absent | Absent |
| MAG217 | Present | Absent | Absent | Absent | Absent | Absent | Absent |
| MAG220 | Absent | Absent | Present | Present | Present | Absent | Absent |
| MAG221 | Absent | Absent | Present | Present | Present | Absent | Absent |
| MAG222 | Absent | Absent | Present | Present | Present | Absent | Absent |
| MAG223 | Absent | Absent | Absent | Present | Present | Absent | Absent |
| MAG225 | Absent | Absent | Present | Present | Present | Absent | Absent |
| MAG230 | Absent | Absent | Present | Present | Present | Absent | Absent |
| MAG231 | Absent | Absent | Absent | Absent | Absent | Absent | Absent |
| MAG232 | Absent | Absent | Present | Present | Present | Absent | Absent |
| MAG237 | Absent | Absent | Absent | Absent | Absent | Absent | Absent |
| MAG238 | Absent | Absent | Absent | Absent | Absent | Absent | Absent |
| MAG239 | Absent | Absent | Absent | Absent | Absent | Absent | Absent |
| MAG240 | Present | Absent | Absent | Absent | Absent | Absent | Absent |
| MAG242 | Absent | Absent | Present | Present | Present | Absent | Absent |
| MAG247 | Absent | Absent | Present | Present | Present | Absent | Absent |
| MAG249 | Present | Absent | Absent | Absent | Absent | Absent | Absent |
| MAG252 | Present | Absent | Absent | Absent | Absent | Absent | Absent |
| MAG253 | Absent | Absent | Present | Present | Absent | Absent | Absent |
| MAG257 | Present | Absent | Absent | Absent | Absent | Absent | Absent |

| Function | CBB cycle - Rubisco | | Wood Ljungdahl pathway | | | Reverse TCA cycle | |
|---|---|---|---|---|---|---|---|
| MAG261 | Present | Absent | Absent | Absent | Absent | Absent | Absent |
| MAG271 | Absent | Absent | Absent | Absent | Absent | Absent | Absent |
| MAG273 | Absent | Absent | Present | Present | Present | Absent | Absent |
| MAG277 | Present | Present | Absent | Absent | Absent | Absent | Absent |
| MAG279 | Absent | Absent | Present | Present | Present | Absent | Absent |
| MAG282 | Present | Absent | Absent | Absent | Absent | Absent | Absent |
| MAG284 | Absent | Absent | Absent | Absent | Absent | Absent | Absent |
| MAG285 | Present | Present | Absent | Absent | Absent | Absent | Absent |
| MAG286 | Absent | Absent | Present | Present | Present | Absent | Absent |
| MAG291 | Absent | Absent | Absent | Absent | Absent | Present | Present |
| MAG292 | Absent | Absent | Absent | Absent | Absent | Absent | Absent |
| MAG293 | Present | Absent | Absent | Absent | Absent | Absent | Absent |
| MAG294 | Absent | Absent | Absent | Absent | Absent | Absent | Absent |
| MAG301 | Absent | Absent | Absent | Absent | Absent | Absent | Absent |
| MAG303 | Absent | Absent | Present | Present | Present | Absent | Absent |
| MAG306 | Absent | Absent | Absent | Absent | Absent | Absent | Absent |
| MAG307 | Present | Absent | Absent | Absent | Absent | Absent | Absent |
| MAG309 | Present | Present | Absent | Absent | Absent | Absent | Absent |
| MAG310 | Absent | Absent | Present | Present | Present | Absent | Absent |
| MAG311 | Absent | Present | Present | Present | Present | Absent | Absent |
| MAG312 | Absent | Absent | Absent | Absent | Absent | Absent | Absent |
| MAG318 | Absent | Absent | Present | Absent | Present | Absent | Absent |
| MAG320 | Absent | Absent | Absent | Absent | Absent | Absent | Absent |
| MAG321 | Absent | Absent | Absent | Present | Absent | Absent | Absent |
| MAG325 | Present | Present | Absent | Absent | Absent | Absent | Absent |
| MAG328 | Absent | Absent | Present | Present | Present | Absent | Absent |
| MAG330 | Absent | Present | Absent | Absent | Absent | Absent | Absent |

### Table S5. Presence/absence of 3-Hydroxypropionate/4-Hydroxybutyrate

| Gene-Description | 4-hydroxybutyrate---CoA ligase (AMP-forming) [EC:6.2.1.40] [RN:R09279] | 4-hydroxybutyryl-CoA dehydratase / vinylacetyl-CoA-Delta-isomerase [EC:4.2.1.120 5.3.3.3] [RN:R10782] | 3-hydroxybutyryl-CoA dehydratase / 3-hydroxyacyl-CoA dehydrogenase [EC:4.2.1.17 1.1.1.35] [RN:R03026 R01975] | 3-hydroxypropionyl-coenzyme A synthetase [EC:6.2.1.36] [RN:R09286] | 3-hydroxypropionyl-coenzyme A dehydratase [EC:4.2.1.116] [RN:R03045] | 3-hydroxypropionate dehydrogenase (NADP+) [EC:1.1.1.298] [RN:R09289] | 4-hydroxybutyrate---CoA ligase (AMP-forming) [EC:6.2.1.40] [RN:R09279] |
|---|---|---|---|---|---|---|---|
| MAG028 | Absent | Present | Present | Absent | Present | Absent | Absent |
| MAG045 | Absent | Present | Present | Absent | Present | Absent | Absent |
| MAG229 | Absent | Present | Present | Absent | Present | Absent | Absent |
| MAG294 | Absent | Present | Present | Absent | Present | Absent | Absent |

### Table S6. Presence/absence of Hydrogenases

| Function | FeFe hydrogenase | | | | | Ni-Fe Hydrogenase | | | |
|---|---|---|---|---|---|---|---|---|---|
| Gene | fefe-group-a13 | fefe-group-a2 | fefe-group-a4 | fefe-group-b | fefe-group-c1 | nife-group-1 | nife-group-2ade | nife-group-2bc | nife-group-3abd |
| MAG001 | Absent | Absent | Absent | Absent | Absent | Absent | Absent | Absent | Absent |
| MAG002 | Absent | Absent | Absent | Absent | Absent | Present | Absent | Absent | Present |
| MAG003 | Absent | Absent | Absent | Absent | Absent | Absent | Absent | Absent | Absent |
| MAG005 | Absent | Absent | Absent | Absent | Absent | Absent | Absent | Absent | Absent |
| MAG006 | Absent | Absent | Absent | Absent | Absent | Absent | Absent | Absent | Absent |
| MAG008 | Absent | Absent | Absent | Absent | Absent | Absent | Absent | Absent | Absent |
| MAG009 | Absent | Absent | Absent | Absent | Absent | Absent | Absent | Absent | Present |
| MAG011 | Absent | Absent | Absent | Absent | Absent | Absent | Absent | Absent | Absent |
| MAG012 | Present | Absent | Present | Present | Absent | Absent | Absent | Absent | Absent |
| MAG013 | Present | Absent | Present | Absent | Absent | Present | Absent | Absent | Absent |
| MAG014 | Absent | Absent | Absent | Absent | Absent | Present | Absent | Absent | Absent |
| MAG018 | Absent | Absent | Absent | Absent | Absent | Absent | Absent | Absent | Absent |
| MAG019 | Absent | Absent | Absent | Absent | Absent | Absent | Absent | Absent | Present |
| MAG020 | Present | Present | Present | Present | Present | Absent | Absent | Absent | Absent |
| MAG025 | Absent | Absent | Absent | Absent | Absent | Present | Absent | Absent | Absent |
| MAG026 | Absent | Absent | Absent | Absent | Absent | Absent | Absent | Absent | Absent |
| MAG027 | Present | Absent | Absent | Absent | Absent | Absent | Absent | Absent | Absent |
| MAG028 | Absent | Absent | Absent | Absent | Absent | Absent | Absent | Absent | Absent |
| MAG030 | Absent | Absent | Absent | Absent | Absent | Present | Absent | Absent | Absent |
| MAG037 | Absent | Absent | Absent | Absent | Absent | Absent | Absent | Absent | Absent |
| MAG038 | Absent | Absent | Absent | Absent | Absent | Absent | Absent | Absent | Absent |
| MAG042 | Absent | Absent | Absent | Absent | Absent | Present | Absent | Absent | Absent |
| MAG044 | Absent | Absent | Absent | Absent | Absent | Present | Absent | Absent | Present |
| MAG045 | Absent | Absent | Absent | Absent | Absent | Absent | Absent | Absent | Absent |
| MAG046 | Absent | Absent | Absent | Absent | Absent | Present | Absent | Absent | Present |
| MAG051 | Absent | Absent | Absent | Absent | Absent | Present | Absent | Absent | Absent |

| Function | FeFe hydrogenase | | | | | Ni-Fe Hydrogenase | | | |
|---|---|---|---|---|---|---|---|---|---|
| Gene | fefe-group-a13 | fefe-group-a2 | fefe-group-a4 | fefe-group-b | fefe-group-c1 | nife-group-1 | nife-group-2ade | nife-group-2bc | nife-group-3abd |
| MAG052 | Absent | Absent | Absent | Absent | Absent | Absent | Absent | Absent | Absent |
| MAG053 | Present | Absent | Absent | Absent | Present | Absent | Absent | Absent | Absent |
| MAG054 | Present | Absent | Absent | Present | Absent | Absent | Absent | Absent | Absent |
| MAG055 | Present | Absent | Absent | Absent | Present | Absent | Absent | Absent | Absent |
| MAG056 | Absent | Absent | Absent | Absent | Absent | Absent | Absent | Absent | Absent |
| MAG057 | Present | Absent | Absent | Absent | Absent | Present | Absent | Absent | Absent |
| MAG059 | Absent | Absent | Absent | Absent | Present | Absent | Absent | Absent | Absent |
| MAG062 | Absent | Absent | Absent | Absent | Absent | Present | Absent | Absent | Absent |
| MAG064 | Absent | Absent | Absent | Absent | Absent | Absent | Absent | Absent | Absent |
| MAG066 | Absent | Absent | Absent | Absent | Absent | Absent | Absent | Absent | Absent |
| MAG067 | Absent | Absent | Absent | Absent | Absent | Present | Absent | Absent | Absent |
| MAG069 | Present | Absent | Absent | Present | Absent | Absent | Absent | Absent | Absent |
| MAG071 | Absent | Absent | Absent | Absent | Absent | Absent | Absent | Absent | Absent |
| MAG072 | Absent | Absent | Absent | Absent | Absent | Absent | Absent | Absent | Absent |
| MAG073 | Absent | Absent | Absent | Absent | Absent | Absent | Absent | Absent | Absent |
| MAG074 | Present | Absent | Absent | Absent | Present | Present | Absent | Absent | Absent |
| MAG079 | Absent | Absent | Absent | Absent | Absent | Absent | Absent | Absent | Absent |
| MAG080 | Present | Absent | Absent | Absent | Absent | Absent | Absent | Absent | Absent |
| MAG081 | Present | Absent | Present | Present | Absent | Present | Absent | Absent | Absent |
| MAG085 | Absent | Absent | Absent | Absent | Absent | Absent | Absent | Absent | Absent |
| MAG086 | Present | Absent | Absent | Absent | Present | Present | Absent | Absent | Absent |
| MAG088 | Absent | Absent | Absent | Absent | Absent | Present | Absent | Absent | Present |
| MAG089 | Absent | Absent | Absent | Absent | Absent | Absent | Absent | Absent | Absent |
| MAG091 | Absent | Absent | Absent | Absent | Absent | Absent | Absent | Absent | Absent |
| MAG096 | Absent | Absent | Absent | Absent | Absent | Absent | Absent | Absent | Absent |
| MAG101 | Absent | Absent | Absent | Absent | Absent | Present | Absent | Absent | Present |
| MAG102 | Present | Absent | Absent | Absent | Absent | Present | Absent | Absent | Present |
| MAG104 | Present | Absent | Absent | Absent | Present | Absent | Absent | Absent | Absent |
| MAG106 | Absent | Absent | Absent | Absent | Absent | Absent | Absent | Absent | Absent |
| MAG107 | Absent | Absent | Absent | Absent | Absent | Absent | Absent | Absent | Present |
| MAG109 | Absent | Absent | Absent | Absent | Absent | Absent | Absent | Absent | Absent |
| MAG110 | Absent | Absent | Absent | Absent | Absent | Absent | Absent | Absent | Absent |
| MAG111 | Absent | Absent | Absent | Absent | Absent | Absent | Absent | Absent | Absent |
| MAG112 | Absent | Absent | Absent | Absent | Absent | Absent | Absent | Absent | Absent |
| MAG114 | Absent | Absent | Absent | Absent | Absent | Present | Absent | Absent | Absent |
| MAG115 | Absent | Absent | Absent | Absent | Absent | Absent | Absent | Absent | Absent |
| MAG118 | Absent | Absent | Absent | Absent | Absent | Absent | Absent | Absent | Absent |
| MAG120 | Absent | Absent | Absent | Absent | Absent | Absent | Present | Absent | Absent |
| MAG121 | Absent | Absent | Absent | Absent | Absent | Present | Absent | Absent | Absent |
| MAG124 | Absent | Absent | Absent | Absent | Absent | Present | Absent | Absent | Absent |
| MAG127 | Absent | Absent | Absent | Absent | Absent | Absent | Absent | Absent | Present |
| MAG132 | Absent | Absent | Absent | Absent | Absent | Present | Absent | Absent | Absent |
| MAG133 | Absent | Absent | Absent | Absent | Absent | Absent | Absent | Absent | Absent |

| Function | FeFe hydrogenase | | | | | Ni-Fe Hydrogenase | | | |
|---|---|---|---|---|---|---|---|---|---|
| Gene | fefe-group-a13 | fefe-group-a2 | fefe-group-a4 | fefe-group-b | fefe-group-c1 | nife-group-1 | nife-group-2ade | nife-group-2bc | nife-group-3abd |
| MAG134 | Present | Absent | Absent | Absent | Absent | Present | Absent | Absent | Present |
| MAG137 | Present | Absent | Present | Absent | Present | Absent | Absent | Absent | Absent |
| MAG138 | Absent | Absent | Absent | Absent | Absent | Present | Absent | Absent | Absent |
| MAG142 | Absent | Absent | Absent | Absent | Absent | Present | Absent | Absent | Absent |
| MAG144 | Absent | Absent | Absent | Absent | Absent | Absent | Absent | Absent | Absent |
| MAG149 | Absent | Absent | Absent | Absent | Absent | Present | Absent | Present | Absent |
| MAG154 | Absent | Absent | Absent | Absent | Absent | Absent | Absent | Absent | Absent |
| MAG156 | Absent | Absent | Absent | Absent | Absent | Present | Absent | Absent | Absent |
| MAG158 | Present | Absent | Absent | Absent | Present | Present | Absent | Absent | Absent |
| MAG160 | Absent | Absent | Absent | Absent | Absent | Absent | Absent | Absent | Present |
| MAG162 | Absent | Absent | Absent | Absent | Absent | Present | Absent | Absent | Present |
| MAG163 | Absent | Absent | Absent | Absent | Absent | Present | Absent | Absent | Present |
| MAG164 | Present | Absent | Absent | Absent | Absent | Absent | Absent | Absent | Absent |
| MAG166 | Absent | Absent | Absent | Absent | Absent | Absent | Absent | Absent | Present |
| MAG170 | Present | Absent | Absent | Absent | Absent | Absent | Absent | Absent | Absent |
| MAG174 | Absent | Absent | Absent | Absent | Absent | Present | Absent | Absent | Absent |
| MAG175 | Absent | Absent | Absent | Absent | Absent | Absent | Absent | Absent | Present |
| MAG176 | Absent | Absent | Absent | Absent | Absent | Present | Absent | Absent | Present |
| MAG181 | Absent | Absent | Absent | Absent | Absent | Present | Absent | Absent | Absent |
| MAG184 | Absent | Absent | Absent | Absent | Absent | Present | Absent | Absent | Absent |
| MAG187 | Present | Absent | Absent | Absent | Present | Absent | Absent | Absent | Absent |
| MAG188 | Present | Absent | Present | Present | Absent | Present | Absent | Absent | Absent |
| MAG189 | Absent | Absent | Absent | Absent | Absent | Absent | Absent | Absent | Absent |
| MAG190 | Absent | Absent | Absent | Absent | Absent | Present | Absent | Absent | Absent |
| MAG191 | Absent | Absent | Absent | Absent | Absent | Absent | Absent | Absent | Absent |
| MAG196 | Absent | Absent | Absent | Absent | Absent | Present | Absent | Absent | Absent |
| MAG197 | Absent | Absent | Absent | Absent | Absent | Present | Absent | Absent | Absent |
| MAG198 | Absent | Absent | Absent | Absent | Absent | Present | Absent | Present | Absent |
| MAG203 | Absent | Absent | Absent | Absent | Absent | Absent | Absent | Absent | Absent |
| MAG204 | Absent | Absent | Absent | Absent | Absent | Absent | Absent | Absent | Absent |
| MAG205 | Absent | Absent | Absent | Absent | Absent | Absent | Absent | Absent | Absent |
| MAG206 | Present | Absent | Absent | Present | Present | Absent | Absent | Absent | Absent |
| MAG207 | Absent | Absent | Absent | Absent | Absent | Absent | Absent | Absent | Absent |
| MAG217 | Absent | Absent | Absent | Absent | Absent | Present | Absent | Present | Absent |
| MAG220 | Absent | Absent | Absent | Absent | Absent | Present | Absent | Absent | Present |
| MAG221 | Absent | Absent | Absent | Absent | Absent | Present | Absent | Absent | Absent |
| MAG222 | Absent | Absent | Present | Absent | Absent | Absent | Absent | Absent | Absent |
| MAG223 | Absent | Absent | Absent | Absent | Absent | Present | Absent | Absent | Absent |
| MAG225 | Present | Absent | Present | Absent | Present | Absent | Absent | Absent | Absent |
| MAG230 | Absent | Absent | Absent | Absent | Absent | Absent | Absent | Absent | Absent |
| MAG231 | Present | Absent | Absent | Absent | Absent | Absent | Absent | Absent | Absent |
| MAG232 | Absent | Absent | Absent | Absent | Absent | Absent | Absent | Absent | Absent |
| MAG237 | Present | Absent | Absent | Absent | Absent | Absent | Absent | Absent | Absent |

| Function | FeFe hydrogenase | | | | | Ni-Fe Hydrogenase | | | |
|---|---|---|---|---|---|---|---|---|---|
| Gene | fefe-group-a13 | fefe-group-a2 | fefe-group-a4 | fefe-group-b | fefe-group-c1 | nife-group-1 | nife-group-2ade | nife-group-2bc | nife-group-3abd |
| MAG238 | Absent | Absent | Absent | Absent | Absent | Absent | Absent | Absent | Present |
| MAG239 | Present | Absent | Absent | Absent | Absent | Absent | Absent | Absent | Absent |
| MAG240 | Absent | Absent | Absent | Absent | Absent | Present | Absent | Present | Absent |
| MAG242 | Absent | Absent | Absent | Absent | Absent | Present | Absent | Absent | Absent |
| MAG247 | Absent | Absent | Absent | Absent | Absent | Present | Absent | Absent | Absent |
| MAG249 | Absent | Absent | Absent | Absent | Absent | Present | Absent | Absent | Absent |
| MAG252 | Absent | Absent | Absent | Absent | Absent | Present | Absent | Present | Absent |
| MAG253 | Present | Absent | Absent | Absent | Present | Present | Absent | Absent | Absent |
| MAG257 | Absent | Absent | Absent | Absent | Absent | Present | Absent | Present | Present |
| MAG261 | Absent | Absent | Absent | Absent | Absent | Present | Absent | Present | Absent |
| MAG271 | Absent | Absent | Absent | Absent | Absent | Present | Absent | Absent | Absent |
| MAG273 | Absent | Absent | Absent | Absent | Absent | Present | Absent | Absent | Absent |
| MAG277 | Absent | Absent | Absent | Absent | Absent | Absent | Absent | Absent | Absent |
| MAG279 | Absent | Absent | Absent | Absent | Absent | Present | Absent | Absent | Absent |
| MAG282 | Absent | Absent | Absent | Absent | Absent | Absent | Absent | Absent | Absent |
| MAG284 | Absent | Absent | Absent | Absent | Absent | Absent | Absent | Absent | Absent |
| MAG285 | Absent | Absent | Absent | Absent | Absent | Absent | Absent | Absent | Absent |
| MAG286 | Present | Absent | Present | Absent | Present | Absent | Absent | Absent | Absent |
| MAG291 | Absent | Absent | Absent | Absent | Absent | Present | Absent | Absent | Absent |
| MAG292 | Present | Absent | Absent | Absent | Absent | Absent | Absent | Absent | Absent |
| MAG293 | Absent | Absent | Absent | Absent | Absent | Absent | Absent | Absent | Absent |
| MAG294 | Absent | Absent | Absent | Absent | Absent | Absent | Absent | Absent | Absent |
| MAG301 | Present | Absent | Absent | Absent | Present | Absent | Absent | Absent | Absent |
| MAG303 | Present | Absent | Absent | Absent | Absent | Present | Absent | Absent | Absent |
| MAG306 | Absent | Absent | Absent | Absent | Absent | Absent | Absent | Absent | Absent |
| MAG307 | Absent | Absent | Absent | Absent | Absent | Absent | Present | Present | Absent |
| MAG309 | Absent | Absent | Absent | Absent | Absent | Present | Absent | Present | Present |
| MAG310 | Absent | Absent | Absent | Absent | Absent | Present | Absent | Absent | Absent |
| MAG311 | Absent | Absent | Absent | Absent | Absent | Present | Absent | Absent | Absent |
| MAG312 | Absent | Absent | Absent | Absent | Absent | Present | Absent | Absent | Absent |
| MAG318 | Absent | Absent | Absent | Absent | Absent | Present | Absent | Absent | Present |
| MAG320 | Absent | Absent | Absent | Absent | Absent | Absent | Absent | Absent | Absent |
| MAG321 | Absent | Absent | Absent | Absent | Absent | Absent | Absent | Absent | Absent |
| MAG325 | Absent | Absent | Absent | Absent | Absent | Present | Absent | Present | Present |
| MAG328 | Present | Absent | Absent | Absent | Absent | Absent | Absent | Absent | Absent |
| MAG330 | Absent | Absent | Absent | Absent | Absent | Present | Absent | Absent | Absent |

### Table S7. Number of hydrogenases present in MAGs

| Function | uptake hydrogenase small subunit [EC:1.12.99.6] | uptake hydrogenase large subunit [EC:1.12.99.6] | $F_{420}$-non-reducing hydrogenase large subunit [EC:1.12.99.-1.8.98.5] | $F^{420}$-non-reducing hydrogenase iron-sulfur subunit [EC:1.12.99.-1.8.98.5 1.8.98.6] | $F_{420}$-non-reducing hydrogenase small subunit [EC:1.12.99.-1.8.98.5] | NADP-reducing hydrogenase subunit [EC:1.12.1.3] | [NiFe] hydrogenase small subunit [EC:1.12.2.1] | [NiFe] hydrogenase large subunit [EC:1.12.2.1] | [NiFe] hydrogenase diaphorase moiety large subunit [EC:1.12.1.2] | [NiFe] hydrogenase diaphorase moiety small subunit [EC:1.12.1.2] | NAD-reducing hydrogenase small subunit [EC:1.12.1.2] |
|---|---|---|---|---|---|---|---|---|---|---|---|
| Gene | | *hupV* | *mvhA* | *mvhD* | *mvhG* | *hndB* | *hydA* | *hydB* | *hoxF* | *hoXU* | *hoxY* |
| MAG001 | 0 | 0 | 0 | 0 | 0 | 0 | 0 | 0 | 0 | 0 | 0 |
| MAG002 | 0 | 0 | 2 | 2 | 2 | 0 | 0 | 0 | 0 | 0 | 0 |
| MAG006 | 0 | 0 | 0 | 0 | 0 | 0 | 0 | 0 | 0 | 0 | 0 |
| MAG009 | 0 | 0 | 0 | 0 | 0 | 0 | 0 | 0 | 0 | 0 | 0 |
| MAG011 | 0 | 0 | 0 | 0 | 0 | 0 | 0 | 0 | 0 | 0 | 0 |
| MAG012 | 0 | 0 | 0 | 0 | 0 | 0 | 0 | 0 | 0 | 0 | 0 |
| MAG013 | 0 | 0 | 0 | 3 | 1 | 0 | 0 | 0 | 0 | 0 | 0 |
| MAG018 | 0 | 0 | 0 | 0 | 0 | 0 | 0 | 0 | 0 | 0 | 0 |
| MAG019 | 0 | 0 | 0 | 0 | 0 | 0 | 0 | 0 | 0 | 0 | 0 |
| MAG020 | 0 | 0 | 0 | 0 | 0 | 2 | 0 | 0 | 0 | 0 | 0 |
| MAG025 | 0 | 0 | 0 | 0 | 0 | 1 | 0 | 0 | 0 | 0 | 0 |
| MAG027 | 0 | 0 | 0 | 0 | 0 | 0 | 0 | 0 | 1 | 0 | 0 |
| MAG028 | 0 | 0 | 0 | 0 | 0 | 0 | 0 | 0 | 0 | 0 | 0 |
| MAG037 | 0 | 0 | 0 | 0 | 0 | 0 | 0 | 0 | 0 | 0 | 0 |
| MAG038 | 0 | 0 | 0 | 0 | 0 | 0 | 0 | 0 | 0 | 0 | 0 |
| MAG042 | 1 | 1 | 0 | 0 | 0 | 1 | 0 | 0 | 0 | 0 | 0 |
| MAG044 | 0 | 0 | 2 | 2 | 2 | 0 | 0 | 0 | 0 | 0 | 0 |
| MAG045 | 0 | 0 | 0 | 0 | 0 | 0 | 0 | 0 | 0 | 0 | 0 |
| MAG046 | 0 | 0 | 0 | 0 | 0 | 0 | 0 | 0 | 1 | 1 | 1 |
| MAG051 | 0 | 0 | 0 | 0 | 0 | 0 | 0 | 0 | 0 | 0 | 0 |
| MAG055 | 0 | 0 | 0 | 4 | 0 | 1 | 0 | 0 | 0 | 0 | 0 |
| MAG056 | 0 | 0 | 0 | 0 | 0 | 0 | 0 | 0 | 0 | 0 | 0 |
| MAG057 | 0 | 0 | 0 | 2 | 1 | 0 | 0 | 0 | 0 | 0 | 0 |
| MAG059 | 0 | 0 | 1 | 0 | 1 | 1 | 0 | 0 | 0 | 0 | 0 |
| MAG062 | 0 | 0 | 0 | 0 | 0 | 0 | 0 | 0 | 0 | 0 | 0 |
| MAG064 | 0 | 0 | 1 | 2 | 1 | 1 | 0 | 0 | 0 | 0 | 0 |
| MAG066 | 0 | 0 | 0 | 0 | 0 | 0 | 0 | 0 | 0 | 0 | 0 |
| MAG069 | 0 | 0 | 0 | 0 | 0 | 0 | 0 | 0 | 1 | 0 | 0 |
| MAG071 | 0 | 0 | 1 | 2 | 1 | 0 | 0 | 0 | 0 | 0 | 0 |
| MAG072 | 0 | 0 | 1 | 5 | 1 | 1 | 0 | 0 | 0 | 0 | 0 |
| MAG074 | 0 | 0 | 0 | 1 | 1 | 1 | 0 | 0 | 0 | 0 | 0 |
| MAG079 | 0 | 0 | 0 | 0 | 0 | 0 | 0 | 0 | 0 | 0 | 0 |
| MAG081 | 0 | 0 | 0 | 1 | 1 | 0 | 0 | 0 | 0 | 0 | 0 |
| MAG086 | 0 | 0 | 0 | 3 | 0 | 1 | 0 | 0 | 0 | 0 | 0 |

| | | | | | | | | | | | |
|---|---|---|---|---|---|---|---|---|---|---|---|
| MAG088 | 0 | 0 | 1 | 1 | 1 | 0 | 0 | 0 | 0 | 0 | 0 |
| MAG091 | 0 | 0 | 0 | 0 | 0 | 0 | 0 | 0 | 0 | 0 | 0 |
| MAG096 | 0 | 0 | 0 | 0 | 0 | 0 | 0 | 0 | 0 | 0 | 0 |
| MAG101 | 0 | 0 | 0 | 3 | 0 | 1 | 1 | 0 | 0 | 0 | 1 |
| MAG102 | 0 | 0 | 0 | 0 | 0 | 0 | 0 | 0 | 1 | 1 | 1 |
| MAG104 | 0 | 0 | 1 | 3 | 1 | 1 | 0 | 0 | 0 | 0 | 0 |
| MAG106 | 0 | 0 | 0 | 0 | 0 | 0 | 0 | 0 | 0 | 0 | 0 |
| MAG107 | 0 | 0 | 1 | 6 | 1 | 0 | 0 | 0 | 0 | 0 | 0 |
| MAG109 | 0 | 0 | 0 | 0 | 0 | 0 | 0 | 0 | 0 | 0 | 0 |
| MAG110 | 0 | 0 | 0 | 0 | 0 | 0 | 0 | 0 | 0 | 0 | 0 |
| MAG111 | 0 | 0 | 0 | 0 | 0 | 0 | 0 | 0 | 0 | 0 | 0 |
| MAG112 | 0 | 0 | 0 | 0 | 0 | 0 | 0 | 0 | 0 | 0 | 0 |
| MAG114 | 0 | 0 | 0 | 0 | 0 | 0 | 0 | 0 | 0 | 0 | 0 |
| MAG115 | 0 | 0 | 0 | 0 | 0 | 0 | 0 | 0 | 0 | 0 | 0 |
| MAG118 | 0 | 0 | 0 | 0 | 0 | 0 | 0 | 0 | 0 | 0 | 0 |
| MAG120 | 1 | 1 | 0 | 0 | 0 | 0 | 0 | 0 | 0 | 0 | 0 |
| MAG121 | 0 | 0 | 0 | 3 | 0 | 0 | 0 | 0 | 0 | 0 | 0 |
| MAG124 | 0 | 0 | 1 | 4 | 1 | 0 | 0 | 0 | 0 | 0 | 0 |
| MAG127 | 0 | 0 | 0 | 0 | 0 | 0 | 0 | 0 | 1 | 1 | 1 |
| MAG132 | 0 | 0 | 0 | 1 | 0 | 0 | 0 | 0 | 0 | 0 | 0 |
| MAG133 | 0 | 0 | 1 | 1 | 1 | 0 | 0 | 0 | 0 | 0 | 0 |
| MAG137 | 0 | 0 | 0 | 3 | 0 | 1 | 0 | 0 | 0 | 0 | 0 |
| MAG144 | 0 | 0 | 0 | 0 | 0 | 0 | 0 | 0 | 0 | 0 | 0 |
| MAG149 | 1 | 1 | 0 | 0 | 0 | 0 | 0 | 0 | 0 | 0 | 0 |
| MAG154 | 0 | 0 | 0 | 0 | 0 | 0 | 0 | 0 | 0 | 0 | 0 |
| MAG156 | 0 | 0 | 0 | 8 | 0 | 0 | 1 | 1 | 0 | 0 | 0 |
| MAG158 | 0 | 0 | 1 | 2 | 1 | 1 | 0 | 0 | 0 | 0 | 0 |
| MAG160 | 0 | 0 | 0 | 0 | 0 | 0 | 0 | 0 | 1 | 1 | 1 |
| MAG162 | 0 | 0 | 2 | 2 | 2 | 0 | 0 | 0 | 0 | 0 | 0 |
| MAG170 | 0 | 0 | 0 | 0 | 0 | 0 | 0 | 0 | 1 | 0 | 0 |
| MAG174 | 0 | 0 | 1 | 2 | 2 | 0 | 0 | 0 | 0 | 0 | 0 |
| MAG176 | 0 | 0 | 0 | 0 | 0 | 0 | 0 | 0 | 0 | 0 | 0 |
| MAG187 | 0 | 0 | 0 | 1 | 0 | 1 | 0 | 0 | 0 | 0 | 0 |
| MAG188 | 0 | 0 | 0 | 0 | 0 | 0 | 0 | 0 | 0 | 0 | 0 |
| MAG190 | 0 | 0 | 0 | 0 | 0 | 0 | 0 | 0 | 0 | 0 | 0 |
| MAG196 | 0 | 0 | 1 | 1 | 1 | 1 | 0 | 0 | 0 | 0 | 0 |
| MAG197 | 0 | 0 | 0 | 0 | 0 | 0 | 0 | 0 | 0 | 0 | 0 |
| MAG198 | 1 | 1 | 0 | 0 | 0 | 0 | 0 | 0 | 0 | 0 | 0 |
| MAG203 | 0 | 0 | 0 | 0 | 0 | 0 | 0 | 0 | 0 | 0 | 0 |
| MAG204 | 0 | 0 | 0 | 0 | 0 | 0 | 0 | 0 | 0 | 0 | 0 |
| MAG217 | 1 | 1 | 0 | 0 | 0 | 0 | 0 | 0 | 0 | 0 | 0 |
| MAG220 | 0 | 0 | 0 | 6 | 1 | 1 | 1 | 1 | 0 | 0 | 1 |
| MAG221 | 1 | 1 | 1 | 1 | 1 | 0 | 0 | 0 | 0 | 0 | 0 |
| MAG223 | 0 | 0 | 0 | 0 | 0 | 0 | 0 | 0 | 0 | 0 | 0 |
| MAG225 | 0 | 0 | 1 | 3 | 1 | 1 | 0 | 0 | 0 | 0 | 0 |
| MAG230 | 0 | 0 | 0 | 4 | 0 | 0 | 0 | 0 | 0 | 0 | 0 |
| MAG232 | 0 | 0 | 0 | 1 | 0 | 1 | 0 | 0 | 0 | 0 | 0 |

| | | | | | | | | | | | |
|---|---|---|---|---|---|---|---|---|---|---|---|
| MAG238 | 0 | 0 | 1 | 1 | 1 | 0 | 0 | 0 | 0 | 0 | 0 |
| MAG240 | 1 | 1 | 0 | 0 | 0 | 0 | 0 | 0 | 0 | 0 | 0 |
| MAG242 | 0 | 0 | 0 | 0 | 0 | 0 | 0 | 0 | 0 | 0 | 0 |
| MAG247 | 0 | 0 | 0 | 4 | 0 | 0 | 0 | 0 | 0 | 0 | 0 |
| MAG249 | 0 | 0 | 0 | 0 | 0 | 0 | 0 | 0 | 0 | 0 | 0 |
| MAG252 | 1 | 1 | 0 | 0 | 0 | 0 | 0 | 0 | 0 | 0 | 0 |
| MAG253 | 0 | 0 | 0 | 0 | 0 | 1 | 0 | 0 | 0 | 0 | 0 |
| MAG273 | 0 | 0 | 1 | 3 | 1 | 0 | 0 | 0 | 0 | 0 | 0 |
| MAG277 | 0 | 0 | 0 | 0 | 0 | 0 | 0 | 0 | 0 | 0 | 0 |
| MAG279 | 0 | 0 | 0 | 7 | 0 | 1 | 1 | 1 | 0 | 0 | 0 |
| MAG282 | 0 | 0 | 0 | 0 | 0 | 0 | 0 | 0 | 0 | 0 | 0 |
| MAG284 | 0 | 0 | 0 | 0 | 0 | 0 | 0 | 0 | 0 | 0 | 0 |
| MAG285 | 0 | 0 | 0 | 0 | 0 | 0 | 0 | 0 | 0 | 0 | 0 |
| MAG286 | 0 | 0 | 1 | 3 | 1 | 1 | 0 | 0 | 0 | 0 | 0 |
| MAG293 | 0 | 0 | 0 | 0 | 0 | 0 | 0 | 0 | 0 | 0 | 0 |
| MAG294 | 0 | 0 | 0 | 0 | 0 | 0 | 0 | 0 | 0 | 0 | 0 |
| MAG303 | 0 | 0 | 0 | 2 | 0 | 1 | 0 | 0 | 0 | 0 | 0 |
| MAG306 | 0 | 0 | 0 | 0 | 0 | 0 | 0 | 0 | 0 | 0 | 0 |
| MAG307 | 2 | 2 | 0 | 0 | 0 | 0 | 0 | 0 | 0 | 0 | 0 |
| MAG318 | 1 | 1 | 1 | 7 | 1 | 1 | 1 | 0 | 0 | 0 | 1 |
| MAG321 | 0 | 0 | 2 | 1 | 2 | 1 | 0 | 0 | 0 | 0 | 0 |
| MAG325 | 1 | 1 | 0 | 0 | 0 | 0 | 0 | 0 | 1 | 1 | 1 |
| MAG330 | 1 | 0 | 0 | 0 | 0 | 0 | 0 | 0 | 0 | 0 | 0 |

### Table S8. Presence/ absence of nitrogen reduction pathway

| Function | Nitrate reduction | | | | Nitrite reduction to ammonia | | | Nitrite reduction | | Nitric oxide reduction | | |
|---|---|---|---|---|---|---|---|---|---|---|---|---|
| Gene | napA | napB | narG | narH | nrfA | nirB | nirD | nirK | nirS | norB | norC | nosZ |
| MAG001 | Absent | Absent | Present | Present | Absent | Absent | Absent | Absent | Absent | Absent | Absent | Present |
| MAG002 | Absent | Absent | Absent | Absent | Absent | Absent | Absent | Absent | Absent | Absent | Absent | Absent |
| MAG003 | Absent | Absent | Absent | Absent | Absent | Absent | Absent | Absent | Absent | Absent | Absent | Absent |
| MAG005 | Absent | Absent | Absent | Absent | Present | Absent | Absent | Absent | Absent | Absent | Absent | Absent |
| MAG006 | Absent | Absent | Present | Present | Absent | Present | Present | Absent | Absent | Absent | Absent | Absent |
| MAG008 | Absent | Absent | Present | Present | Absent | Absent | Absent | Absent | Absent | Present | Absent | Absent |
| MAG009 | Absent | Absent | Absent | Absent | Absent | Present | Present | Absent | Absent | Absent | Absent | Absent |
| MAG011 | Absent | Absent | Absent | Absent | Absent | Present | Present | Absent | Absent | Absent | Absent | Absent |
| MAG012 | Absent | Absent | Absent | Absent | Present | Absent | Absent | Absent | Absent | Absent | Absent | Absent |
| MAG013 | Absent | Absent | Absent | Absent | Absent | Absent | Absent | Absent | Absent | Absent | Absent | Absent |
| MAG014 | Absent | Absent | Absent | Absent | Present | Absent | Absent | Absent | Absent | Present | Absent | Absent |
| MAG018 | Absent | Absent | Absent | Absent | Absent | Present | Present | Absent | Absent | Absent | Absent | Absent |

| Function | Nitrate reduction | | | | Nitrite reduction to ammonia | | | Nitrite reduction | | Nitric oxide reduction | | |
|---|---|---|---|---|---|---|---|---|---|---|---|---|
| Gene | napA | napB | narG | narH | nrfA | nirB | nirD | nirK | nirS | norB | norC | nosZ |
| MAG019 | Absent | Absent | Absent | Absent | Absent | Present | Present | Absent | Absent | Absent | Absent | Absent |
| MAG020 | Absent | Absent | Absent | Absent | Absent | Absent | Absent | Absent | Absent | Absent | Absent | Absent |
| MAG025 | Absent | Absent | Absent | Absent | Present | Absent | Absent | Absent | Absent | Present | Present | Absent |
| MAG026 | Absent | Absent | Absent | Absent | Absent | Absent | Absent | Absent | Absent | Absent | Absent | Absent |
| MAG027 | Absent | Absent | Absent | Absent | Absent | Absent | Absent | Absent | Absent | Absent | Absent | Absent |
| MAG028 | Absent | Absent | Absent | Absent | Absent | Absent | Absent | Absent | Absent | Absent | Absent | Absent |
| MAG030 | Absent | Absent | Absent | Absent | Absent | Absent | Absent | Absent | Absent | Absent | Absent | Absent |
| MAG037 | Absent | Absent | Absent | Absent | Absent | Absent | Absent | Absent | Absent | Absent | Absent | Absent |
| MAG038 | Absent | Absent | Absent | Absent | Absent | Present | Present | Present | Absent | Absent | Absent | Absent |
| MAG042 | Absent | Absent | Absent | Absent | Absent | Absent | Absent | Absent | Absent | Absent | Absent | Absent |
| MAG044 | Absent | Absent | Absent | Absent | Absent | Absent | Absent | Absent | Absent | Absent | Absent | Absent |
| MAG045 | Absent | Absent | Absent | Absent | Absent | Absent | Absent | Absent | Absent | Absent | Absent | Absent |
| MAG046 | Present | Present | Absent | Absent | Present | Present | Present | Absent | Absent | Absent | Absent | Absent |
| MAG051 | Absent | Absent | Present | Present | Absent | Present | Present | Absent | Absent | Present | Absent | Absent |
| MAG052 | Absent | Absent | Absent | Absent | Absent | Absent | Absent | Present | Absent | Absent | Absent | Absent |
| MAG053 | Absent | Absent | Absent | Absent | Present | Absent | Absent | Absent | Absent | Absent | Absent | Absent |
| MAG054 | Absent | Absent | Absent | Absent | Absent | Absent | Absent | Absent | Absent | Absent | Absent | Absent |
| MAG055 | Absent | Absent | Absent | Absent | Present | Absent | Absent | Absent | Absent | Absent | Absent | Absent |
| MAG056 | Absent | Absent | Present | Present | Absent | Present | Present | Absent | Absent | Absent | Absent | Absent |
| MAG057 | Absent | Absent | Absent | Absent | Absent | Absent | Absent | Absent | Absent | Absent | Absent | Absent |
| MAG059 | Absent | Absent | Absent | Absent | Absent | Absent | Absent | Absent | Absent | Absent | Absent | Absent |
| MAG062 | Absent | Absent | Absent | Absent | Absent | Present | Present | Absent | Absent | Absent | Absent | Absent |
| MAG064 | Absent | Absent | Absent | Absent | Absent | Absent | Absent | Absent | Absent | Absent | Absent | Absent |
| MAG066 | Absent | Absent | Present | Present | Absent | Present | Present | Absent | Absent | Present | Present | Absent |
| MAG067 | Absent | Absent | Absent | Absent | Absent | Present | Absent | Absent | Present | Present | Present | Present |
| MAG069 | Absent | Absent | Absent | Absent | Absent | Absent | Absent | Absent | Absent | Absent | Absent | Absent |
| MAG071 | Absent | Absent | Absent | Absent | Present | Absent | Present | Absent | Absent | Absent | Absent | Absent |
| MAG072 | Absent | Absent | Absent | Absent | Absent | Absent | Absent | Absent | Absent | Absent | Absent | Absent |
| MAG073 | Absent | Absent | Absent | Absent | Absent | Present | Present | Absent | Absent | Absent | Absent | Absent |
| MAG074 | Absent | Absent | Absent | Absent | Absent | Absent | Absent | Absent | Absent | Absent | Absent | Present |
| MAG079 | Absent | Absent | Present | Present | Absent | Present | Present | Absent | Absent | Absent | Absent | Absent |
| MAG080 | Absent | Absent | Absent | Absent | Absent | Absent | Absent | Absent | Absent | Absent | Absent | Absent |

| Function | Nitrate reduction | | | | Nitrite reduction to ammonia | | | Nitrite reduction | | Nitric oxide reduction | | |
|---|---|---|---|---|---|---|---|---|---|---|---|---|
| Gene | napA | napB | narG | narH | nrfA | nirB | nirD | nirK | nirS | norB | norC | nosZ |
| MAG081 | Absent | Absent | Absent | Absent | Absent | Absent | Absent | Absent | Absent | Absent | Absent | Absent |
| MAG085 | Absent | Absent | Absent | Absent | Absent | Absent | Absent | Absent | Absent | Absent | Absent | Absent |
| MAG086 | Absent | Absent | Absent | Absent | Present | Absent | Absent | Absent | Absent | Absent | Absent | Absent |
| MAG088 | Absent | Absent | Absent | Absent | Absent | Absent | Absent | Absent | Absent | Absent | Absent | Absent |
| MAG089 | Absent | Absent | Absent | Absent | Present | Absent | Absent | Absent | Absent | Absent | Absent | Absent |
| MAG091 | Absent | Absent | Absent | Absent | Absent | Present | Present | Absent | Absent | Absent | Absent | Absent |
| MAG096 | Absent | Absent | Present | Present | Absent | Present | Present | Absent | Absent | Absent | Absent | Absent |
| MAG101 | Present | Absent | Absent | Absent | Present | Absent | Absent | Absent | Absent | Present | Absent | Absent |
| MAG102 | Absent | Absent | Absent | Absent | Present | Absent | Absent | Absent | Absent | Present | Absent | Absent |
| MAG104 | Absent | Absent | Absent | Absent | Present | Absent | Absent | Absent | Absent | Absent | Absent | Absent |
| MAG106 | Absent | Absent | Absent | Absent | Absent | Absent | Absent | Absent | Absent | Absent | Absent | Absent |
| MAG107 | Absent | Absent | Absent | Absent | Present | Absent | Absent | Absent | Absent | Absent | Absent | Absent |
| MAG109 | Absent | Absent | Absent | Absent | Absent | Absent | Absent | Absent | Absent | Absent | Absent | Absent |
| MAG110 | Absent | Absent | Absent | Absent | Absent | Present | Present | Absent | Absent | Absent | Absent | Absent |
| MAG111 | Absent | Absent | Absent | Absent | Absent | Present | Present | Present | Absent | Present | Absent | Absent |
| MAG112 | Absent | Absent | Absent | Absent | Absent | Present | Present | Absent | Absent | Absent | Absent | Absent |
| MAG114 | Present | Present | Present | Present | Absent | Present | Present | Absent | Present | Present | Present | Absent |
| MAG115 | Absent | Absent | Absent | Absent | Absent | Absent | Absent | Absent | Absent | Present | Absent | Absent |
| MAG118 | Absent | Absent | Absent | Absent | Absent | Absent | Absent | Absent | Absent | Absent | Absent | Absent |
| MAG120 | Absent | Absent | Absent | Absent | Absent | Present | Present | Present | Absent | Absent | Absent | Absent |
| MAG121 | Absent | Absent | Absent | Absent | Present | Absent | Absent | Absent | Absent | Absent | Absent | Absent |
| MAG124 | Absent | Absent | Absent | Absent | Absent | Absent | Absent | Absent | Absent | Absent | Absent | Absent |
| MAG127 | Absent | Absent | Absent | Absent | Present | Absent | Absent | Absent | Absent | Absent | Absent | Present |
| MAG132 | Absent | Absent | Absent | Absent | Absent | Absent | Absent | Absent | Absent | Absent | Absent | Absent |
| MAG133 | Absent | Absent | Absent | Absent | Present | Absent | Absent | Absent | Present | Absent | Absent | Absent |
| MAG134 | Absent | Absent | Absent | Absent | Absent | Absent | Absent | Absent | Absent | Absent | Absent | Absent |
| MAG137 | Absent | Absent | Absent | Absent | Present | Absent | Absent | Absent | Absent | Absent | Absent | Absent |
| MAG138 | Absent | Absent | Absent | Absent | Absent | Absent | Absent | Absent | Absent | Absent | Absent | Absent |
| MAG142 | Absent | Absent | Absent | Absent | Present | Absent | Absent | Absent | Present | Absent | Absent | Absent |
| MAG144 | Absent | Absent | Absent | Absent | Present | Present | Present | Present | Absent | Present | Absent | Present |
| MAG149 | Absent | Absent | Absent | Absent | Absent | Absent | Present | Present | Absent | Absent | Absent | Absent |
| MAG154 | Present | Present | Present | Present | Absent | Present | Present | Absent | Present | Present | Present | Absent |

| Function | Nitrate reduction | | | | Nitrite reduction to ammonia | | | Nitrite reduction | | Nitric oxide reduction | | |
|---|---|---|---|---|---|---|---|---|---|---|---|---|
| Gene | napA | napB | narG | narH | nrfA | nirB | nirD | nirK | nirS | norB | norC | nosZ |
| MAG156 | Absent | Absent | Absent | Absent | Absent | Absent | Absent | Absent | Absent | Absent | Absent | Absent |
| MAG158 | Absent | Absent | Absent | Absent | Present | Absent | Absent | Absent | Absent | Absent | Absent | Absent |
| MAG160 | Absent | Absent | Absent | Absent | Present | Absent | Absent | Absent | Absent | Absent | Absent | Present |
| MAG162 | Absent | Absent | Absent | Absent | Absent | Absent | Absent | Absent | Absent | Absent | Absent | Absent |
| MAG163 | Absent | Absent | Absent | Absent | Absent | Absent | Absent | Absent | Absent | Absent | Absent | Absent |
| MAG164 | Absent | Absent | Absent | Absent | Absent | Absent | Absent | Absent | Absent | Absent | Absent | Absent |
| MAG166 | Absent | Absent | Absent | Absent | Absent | Absent | Absent | Absent | Absent | Absent | Absent | Absent |
| MAG170 | Absent | Absent | Absent | Absent | Present | Absent | Present | Present | Absent | Absent | Absent | Absent |
| MAG174 | Absent | Absent | Absent | Absent | Absent | Absent | Absent | Absent | Absent | Absent | Absent | Absent |
| MAG175 | Absent | Absent | Absent | Absent | Present | Absent | Absent | Absent | Present | Absent | Absent | Absent |
| MAG176 | Absent | Absent | Absent | Absent | Present | Present | Present | Absent | Absent | Present | Present | Absent |
| MAG181 | Absent | Absent | Absent | Absent | Present | Absent | Absent | Absent | Present | Absent | Absent | Absent |
| MAG184 | Absent | Absent | Absent | Absent | Present | Absent | Absent | Absent | Absent | Present | Absent | Present |
| MAG187 | Absent | Absent | Absent | Absent | Absent | Absent | Absent | Absent | Absent | Absent | Absent | Absent |
| MAG188 | Absent | Absent | Absent | Absent | Absent | Absent | Absent | Absent | Absent | Absent | Absent | Absent |
| MAG189 | Absent | Absent | Absent | Absent | Absent | Absent | Absent | Absent | Absent | Absent | Absent | Absent |
| MAG190 | Absent | Absent | Absent | Absent | Absent | Absent | Absent | Absent | Absent | Absent | Absent | Present |
| MAG191 | Present | Absent | Absent | Absent | Absent | Absent | Absent | Absent | Absent | Present | Absent | Absent |
| MAG196 | Present | Absent | Absent | Absent | Present | Absent | Absent | Absent | Absent | Present | Present | Absent |
| MAG197 | Absent | Absent | Absent | Absent | Absent | Present | Present | Present | Absent | Present | Absent | Absent |
| MAG198 | Absent | Absent | Present | Present | Absent | Present | Present | Absent | Present | Present | Present | Absent |
| MAG203 | Present | Present | Absent | Absent | Absent | Present | Present | Absent | Present | Present | Present | Absent |
| MAG204 | Absent | Absent | Present | Present | Absent | Present | Present | Absent | Absent | Present | Absent | Absent |
| MAG205 | Absent | Absent | Present | Present | Absent | Present | Present | Absent | Absent | Absent | Absent | Absent |
| MAG206 | Absent | Absent | Absent | Absent | Present | Absent | Absent | Absent | Absent | Present | Absent | Absent |
| MAG207 | Absent | Absent | Absent | Absent | Absent | Absent | Absent | Absent | Absent | Absent | Absent | Absent |
| MAG217 | Present | Present | Absent | Absent | Absent | Absent | Absent | Present | Absent | Present | Present | Absent |
| MAG220 | Present | Absent | Present | Absent | Absent | Absent | Absent | Absent | Absent | Present | Absent | Present |
| MAG221 | Absent | Absent | Absent | Absent | Present | Absent | Absent | Absent | Absent | Absent | Absent | Absent |
| MAG222 | Absent | Absent | Absent | Absent | Absent | Absent | Absent | Absent | Absent | Absent | Absent | Absent |
| MAG223 | Absent | Absent | Absent | Absent | Absent | Absent | Absent | Absent | Absent | Absent | Absent | Absent |
| MAG225 | Absent | Absent | Absent | Absent | Absent | Absent | Absent | Absent | Absent | Absent | Absent | Absent |

| Function | Nitrate reduction | | | | Nitrite reduction to ammonia | | | Nitrite reduction | | Nitric oxide reduction | | |
|---|---|---|---|---|---|---|---|---|---|---|---|---|
| Gene | napA | napB | narG | narH | nrfA | nirB | nirD | nirK | nirS | norB | norC | nosZ |
| MAG230 | Absent | Absent | Absent | Absent | Present | Absent | Absent | Absent | Absent | Absent | Absent | Absent |
| MAG231 | Absent | Absent | Absent | Absent | Present | Absent | Absent | Absent | Absent | Absent | Absent | Absent |
| MAG232 | Absent | Absent | Absent | Absent | Absent | Absent | Absent | Absent | Absent | Absent | Absent | Absent |
| MAG237 | Absent | Absent | Absent | Absent | Present | Absent | Present | Absent | Absent | Absent | Absent | Present |
| MAG238 | Absent | Absent | Absent | Absent | Absent | Absent | Absent | Absent | Absent | Present | Present | Absent |
| MAG239 | Absent | Absent | Absent | Absent | Present | Absent | Absent | Absent | Absent | Absent | Absent | Absent |
| MAG240 | Absent | Absent | Present | Present | Absent | Absent | Absent | Absent | Absent | Present | Absent | Absent |
| MAG242 | Absent | Absent | Absent | Absent | Absent | Absent | Absent | Absent | Absent | Absent | Absent | Absent |
| MAG247 | Absent | Absent | Absent | Absent | Absent | Absent | Absent | Absent | Absent | Absent | Absent | Absent |
| MAG249 | Present | Present | Present | Present | Absent | Present | Present | Absent | Present | Present | Present | Absent |
| MAG252 | Present | Present | Present | Present | Absent | Present | Present | Absent | Present | Present | Present | Absent |
| MAG253 | Absent | Absent | Absent | Absent | Absent | Absent | Absent | Absent | Absent | Absent | Absent | Absent |
| MAG257 | Present | Present | Present | Present | Absent | Present | Present | Absent | Present | Present | Absent | Absent |
| MAG261 | Absent | Absent | Present | Present | Present | Present | Present | Absent | Present | Absent | Absent | Absent |
| MAG271 | Present | Absent | Absent | Absent | Present | Absent | Absent | Absent | Absent | Present | Present | Absent |
| MAG273 | Absent | Absent | Absent | Absent | Absent | Absent | Absent | Absent | Absent | Absent | Absent | Absent |
| MAG277 | Absent | Absent | Absent | Absent | Absent | Present | Present | Absent | Absent | Absent | Absent | Absent |
| MAG279 | Absent | Absent | Absent | Absent | Present | Absent | Absent | Absent | Absent | Present | Absent | Absent |
| MAG282 | Absent | Absent | Absent | Absent | Absent | Present | Present | Absent | Present | Absent | Absent | Absent |
| MAG284 | Absent | Absent | Absent | Absent | Absent | Absent | Absent | Absent | Absent | Absent | Absent | Absent |
| MAG285 | Absent | Absent | Absent | Absent | Absent | Present | Present | Absent | Absent | Absent | Absent | Absent |
| MAG286 | Absent | Absent | Absent | Absent | Present | Absent | Absent | Absent | Absent | Absent | Absent | Absent |
| MAG291 | Absent | Absent | Absent | Absent | Absent | Absent | Absent | Absent | Absent | Absent | Absent | Absent |
| MAG292 | Absent | Absent | Absent | Absent | Absent | Absent | Absent | Absent | Absent | Absent | Absent | Absent |
| MAG293 | Absent | Absent | Present | Present | Absent | Present | Present | Absent | Absent | Present | Present | Absent |
| MAG294 | Absent | Absent | Absent | Absent | Absent | Absent | Absent | Absent | Absent | Absent | Absent | Absent |
| MAG301 | Absent | Absent | Absent | Absent | Present | Absent | Absent | Absent | Absent | Absent | Absent | Absent |
| MAG303 | Absent | Absent | Absent | Absent | Absent | Absent | Absent | Absent | Absent | Absent | Absent | Absent |
| MAG306 | Absent | Absent | Absent | Absent | Absent | Present | Present | Absent | Absent | Absent | Absent | Absent |
| MAG307 | Absent | Absent | Present | Present | Absent | Present | Present | Absent | Present | Present | Absent | Absent |
| MAG309 | Present | Present | Present | Present | Present | Present | Present | Absent | Absent | Present | Present | Absent |
| MAG310 | Absent | Absent | Absent | Absent | Present | Absent | Absent | Present | Present | Absent | Absent | Absent |

| Function | Nitrate reduction | | | | Nitrite reduction to ammonia | | | Nitrite reduction | | Nitric oxide reduction | | |
|---|---|---|---|---|---|---|---|---|---|---|---|---|
| Gene | napA | napB | narG | narH | nrfA | nirB | nirD | nirK | nirS | norB | norC | nosZ |
| MAG311 | Absent | Absent | Absent | Absent | Absent | Absent | Absent | Absent | Absent | Absent | Absent | Absent |
| MAG312 | Absent | Absent | Absent | Absent | Present | Absent | Absent | Absent | Absent | Present | Absent | Absent |
| MAG318 | Absent | Absent | Absent | Absent | Absent | Absent | Absent | Absent | Absent | Present | Absent | Absent |
| MAG320 | Absent | Absent | Absent | Absent | Absent | Absent | Absent | Absent | Absent | Absent | Absent | Absent |
| MAG321 | Absent | Absent | Absent | Absent | Present | Absent | Absent | Absent | Absent | Absent | Absent | Absent |
| MAG325 | Present | Present | Absent | Absent | Absent | Present | Present | Absent | Present | Present | Present | Present |
| MAG328 | Absent | Absent | Absent | Absent | Present | Absent | Absent | Absent | Absent | Absent | Absent | Absent |
| MAG330 | Absent | Present | Absent | Absent | Absent | Absent | Absent | Absent | Present | Present | Present | Absent |

### Table S9. Presence/ absence of nitrification and nitrogen fixation.

| Function | Ammonia oxidation | | | Nitrite oxidation | | $N_2$ fixation | | | |
|---|---|---|---|---|---|---|---|---|---|
| Gene | amoA | amoB | amoC | nxrA | nxrB | nifD | nifK | vnfD | nifH |
| MAG001 | Absent | Absent | Absent | Absent | Absent | Absent | Absent | Absent | Absent |
| MAG002 | Absent | Absent | Absent | Absent | Absent | Absent | Absent | Absent | Present |
| MAG003 | Absent | Absent | Absent | Absent | Absent | Absent | Absent | Absent | Absent |
| MAG005 | Absent | Absent | Absent | Absent | Absent | Absent | Absent | Absent | Absent |
| MAG006 | Absent | Absent | Absent | Absent | Absent | Absent | Absent | Absent | Absent |
| MAG008 | Absent | Absent | Absent | Absent | Absent | Absent | Absent | Absent | Absent |
| MAG009 | Absent | Absent | Absent | Absent | Absent | Present | Present | Absent | Present |
| MAG011 | Absent | Absent | Absent | Absent | Absent | Absent | Absent | Absent | Absent |
| MAG012 | Absent | Absent | Absent | Absent | Absent | Present | Present | Absent | Present |
| MAG013 | Absent | Absent | Absent | Absent | Absent | Absent | Absent | Absent | Present |
| MAG014 | Absent | Absent | Absent | Absent | Absent | Absent | Absent | Absent | Absent |
| MAG018 | Absent | Absent | Absent | Absent | Absent | Absent | Absent | Absent | Absent |
| MAG019 | Absent | Absent | Absent | Absent | Absent | Absent | Absent | Absent | Absent |
| MAG020 | Absent | Absent | Absent | Absent | Absent | Absent | Absent | Absent | Absent |
| MAG025 | Absent | Absent | Absent | Present | Present | Present | Present | Absent | Present |
| MAG026 | Absent | Absent | Absent | Absent | Absent | Absent | Absent | Absent | Absent |
| MAG027 | Absent | Absent | Absent | Absent | Absent | Absent | Absent | Absent | Absent |
| MAG028 | Present | Present | Present | Absent | Absent | Absent | Absent | Absent | Absent |
| MAG030 | Absent | Absent | Absent | Absent | Absent | Absent | Absent | Absent | Absent |
| MAG037 | Absent | Absent | Absent | Absent | Absent | Absent | Absent | Absent | Absent |
| MAG038 | Absent | Absent | Absent | Absent | Absent | Present | Present | Absent | Present |
| MAG042 | Absent | Absent | Absent | Absent | Absent | Absent | Absent | Absent | Absent |
| MAG044 | Absent | Absent | Absent | Absent | Absent | Absent | Absent | Absent | Absent |
| MAG045 | Present | Present | Present | Absent | Absent | Absent | Absent | Absent | Absent |
| MAG046 | Absent | Absent | Absent | Absent | Absent | Present | Present | Absent | Present |
| MAG051 | Absent | Absent | Absent | Absent | Absent | Absent | Absent | Absent | Absent |
| MAG052 | Absent | Absent | Absent | Absent | Absent | Absent | Absent | Absent | Absent |
| MAG053 | Absent | Absent | Absent | Absent | Absent | Absent | Absent | Absent | Absent |

| Function | Ammonia oxidation | | | Nitrite oxidation | | $N_2$ fixation | | | |
|---|---|---|---|---|---|---|---|---|---|
| Gene | amoA | amoB | amoC | nxrA | nxrB | nifD | nifK | vnfD | nifH |
| MAG054 | Absent | Absent | Absent | Absent | Absent | Absent | Absent | Absent | Absent |
| MAG055 | Absent | Absent | Absent | Absent | Absent | Absent | Absent | Absent | Absent |
| MAG056 | Absent | Absent | Absent | Absent | Absent | Absent | Absent | Absent | Absent |
| MAG057 | Absent | Absent | Absent | Absent | Absent | Absent | Absent | Absent | Present |
| MAG059 | Absent | Absent | Absent | Absent | Absent | Absent | Absent | Absent | Absent |
| MAG062 | Absent | Absent | Absent | Absent | Absent | Absent | Absent | Absent | Absent |
| MAG064 | Absent | Absent | Absent | Absent | Absent | Absent | Absent | Absent | Absent |
| MAG066 | Absent | Absent | Absent | Absent | Absent | Absent | Absent | Absent | Absent |
| MAG067 | Absent | Absent | Absent | Present | Present | Absent | Absent | Absent | Absent |
| MAG069 | Absent | Absent | Absent | Absent | Absent | Absent | Absent | Absent | Absent |
| MAG071 | Absent | Absent | Absent | Absent | Absent | Absent | Absent | Absent | Absent |
| MAG072 | Absent | Absent | Absent | Absent | Absent | Absent | Absent | Absent | Absent |
| MAG073 | Absent | Absent | Absent | Absent | Absent | Absent | Absent | Absent | Absent |
| MAG074 | Absent | Absent | Absent | Absent | Absent | Absent | Absent | Absent | Absent |
| MAG079 | Absent | Absent | Absent | Absent | Absent | Absent | Absent | Absent | Absent |
| MAG080 | Absent | Absent | Absent | Absent | Absent | Absent | Absent | Absent | Absent |
| MAG081 | Absent | Absent | Absent | Absent | Absent | Absent | Absent | Absent | Present |
| MAG085 | Absent | Absent | Absent | Absent | Absent | Absent | Absent | Absent | Absent |
| MAG086 | Absent | Absent | Absent | Absent | Absent | Absent | Absent | Absent | Absent |
| MAG088 | Absent | Absent | Absent | Absent | Absent | Absent | Absent | Absent | Absent |
| MAG089 | Absent | Absent | Absent | Absent | Absent | Absent | Absent | Absent | Absent |
| MAG091 | Absent | Absent | Absent | Absent | Absent | Absent | Absent | Absent | Absent |
| MAG096 | Absent | Absent | Absent | Absent | Absent | Absent | Absent | Absent | Absent |
| MAG101 | Absent | Absent | Absent | Absent | Absent | Present | Present | Absent | Present |
| MAG102 | Absent | Absent | Absent | Absent | Absent | Absent | Absent | Absent | Absent |
| MAG104 | Absent | Absent | Absent | Absent | Absent | Absent | Absent | Absent | Absent |
| MAG106 | Absent | Absent | Absent | Absent | Absent | Absent | Absent | Absent | Absent |
| MAG107 | Absent | Absent | Absent | Absent | Absent | Present | Present | Absent | Present |
| MAG109 | Absent | Absent | Absent | Absent | Absent | Absent | Absent | Absent | Absent |
| MAG110 | Absent | Absent | Absent | Absent | Absent | Absent | Absent | Absent | Absent |
| MAG111 | Absent | Absent | Absent | Absent | Absent | Absent | Absent | Absent | Absent |
| MAG112 | Absent | Absent | Absent | Absent | Absent | Absent | Absent | Absent | Absent |
| MAG114 | Absent | Absent | Absent | Absent | Absent | Absent | Absent | Absent | Absent |
| MAG115 | Absent | Absent | Absent | Absent | Absent | Absent | Absent | Absent | Absent |
| MAG118 | Absent | Absent | Absent | Absent | Absent | Absent | Absent | Absent | Absent |
| MAG120 | Absent | Absent | Absent | Absent | Absent | Absent | Absent | Absent | Absent |
| MAG121 | Absent | Absent | Absent | Absent | Absent | Present | Present | Absent | Present |
| MAG124 | Absent | Absent | Absent | Absent | Absent | Absent | Absent | Absent | Absent |
| MAG127 | Absent | Absent | Absent | Present | Present | Absent | Absent | Absent | Absent |
| MAG132 | Absent | Absent | Absent | Absent | Absent | Absent | Absent | Absent | Absent |
| MAG133 | Absent | Absent | Absent | Absent | Absent | Absent | Absent | Absent | Absent |
| MAG134 | Absent | Absent | Absent | Absent | Absent | Absent | Absent | Absent | Absent |
| MAG137 | Absent | Absent | Absent | Absent | Absent | Absent | Absent | Absent | Absent |
| MAG138 | Absent | Absent | Absent | Absent | Absent | Absent | Absent | Absent | Absent |
| MAG142 | Absent | Absent | Absent | Absent | Absent | Absent | Absent | Absent | Absent |

| Function | Ammonia oxidation | | | Nitrite oxidation | | $N_2$ fixation | | | |
|---|---|---|---|---|---|---|---|---|---|
| Gene | amoA | amoB | amoC | nxrA | nxrB | nifD | nifK | vnfD | nifH |
| MAG144 | Absent | Absent | Absent | Present | Present | Absent | Absent | Absent | Absent |
| MAG149 | Absent | Absent | Absent | Present | Absent | Absent | Absent | Absent | Absent |
| MAG154 | Absent | Absent | Absent | Absent | Absent | Absent | Absent | Absent | Absent |
| MAG156 | Absent | Absent | Absent | Absent | Absent | Present | Absent | Absent | Present |
| MAG158 | Absent | Absent | Absent | Absent | Absent | Absent | Absent | Absent | Absent |
| MAG160 | Absent | Absent | Absent | Absent | Absent | Absent | Absent | Absent | Absent |
| MAG162 | Absent | Absent | Absent | Absent | Absent | Absent | Absent | Absent | Absent |
| MAG163 | Absent | Absent | Absent | Absent | Absent | Absent | Absent | Absent | Present |
| MAG164 | Absent | Absent | Absent | Absent | Absent | Absent | Absent | Absent | Absent |
| MAG166 | Absent | Absent | Absent | Absent | Absent | Absent | Absent | Absent | Absent |
| MAG170 | Absent | Absent | Absent | Absent | Absent | Absent | Absent | Absent | Absent |
| MAG174 | Absent | Absent | Absent | Absent | Absent | Absent | Absent | Absent | Present |
| MAG175 | Absent | Absent | Absent | Absent | Absent | Absent | Absent | Absent | Absent |
| MAG176 | Present | Absent | Absent | Present | Present | Absent | Absent | Absent | Absent |
| MAG181 | Absent | Absent | Absent | Absent | Absent | Absent | Absent | Absent | Absent |
| MAG184 | Absent | Absent | Absent | Absent | Absent | Absent | Absent | Absent | Absent |
| MAG187 | Absent | Absent | Absent | Absent | Absent | Absent | Absent | Absent | Absent |
| MAG188 | Absent | Absent | Absent | Absent | Absent | Present | Present | Absent | Present |
| MAG189 | Absent | Absent | Absent | Absent | Absent | Absent | Absent | Absent | Absent |
| MAG190 | Absent | Absent | Absent | Absent | Absent | Absent | Absent | Absent | Present |
| MAG191 | Absent | Absent | Absent | Absent | Absent | Present | Present | Absent | Present |
| MAG196 | Absent | Absent | Absent | Absent | Absent | Present | Present | Absent | Present |
| MAG197 | Present | Absent | Absent | Absent | Absent | Present | Present | Absent | Present |
| MAG198 | Absent | Absent | Absent | Absent | Absent | Present | Present | Absent | Present |
| MAG203 | Absent | Absent | Absent | Absent | Absent | Absent | Absent | Absent | Absent |
| MAG204 | Absent | Absent | Absent | Absent | Absent | Absent | Absent | Absent | Absent |
| MAG205 | Absent | Absent | Absent | Absent | Absent | Present | Present | Absent | Present |
| MAG206 | Absent | Absent | Absent | Absent | Absent | Absent | Absent | Absent | Absent |
| MAG207 | Absent | Absent | Absent | Absent | Absent | Absent | Absent | Absent | Absent |
| MAG217 | Absent | Absent | Absent | Absent | Absent | Absent | Absent | Absent | Absent |
| MAG220 | Absent | Absent | Absent | Absent | Absent | Present | Present | Absent | Present |
| MAG221 | Absent | Absent | Absent | Absent | Absent | Absent | Absent | Absent | Absent |
| MAG222 | Absent | Absent | Absent | Absent | Absent | Absent | Absent | Absent | Absent |
| MAG223 | Absent | Absent | Absent | Absent | Absent | Absent | Absent | Absent | Absent |
| MAG225 | Absent | Absent | Absent | Absent | Absent | Absent | Absent | Absent | Absent |
| MAG230 | Absent | Absent | Absent | Absent | Absent | Absent | Absent | Absent | Absent |
| MAG231 | Absent | Absent | Absent | Absent | Absent | Absent | Absent | Absent | Absent |
| MAG232 | Absent | Absent | Absent | Absent | Absent | Absent | Absent | Absent | Absent |
| MAG237 | Absent | Absent | Absent | Absent | Absent | Absent | Absent | Absent | Absent |
| MAG238 | Absent | Absent | Absent | Absent | Absent | Absent | Absent | Absent | Absent |
| MAG239 | Absent | Absent | Absent | Absent | Absent | Absent | Absent | Absent | Absent |
| MAG240 | Absent | Absent | Absent | Absent | Absent | Absent | Absent | Absent | Absent |
| MAG242 | Absent | Absent | Absent | Absent | Absent | Absent | Absent | Absent | Absent |
| MAG247 | Absent | Absent | Absent | Absent | Absent | Absent | Absent | Absent | Absent |
| MAG249 | Absent | Absent | Absent | Absent | Absent | Absent | Absent | Absent | Absent |

| Function | Ammonia oxidation | | | Nitrite oxidation | | $N_2$ fixation | | | |
|---|---|---|---|---|---|---|---|---|---|
| Gene | amoA | amoB | amoC | nxrA | nxrB | nifD | nifK | vnfD | nifH |
| MAG252 | Absent | Absent | Absent | Absent | Absent | Absent | Absent | Absent | Absent |
| MAG253 | Absent | Absent | Absent | Absent | Absent | Present | Present | Absent | Present |
| MAG257 | Absent | Absent | Absent | Absent | Absent | Present | Present | Absent | Present |
| MAG261 | Absent | Absent | Absent | Absent | Absent | Absent | Absent | Absent | Absent |
| MAG271 | Absent | Absent | Absent | Absent | Absent | Absent | Absent | Absent | Absent |
| MAG273 | Absent | Absent | Absent | Absent | Absent | Present | Present | Absent | Present |
| MAG277 | Absent | Absent | Absent | Absent | Absent | Absent | Absent | Absent | Absent |
| MAG279 | Absent | Absent | Absent | Absent | Absent | Present | Present | Absent | Present |
| MAG282 | Absent | Absent | Absent | Absent | Absent | Absent | Absent | Absent | Absent |
| MAG284 | Absent | Absent | Absent | Absent | Absent | Present | Present | Absent | Present |
| MAG285 | Absent | Absent | Absent | Absent | Absent | Present | Present | Absent | Present |
| MAG286 | Absent | Absent | Absent | Absent | Absent | Absent | Absent | Absent | Absent |
| MAG291 | Absent | Absent | Absent | Absent | Absent | Present | Present | Absent | Present |
| MAG292 | Absent | Absent | Absent | Absent | Absent | Absent | Absent | Absent | Present |
| MAG293 | Absent | Absent | Absent | Absent | Absent | Present | Present | Absent | Present |
| MAG294 | Absent | Present | Present | Absent | Absent | Absent | Absent | Absent | Absent |
| MAG301 | Absent | Absent | Absent | Absent | Absent | Absent | Absent | Absent | Absent |
| MAG303 | Absent | Absent | Absent | Absent | Absent | Absent | Absent | Absent | Absent |
| MAG306 | Absent | Absent | Absent | Absent | Absent | Absent | Absent | Absent | Absent |
| MAG307 | Absent | Absent | Absent | Absent | Absent | Absent | Absent | Absent | Absent |
| MAG309 | Absent | Absent | Absent | Absent | Absent | Present | Present | Absent | Present |
| MAG310 | Absent | Absent | Absent | Absent | Absent | Absent | Absent | Absent | Absent |
| MAG311 | Absent | Absent | Absent | Absent | Absent | Absent | Absent | Absent | Absent |
| MAG312 | Absent | Absent | Absent | Absent | Absent | Absent | Absent | Absent | Absent |
| MAG318 | Absent | Absent | Absent | Absent | Absent | Absent | Absent | Absent | Absent |
| MAG320 | Absent | Absent | Absent | Absent | Absent | Absent | Absent | Absent | Absent |
| MAG321 | Absent | Absent | Absent | Absent | Absent | Absent | Absent | Absent | Absent |
| MAG325 | Absent | Absent | Absent | Absent | Absent | Present | Present | Absent | Present |
| MAG328 | Absent | Absent | Absent | Absent | Absent | Absent | Absent | Absent | Present |
| MAG330 | Absent | Absent | Absent | Absent | Absent | Present | Present | Absent | Present |

### Table S10. Presence/absence of chlorite dismutation genes and cytochrome oxidases.

| Function | Chlorite reduction | Oxygen metabolism - cytochrome c oxidase, $cbb_3$-type | | |
|---|---|---|---|---|
| Gene | cld | ccoN | ccoO | ccoP |
| MAG001 | Absent | Present | Present | Present |
| MAG002 | Absent | Absent | Absent | Absent |
| MAG003 | Absent | Absent | Absent | Absent |
| MAG005 | Present | Absent | Absent | Absent |
| MAG006 | Absent | Present | Present | Present |
| MAG008 | Present | Absent | Absent | Absent |
| MAG009 | Absent | Present | Present | Present |
| MAG011 | Absent | Present | Present | Present |
| MAG012 | Absent | Absent | Absent | Absent |
| MAG013 | Absent | Absent | Absent | Absent |
| MAG014 | Absent | Absent | Absent | Absent |

| Function | Chlorite reduction | Oxygen metabolism - cytochrome c oxidase, $cbb_3$-type | | |
|---|---|---|---|---|
| Gene | cld | ccoN | ccoO | ccoP |
| MAG018 | Absent | Present | Present | Present |
| MAG019 | Absent | Present | Present | Present |
| MAG020 | Absent | Absent | Absent | Absent |
| MAG025 | Absent | Absent | Absent | Present |
| MAG026 | Absent | Absent | Absent | Absent |
| MAG027 | Absent | Absent | Absent | Absent |
| MAG028 | Present | Absent | Absent | Absent |
| MAG030 | Absent | Absent | Absent | Absent |
| MAG037 | Absent | Present | Present | Present |
| MAG038 | Absent | Present | Present | Present |
| MAG042 | Absent | Absent | Absent | Absent |
| MAG044 | Absent | Absent | Absent | Absent |
| MAG045 | Present | Absent | Absent | Absent |
| MAG046 | Absent | Present | Present | Present |
| MAG051 | Present | Absent | Absent | Absent |
| MAG052 | Absent | Absent | Absent | Present |
| MAG053 | Absent | Absent | Absent | Absent |
| MAG054 | Absent | Absent | Absent | Absent |
| MAG055 | Absent | Absent | Absent | Absent |
| MAG056 | Absent | Present | Present | Present |
| MAG057 | Absent | Absent | Absent | Absent |
| MAG059 | Absent | Absent | Absent | Absent |
| MAG062 | Absent | Present | Present | Present |
| MAG064 | Absent | Absent | Absent | Absent |
| MAG066 | Absent | Present | Present | Present |
| MAG067 | Absent | Absent | Absent | Absent |
| MAG069 | Absent | Absent | Absent | Absent |
| MAG071 | Present | Absent | Absent | Present |
| MAG072 | Absent | Absent | Absent | Absent |
| MAG073 | Absent | Absent | Absent | Absent |
| MAG074 | Absent | Absent | Absent | Absent |
| MAG079 | Absent | Present | Present | Present |
| MAG080 | Absent | Absent | Absent | Absent |
| MAG081 | Absent | Absent | Absent | Absent |
| MAG085 | Absent | Absent | Absent | Absent |
| MAG086 | Absent | Absent | Absent | Absent |
| MAG088 | Absent | Absent | Absent | Absent |
| MAG089 | Absent | Absent | Absent | Present |
| MAG091 | Absent | Absent | Absent | Absent |
| MAG096 | Absent | Present | Present | Present |
| MAG101 | Absent | Absent | Absent | Absent |
| MAG102 | Absent | Absent | Absent | Absent |
| MAG104 | Absent | Absent | Absent | Absent |
| MAG106 | Absent | Present | Present | Present |
| MAG107 | Absent | Absent | Absent | Absent |
| MAG109 | Absent | Present | Present | Present |
| MAG110 | Absent | Absent | Absent | Present |
| MAG111 | Absent | Present | Present | Present |
| MAG112 | Absent | Absent | Absent | Absent |
| MAG114 | Absent | Present | Present | Present |
| MAG115 | Absent | Absent | Absent | Absent |
| MAG118 | Present | Absent | Absent | Absent |
| MAG120 | Absent | Present | Present | Absent |
| MAG121 | Absent | Absent | Absent | Absent |
| MAG124 | Absent | Absent | Absent | Absent |

| Function | Chlorite reduction | Oxygen metabolism - cytochrome c oxidase, $cbb_3$-type | | |
|---|---|---|---|---|
| Gene | cld | ccoN | ccoO | ccoP |
| MAG127 | Absent | Present | Present | Present |
| MAG132 | Absent | Absent | Absent | Absent |
| MAG133 | Absent | Absent | Absent | Present |
| MAG134 | Absent | Absent | Absent | Absent |
| MAG137 | Absent | Absent | Absent | Absent |
| MAG138 | Absent | Absent | Absent | Absent |
| MAG142 | Absent | Absent | Absent | Absent |
| MAG144 | Absent | Present | Present | Present |
| MAG149 | Present | Present | Present | Absent |
| MAG154 | Absent | Present | Present | Present |
| MAG156 | Absent | Absent | Absent | Absent |
| MAG158 | Absent | Absent | Absent | Absent |
| MAG160 | Absent | Absent | Absent | Absent |
| MAG162 | Absent | Absent | Absent | Absent |
| MAG163 | Absent | Absent | Absent | Absent |
| MAG164 | Absent | Absent | Absent | Absent |
| MAG166 | Absent | Absent | Absent | Absent |
| MAG170 | Absent | Present | Present | Present |
| MAG174 | Absent | Absent | Absent | Absent |
| MAG175 | Present | Absent | Absent | Present |
| MAG176 | Present | Absent | Absent | Absent |
| MAG181 | Absent | Absent | Absent | Present |
| MAG184 | Absent | Absent | Absent | Absent |
| MAG187 | Absent | Absent | Absent | Absent |
| MAG188 | Absent | Absent | Absent | Absent |
| MAG189 | Absent | Absent | Absent | Absent |
| MAG190 | Present | Absent | Absent | Absent |
| MAG191 | Absent | Absent | Absent | Absent |
| MAG196 | Absent | Absent | Absent | Absent |
| MAG197 | Absent | Present | Present | Present |
| MAG198 | Absent | Present | Present | Present |
| MAG203 | Absent | Present | Present | Present |
| MAG204 | Absent | Present | Present | Present |
| MAG205 | Absent | Present | Present | Present |
| MAG206 | Absent | Present | Present | Present |
| MAG207 | Absent | Absent | Absent | Absent |
| MAG217 | Present | Absent | Absent | Present |
| MAG220 | Absent | Absent | Absent | Absent |
| MAG221 | Absent | Absent | Absent | Absent |
| MAG222 | Absent | Absent | Absent | Absent |
| MAG223 | Absent | Absent | Absent | Absent |
| MAG225 | Absent | Absent | Absent | Absent |
| MAG230 | Absent | Absent | Absent | Absent |
| MAG231 | Absent | Absent | Absent | Absent |
| MAG232 | Absent | Absent | Absent | Absent |
| MAG237 | Absent | Present | Present | Present |
| MAG238 | Absent | Absent | Absent | Absent |
| MAG239 | Present | Absent | Absent | Absent |
| MAG240 | Present | Present | Present | Present |
| MAG242 | Absent | Absent | Absent | Absent |
| MAG247 | Absent | Absent | Absent | Absent |
| MAG249 | Absent | Present | Present | Present |
| MAG252 | Absent | Present | Present | Present |
| MAG253 | Absent | Absent | Absent | Absent |
| MAG257 | Absent | Present | Present | Absent |

| Function | Chlorite reduction | Oxygen metabolism - cytochrome c oxidase, $cbb_3$-type | | |
|---|---|---|---|---|
| Gene | cld | ccoN | ccoO | ccoP |
| MAG261 | Absent | Present | Present | Present |
| MAG271 | Absent | Absent | Absent | Absent |
| MAG273 | Absent | Absent | Absent | Absent |
| MAG277 | Absent | Absent | Absent | Absent |
| MAG279 | Absent | Absent | Absent | Absent |
| MAG282 | Absent | Present | Present | Present |
| MAG284 | Absent | Present | Present | Present |
| MAG285 | Absent | Present | Present | Present |
| MAG286 | Absent | Absent | Absent | Absent |
| MAG291 | Absent | Present | Present | Present |
| MAG292 | Absent | Absent | Absent | Absent |
| MAG293 | Absent | Present | Present | Present |
| MAG294 | Present | Absent | Absent | Absent |
| MAG301 | Absent | Absent | Absent | Absent |
| MAG303 | Absent | Absent | Absent | Absent |
| MAG306 | Absent | Present | Present | Present |
| MAG307 | Absent | Present | Present | Present |
| MAG309 | Absent | Present | Present | Present |
| MAG310 | Present | Absent | Absent | Present |
| MAG311 | Absent | Absent | Absent | Absent |
| MAG312 | Absent | Absent | Absent | Absent |
| MAG318 | Absent | Absent | Absent | Absent |
| MAG320 | Absent | Absent | Absent | Absent |
| MAG321 | Absent | Absent | Absent | Absent |
| MAG325 | Absent | Present | Present | Present |
| MAG328 | Absent | Absent | Absent | Absent |
| MAG330 | Absent | Present | Absent | Absent |

7. McDonough LK, Santos IR, Andersen MS, O’Carroll DM, Rutlidge H, Meredith K, et al. Changes in global groundwater organic carbon driven by climate change and urbanization.

Nat Commun. 2020;11:1–10. http://dx.doi.org/10.1038/s41467-020-14946-1

8. Leroux SJ, Loreau M. Theoretical perspectives on bottom-up and top-down interactions across ecosystems. Trophic Ecol Bottom-Up Top-Down Interact Across Aquat Terr Syst. 2015;3–28.

9. Karwautz C, Zhou Y, Kerros M-E, Weinbauer MG, Griebler C. Bottom-Up Control of the Groundwater Microbial Food-Web in an Alpine Aquifer. Front Ecol Evol. 2022;10:1–14.

10. Hofmann R, Griebler C. DOM and bacterial growth efficiency in oligotrophic groundwater: Absence of priming and co-limitation by organic carbon and phosphorus. Mol Plant-Microbe Interact. 2018;31:311–22.

11. Hofmann R, Uhl J, Hertkorn N, Griebler C. Linkage Between Dissolved Organic Matter Transformation, Bacterial Carbon Production, and Diversity in a Shallow Oligotrophic Aquifer: Results From Flow-Through Sediment Microcosm Experiments. Front Microbiol. 2020;11.

12. Mehrshad M, Lopez-Fernandez M, Sundh J, Bell E, Simone D, Buck M, et al. Energy efficiency and biological interactions define the core microbiome of deep oligotrophic groundwater. Nat Commun. 2021;12:1–12.

13. Hahn CR, Farag IF, Murphy CL, Podar M, Elshahed MS, Youssef NH. Microbial Diversity and Sulfur Cycling in an Early Earth Analogue: From Ancient Novelty to Modern Commonality. MBio. 2022;13.

14. Colman DR, Kraus EA, Thieringer PH, Rempfert K, Templeton AS, Spear JR, et al. Deep-branching acetogens in serpentinized subsurface fluids of Oman. Proc Natl Acad Sci U S A. 2022;119:1–11.

15. Berlendis S, Ranchou-Peyruse M, Fardeau ML, Lascourrèges JF, Joseph M, Ollivier B, et al. *Desulfotomaculum aquiferis* sp. Nov. and *Desulfotomaculum profundi* sp. Nov., isolated from a deep natural gas storage aquifer. Int J Syst Evol Microbiol. 2016;66:4329–38.

16. Itävaara M, Nyyssönen M, Kapanen A, Nousiainen A, Ahonen L, Kukkonen I. Characterization of bacterial diversity to a depth of 1500m in the Outokumpu deep borehole, Fennoscandian Shield. FEMS Microbiol Ecol. 2011;77:295–309.

17. Sousa DZ, Visser M, Van Gelder AH, Boeren S, Pieterse MM, Pinkse MWH, et al. The deep-subsurface sulfate reducer *Desulfotomaculum kuznetsovii* employs two methanol-degrading pathways. Nat Commun. 2018;9.
